# FGF-dependent, polarized SOS activity orchestrates directed migration of *C. elegans* muscle progenitors independently of canonical effectors *in vivo*

**DOI:** 10.1101/2025.04.11.648432

**Authors:** Theresa V Gibney, Laila Latifi, Jacob I Mardick, Neal R Rasmussen, Maria C Lyons, Meera V Sundaram, David J Reiner, Ariel M Pani

**Affiliations:** Department of Biology, University of Virginia, Charlottesville, VA 22903, USA; Centre for Immuno-Oncology, Nuffield Department of Medicine, University of Oxford, Oxford OX3 7DQ, UK; Institute of Biosciences and Technology, Texas A&M Health Science Center, Texas A&M University, Houston, TX 77030, USA; Department of Genetics, Perelman School of Medicine, University of Pennsylvania, Philadelphia, PA 19104, USA; Department of Translational Medical Science, College of Medicine, Texas A&M Health Science Center, Texas A&M University, Houston, TX 77030, USA; Department of Cell Biology, University of Virginia School of Medicine, Charlottesville, VA 22903, USA

## Abstract

Directed cell migration is essential for animal development, tissue maintenance, regeneration, and disease states. Cells often migrate towards, or away from, sources of secreted signaling proteins that impart spatial information. How migrating cells interpret extracellular signals to orient and navigate within living animals is a fundamental question in biology. Receptor Tyrosine Kinase (RTK) signaling plays critical roles in cell migration, and aberrant RTK pathway activity is a key driver of multiple types of cancers. Yet, how RTKs control cell migration in living animals remains unclear, in part due to essential, pleiotropic roles of key proteins in development. To elucidate how RTK signaling controls cell migration *in vivo*, we dissected spatial and temporal requirements for key signal transduction and cytoskeletal regulatory proteins using *C. elegans* muscle progenitor migration as a tractable model. Cell type-specific depletion of endogenously tagged proteins revealed that homologs of FGFR, GRB2, SOS, and Ras control cell migration independently of their canonical ERK, PI3K, Akt, mTOR, and PLCγ effectors. Instead, we found that FGF-dependent, polarized SOS-1 orients migrating cells towards an FGF source, and mislocalizing SOS activity within migrating cells severely disrupts migration independent of ERK. Cell type-specific, gain-of-function experiments demonstrated that activated Ras is largely permissive for anterior migration in this context, and an intragenic revertant identified in a screen for suppressors of activated *Ras/let-60* revealed that signal transduction in migrating muscle progenitors can be genetically uncoupled from Ras-ERK-dependent developmental processes. We found that conserved regulators of branched actin assembly control SM protrusive dynamics but are not essential for accurate, FGF-directed migration. Our findings provide a novel mechanism for RTK-directed cell migration *in vivo* and highlight the importance of cell type-specific approaches to elucidate signal transduction mechanisms in physiologically relevant contexts. Our work also outlines a comprehensive framework for investigating RTK-dependent processes in a multicellular organism and introduces a versatile genetic toolkit for dissecting spatial and temporal signaling dynamics fundamental to development, homeostasis, and disease.

## Introduction

Directed cell migration is a fundamental process in animal development and allows integrated organ systems to be assembled from disparate cell lineages born from distant progenitors^1,2^. Migrating cells often rely on positional information from extracellular ligands to precisely navigate through the complex and dynamic environment of a developing animal to reach their destinations^1–3^. To do so, cells must translate extracellular cues that impart spatial information about their surroundings into coordinated cytoskeletal rearrangements that drive directional cell motility^2,4^. Receptor Tyrosine Kinases (RTKs) are a prominent, evolutionarily conserved class of transmembrane receptors that bind growth factors and other extracellular signaling molecules^5^. Signaling through RTKs is essential for developmental and homeostatic processes including cell migration, proliferation, differentiation, and survival, and abnormal RTK pathway activation is implicated in cancer and congenital disorders^5–7^. Canonical RTK activation leads to recruitment of cytosolic adaptor and effector proteins such as Growth factor Receptor-Bound protein 2 (GRB2), which recruits Son-of-Sevenless (SOS), which in turn activates small GTPases including Ras and Rac and their associated signal transduction pathways^5,8^. RTK signal transduction can involve multiple downstream effectors, including Extracellular signal Regulated Kinase/Mitogen-Activated Protein Kinase (ERK/MAPK), Phosphoinositide 3-Kinase (PI3K), Akt/Protein Kinase B, mechanistic Target of Rapamycin Complex 1 (mTORC1), and Phospholipase C-gamma (PLCγ), which allows diverse functions for individual receptors in different contexts^5,9,10^.

Fibroblast Growth Factors (FGFs) are a class of secreted signaling proteins that play critical roles in animal development and tissue maintenance, can disperse between cells over long distances, and can convey spatial information^11–13^. At the cellular level, FGFs act through their RTK FGF Receptors (FGFRs) to regulate cell behaviors such as migration along with other processes including proliferation and patterning^14–16^. While FGF signaling is required for cell migration in multiple contexts during animal development^17–20^, methodological challenges and pleiotropy have hampered dissecting the signal transduction mechanisms by which FGFs guide migrating cells *in vivo*. In particular, essential roles for key proteins in other aspects of development or physiology have confounded efforts to investigate their roles in cell migration using genetic methods. Consequently, our current understanding of the cell biological mechanisms by which extracellular cues such as FGFs direct cell migration comes largely from experiments in unicellular organisms and cultured cell systems, which are experimentally tractable but devoid of the developmental and organismal contexts that govern cell migration *in vivo*^21,22^. Investigations of how RTK and/or Ras signaling regulate migration in cultured cells and unicellular models for cell motility have revealed roles for Ras, MAPK, and PI3K activity in regulating the activities of cytoskeletal remodeling proteins that can organize local actin polymerization in response to RTK signaling^4,23^. FGFs may also control cell migration through transcriptional mechanisms rather than by directly influencing cell behavior^24–26^. Despite conserved roles for FGF and RTK signaling in cell migration and their importance in disease states, we lack a clear understanding of the signal transduction processes that translate positional information encoded by extracellular FGF into directional cell migration in living animals. Dissecting these mechanisms will require the ability to visualize migrating cell dynamics and to manipulate key endogenous proteins *in vivo* with spatial and temporal precision to bypass pleiotropic effects.

*C. elegans* postembryonic muscle progenitors provide an experimentally tractable framework to dissect cell signaling mechanisms in a living animal. During early larval development, two muscle progenitors, the sex myoblasts (SMs), migrate anteriorly from their birthplace near the tail and stop over the dorsal uterine cells located at the center of the developing gonad. There, SMs proliferate and differentiate into vulval and uterine muscle cells essential for egg-laying^27,28^. The FGF8/17/18 family homolog EGL-17 is a diffusible chemoattractant that acts instructively to guide migrating SMs^29–31^. FGF/EGL-17 is secreted by ventral midline blast cells during early SM migration, followed by high expression in the primary (1°) vulval precursor cell (VPC), and uterine cells^31^. Loss of *FGF/egl-17* or the 5a isoform of *FGFR/egl-15* results in migration failure and posterior displacement of SMs^28,32^. FGF or FGFR loss of function does not affect SM proliferation or cell fates at this stage, thereby making these cells a powerful system to separate mechanisms of signal transduction that direct cell migration from those involved in other FGF-dependent processes. Prior work has explored the role of canonical RTK-Ras-ERK signaling in SM migration^9,33^, but mutant analyses have been constrained by lethality and pleiotropic effects of key genes in this pathway, such as roles for Ras-ERK signaling in fate specification of FGF-expressing cells that attract migrating SMs^34–36^. Current data indicate the Ras homolog *let-60* is required for SM migration, but may act permissively, while roles for other signal transduction pathways that act downstream of RTKs including PI3K-Akt-mTOR and PLCγ have not been reported^1,9,33^. Likewise, the signal transduction mechanisms that are responsible for orienting migrating SMs towards an extracellular FGF source are unknown.

To address these fundamental questions about how spatial information provided by FGF is translated into directed cell migration towards the FGF source, we systematically tested functions for candidate proteins with potential roles in RTK-mediated cell polarization and motility *in vivo*. We used temporal and cell type-specific depletion of tagged endogenous proteins to interrogate spatial and temporal roles of seventeen core signal transduction and cytoskeletal regulatory proteins with plausible roles in FGF-directed cell migration. Surprisingly, we found that canonical RTK effectors including homologs of ERK, PI3K, Akt, mTOR, and PLCγ are dispensable cell-autonomously for SM migration. We identified polarized SOS-1 activity as a key link between extracellular FGF and directed cell migratory behaviors independent of its known functions in activating ERK or Rac. In migrating SMs, tagged endogenous SOS-1 localized to the anterior cell membrane in an FGFR-dependent manner and is polarized towards cells that express FGF/EGL-17. Uniformly activating SOS around the cell membrane disrupted FGF-directed cell migration through an ERK-independent mechanism and led to cell migration defects resembling loss of SOS-1 function. While polarized SOS activity was required to orient migrating SMs, activated alleles of Ras/LET-60 were genetically permissive for anterior migration in this context. Moreover, an intragenic revertant allele of an activating mutation in *Ras/let-60* genetically separated the function of *Ras/let-60* in SM migration from its ERK-dependent roles in larval viability and VPC cell fate specification. These findings offer a mechanism for how FGF signaling guides cell migration in a developing animal and provide a novel toolkit to dissect RTK signal transduction and cytoskeletal regulation *in vivo*.

## RESULTS

### An FGFR-GRB2-SOS-Ras module regulates SM migration

To test functions for candidate proteins in SM migration, we used Cas9-triggered homologous recombination to tag endogenous proteins with an auxin-inducible degron (AID) that targets AID-tagged proteins for degradation in the presence of an exogenous TIR1 adaptor protein and the small molecule auxin or its analog K-NAA^37,38^ (Fig 1A). To deplete proteins of interest in migrating SMs, we expressed TIR1 using an *hlh-8/twist* promoter^39^ that drives strong expression in the post-embryonic M lineage before, during, and after SM migration^40^. Using this approach, we tagged and depleted endogenous proteins hypothesized to act in FGF signal transduction or to be required for FGF-mediated migration.

**Figure 1.**
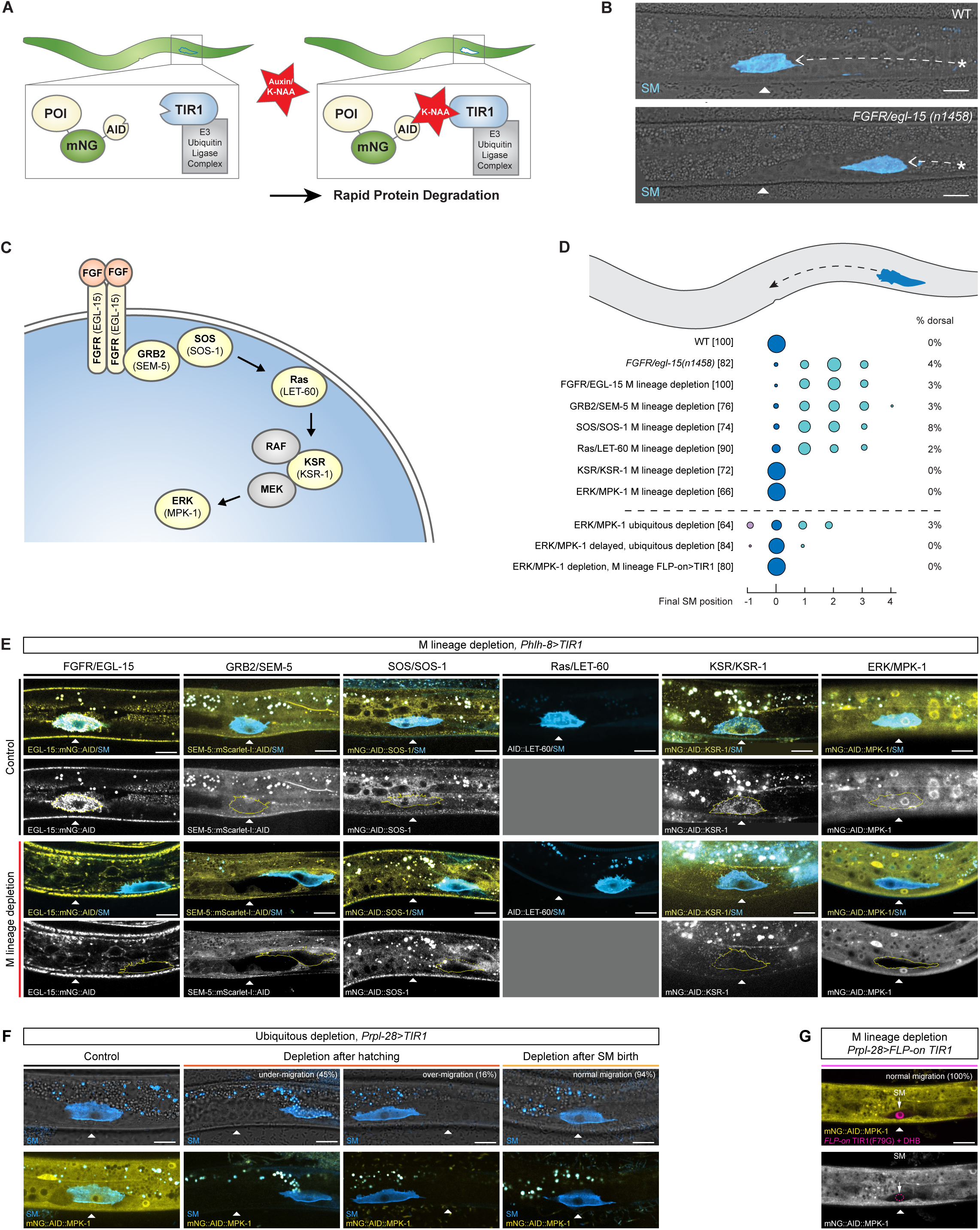
FGFR, GRB2, SOS, and Ras homologs are cell-autonomously required for SM migration independent of KSR or ERK. (**A**) Schematic of the TIR1-AID system for conditional protein depletion in *C. elegans.* AID-tagged proteins of interest (POI) are targeted for proteasomal degradation in the presence of TIR and auxin, or its synthetic analog K-NAA. (**B**) Representative images of SM positioning in wild-type and *FGFR/egl-15(n1458)* mutant animals. White triangles indicate the normal endpoint of migration over the P6.p cell. Dashed arrow indicates the approximate extent of migration. (**C**) Schematic diagram of the FGFR-GRB2-SOS-Ras-ERK signaling module. *C. elegans* homologs are indicated in parentheses, and proteins investigated here are shown as yellow. (**D**) Dot plot summarizing SM migration endpoints after depletion of endogenously tagged proteins. Area of each dot is proportional to the frequency of cells stopping at that position. Numbers of SMs analyzed for each strain are shown in brackets. Source data available in Table S1. (**E**) Representative, live spinning disk confocal images showing SM positioning and localization of endogenously tagged proteins in control and K-NAA-treated animals expressing TIR1 in the M lineage using an *hlh-8* promoter. The AID::Ras/LET-60 tag does not include a fluorescent protein. SMs are outlined in yellow in single channel images of tagged endogenous proteins. (**F**) Ubiquitous mNG::AID::ERK/MPK-1 depletion at two timepoints using *Prpl-28>TIR1* indicates that ERK signaling prior to SM birth is required for normal SM migration, but ERK activity during SM migration is dispensable. (**G**) Continuous ERK/MPK-1 depletion in the M lineage using *Phlh-8>FLP-D5; Prpl-28>FLP-on TIR1(F79G)* confirms that ERK activity in the M lineage is not required for correct SM positioning. The SM nucleus is marked by 2A::2x mKate2::DHB (magenta), which is co-expressed with TIR1(F79G). In all images, white triangles denote the normal endpoint of SM migration. The small, circular objects visible in both fluorescence channels are autofluorescent gut granules. All images are oriented with anterior to left and dorsal to top. Scale bars = 10µm.

We first investigated functions for six core components of the FGFR-Ras-ERK pathway (Fig. 1C), which were previously implicated in SM migration based on mutant analyses^33,41–43^.

Mutations in the 5a isoform of *egl-15,* the sole *C. elegans* FGF receptor, severely disrupt SM migration^44^, which also requires the FGF ligand EGL-17^29,32^ (Fig. 1B)*. Ras/let-60* was previously implicated as a permissive signal for FGF-dependent SM migration using mosaic rescue analyses^33^ but attempts to elucidate the mechanisms that orient SM migration towards an FGF source have been complicated by pleiotropic effects of key candidate genes. To determine how cells translate an extracellular FGF signal into directed migratory behaviors, we endogenously tagged homologs of FGF Receptor (FGFR/EGL-15), Growth factor Receptor-Bound protein 2 (GRB2/SEM-5), Son of Sevenless (SOS/SOS-1), Ras (Ras/LET-60), Kinase Suppressor of Ras (KSR/KSR-1), and Extracellular signal Regulated Kinase 1/2 (ERK/MPK-1) to visualize their localization and allow for cell type-specific protein depletion. FGFR/EGL-15::mNG::AID, GRB2/SEM-5::mScarlet-I::AID, mNG::AID::SOS-1, mNG::AID::KSR-1, and mNG::AID::ERK/MPK-1 knock-in animals appeared wild-type in the absence of the auxin analog K-NAA. We observed larval lethality for mScarlet-I::AID::LET-60 similar to that previously observed with an mNG::LET-60 tag^45^. However, we successfully generated separate, biologically functional mScarlet-I::LET-60 and AID::LET-60 knock-ins using an extended 30 amino acid flexible linker.

To assess functions for these proteins in SM migration, we depleted AID-tagged endogenous proteins using a *Phlh-8-*driven transgene that expresses TIR and a plasma membrane marker in M lineage cells. We treated with K-NAA or vehicle (water) and scored SM position at the early L3 stage based on proximity to the central P6.p-P8.p VPCs as well as the M-lineage-derived body wall muscles (Fig. S1). To validate our cell type-specific depletion approach, we first depleted FGFR/EGL-15, which led to strong SM migration defects (Fig. 1D, E) that phenocopied *egl-15(n1458),* which is a presumptive null mutation in the *egl-15(5a)* isoform required for SM migration^41^. Depleting the RTK adaptor protein GRB2/SEM-5 in SMs using *Phlh-8*-driven TIR1 resulted in migration defects similar to FGFR depletion, consistent with a requirement for GRB2/SEM-5 in FGFR/EGL-15 signal transduction^41,46^. Depleting the guanine nucleotide exchange factor SOS-1 or its small GTPase substrate Ras/LET-60 led to similar, although slightly less penetrant, defects in SM migration (Fig. 1D, E) consistent with essential functions for these proteins downstream of activated FGFR^9^.

Strikingly, we did not observe SM migration defects upon depletion of either KSR-1 or ERK/MPK-1 in the M lineage (Fig. 1D, E), suggesting these proteins are not cell-autonomously essential for FGF signal transduction during SM migration. While ERK/MPK-1-depleted SMs migrated to the correct position, we observed unusually long and thin cells in some animals (Fig. S2), which suggests that ERK signaling performs cell autonomous roles in SMs other than migration. To rule out the possibility of insufficient KSR-1 degradation, we examined SM positioning in animals carrying a *ksr-1(n2526)* mutation, which introduces a premature stop codon that truncates KSR-1 before its catalytic domain. While two missense alleles of *ksr-1* cause SM migration defects^33^, we observed normal SM positioning in *ksr-1(n2526)* null mutants, which corroborated the results of our cell type-specific KSR-1 depletion (Fig. S3). Although *ksr-1* sometimes acts redundantly with *ksr-2*, L2 stage scRNAseq data^47^ indicate that *ksr-2* is not normally expressed in SMs.

ERK/MPK-1 is an essential gene that has pleiotropic roles in specifying the fates of FGF/EGL-17-producing cells that attract migrating SMs^36,48,49^, which has complicated testing its functions in SM migration using mutants. We first validated that depleting our mNG::AID::MPK-1 allele was capable of imposing strong loss of function using a ubiquitously expressed *Prpl-28>TIR1::P2A::mCherry::his-11* transgene to deplete ERK/MPK-1 in all cell types. Plate-level observations showed whole-body mNG::AID::MPK-1 depletion phenocopied the fully penetrant adult sterility and larval lethality of the strongest *mpk-1* mutants^36^. To reconcile our cell type-specific depletion results with prior analyses that showed SM migration defects in *mpk-1* mutants^33^, we examined SM positioning in animals where ERK/MPK-1 was ubiquitously depleted after hatching. Similar to mutants^33^, we observed frequent SM migration defects (39/64 cells) in these animals, which included both slight overmigration (10/64 cells) and more severe undermigration (29/64 cells; Fig. 1D, F). To determine the timing of ERK activity involved in SM migration, we delayed ERK/MPK-1 depletion until after the SMs were born, but before the onset of FGF-dependent migration^31^. Ubiquitous ERK/MPK-1 depletion after SM birth caused only minor migration defects (5/84 cells; Fig. 1D, F), which may be attributed to slight asynchronies in larval development that caused earlier depletion in a subset of animals. To ensure that cell-autonomous and ubiquitous ERK/MPK-1 depletions were performed using the same level of TIR1 expression, we then depleted ERK/MPK-1 using a *Phlh-8>flp-D5; Prpl-28>FLP-on TIR1(F79G)::T2A::2x mKate2::DHB*^50^ strain in which TIR1 is expressed in the M-lineage using the same *rpl-28* promoter as for ubiquitous expression. We did not observe SM migration defects in any animals after M-lineage specific ERK/MPK-1 depletion using *Prpl-28>FLP-on>TIR1* (0/80 cells; Fig. 1D, G). These results indicate that the essential role for ERK signaling in SM migration occurs outside of the M lineage and before the SMs are specified, which is consistent with known roles for *mpk-1* in fate specification of *FGF/egl-17-*expressing VPCs^36^. Taken together, our cell type-specific depletion experiments demonstrated that a conserved FGFR-GRB2-SOS-Ras signaling module regulates SM migration cell-autonomously and independently of ERK/MPK-1, which acts downstream of RTK-Ras signaling in many other contexts^9^.

### SM migration does not require PI3K, Akt, mTOR, or PLCγ activity

As ERK/MPK-1 was not required in the M lineage for SM migration, we next tested functions for the PI3K-Akt pathway (Fig. 2A). PI3K-Akt signaling can mediate ERK-independent signal transduction downstream of Ras^10,23^, and is required for chemoattractant-driven migration in *Dictyostelium*^4,22^. To target PI3K-Akt signaling, we generated functional, mNG::AID-tagged alleles of the PI3K catalytic subunit AGE-1, the homolog of its downstream signaling effector Akt/PKB/AKT-1, and the mTOR homolog LET-363 (Fig. 2A). Because *C. elegans* encodes two *akt* homologs that typically act redundantly^51^, we generated AKT-1::mNG::AID in the viable *akt-2(ok393)* null mutant background. Depletion of PI3K/AGE-1 or AKT activity in the SMs had no effect on SM migration (Fig. 2B,C). To validate effective depletion via our AID-tagged alleles, we depleted the tagged endogenous proteins throughout the body using ubiquitously expressed TIR1. Strong *age-1* or *akt-1; akt-2* null mutants exhibit a maternal effect, constitutive dauer phenotype in the F2 generation^51^. Whole body depletion of AGE-1::mNG::AID or AKT-1::mNG::AID; *akt-2(ok393)* led to 100% F2 dauer progeny, which phenocopied putative null phenotypes and validated strong loss of function using our methodology.

**Figure 2.**
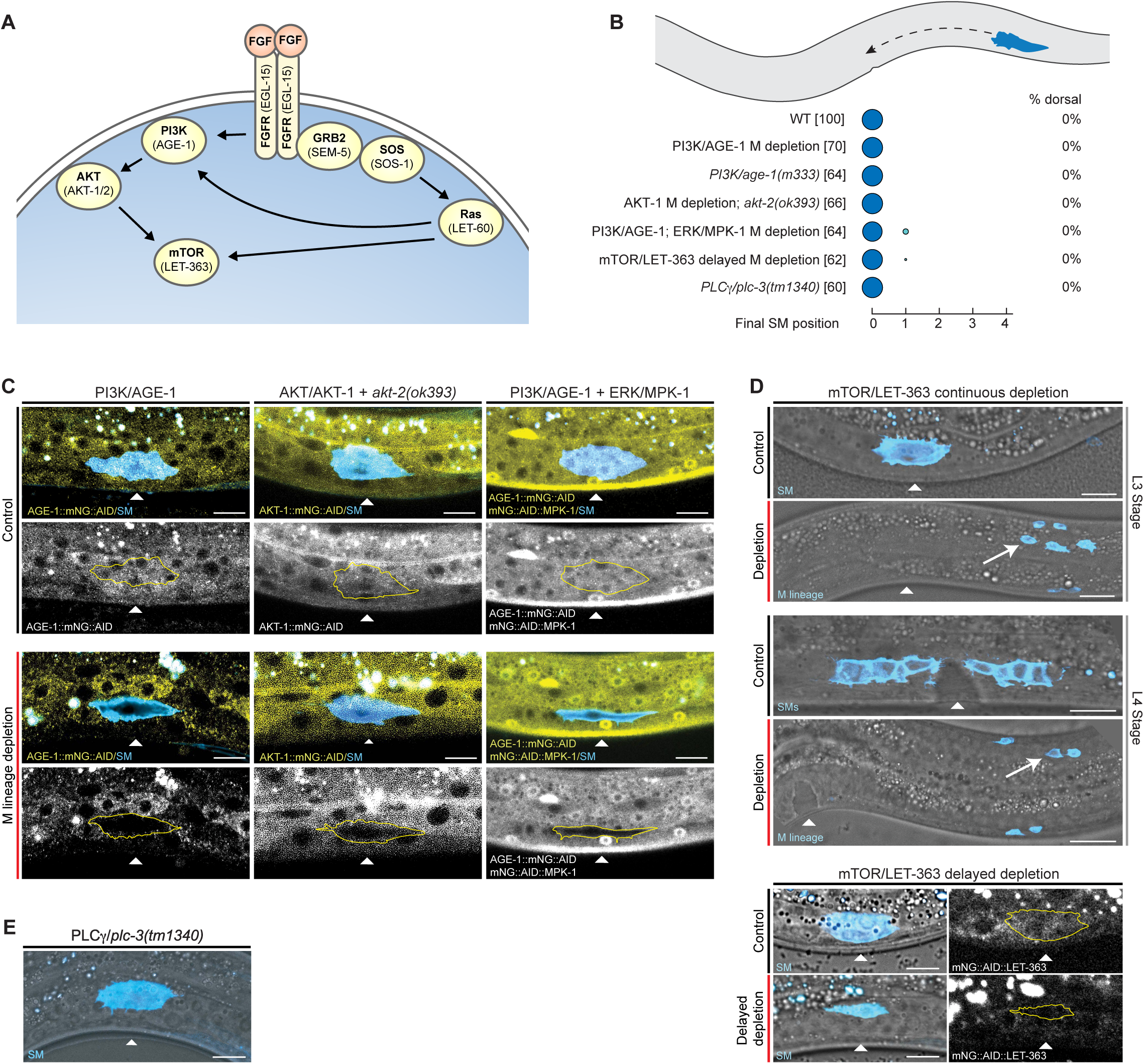
PI3K, AKT, mTOR, and PLCγ homologs are not required cell-autonomously for SM migration. **(A)** Schematic diagram of the conserved PI3K-Akt-mTOR signaling module along with components of the FGFR-GRB2-SOS-Ras module found to be required for SM migration. *C. elegans* homologs are indicated in parentheses. **(B)** Dot plot summarizing SM migration endpoints after M-lineage specific depletion of endogenously tagged proteins or in mutant strains. Area of each dot is proportional to the frequency of cells stopping at that position. Numbers of SMs analyzed for each strain are shown in brackets. Source data available in Table S1 (**C**) Representative, live spinning disk confocal images showing SM positioning and localization of endogenously tagged proteins of interest in control and K-NAA-treated animals. SMs are outlined in yellow in single channel images of tagged endogenous proteins. The small, circular objects visible in both fluorescence channels are autofluorescent gut granules. **(D)** Representative, live spinning disk confocal image showing control vs. K-NAA-treated mNG::AID::mTOR/LET-363 animals. Upper panel shows M lineage development after continuous K-NAA treatment in L3 stage animals when the SMs have completed migrating but have not started to proliferate compared to L4 stage animals at a time when the SMs have normally finished proliferating. Lower panel shows control and K-NAA-treated mNG::AID::mTOR/LET-363 worms with timed protein depletion. K-NAA was added after the SMs are born, but several hours prior to migration to circumvent early M lineage defects. Background subtraction^31^ was used to remove gut granules to better visualize the abnormally small M lineage cells. (**E**) SM positioning in a *PLCγ/plc-3(tm1340)* mutant animal. In all images, white triangles denote the normal endpoint of SM migration, and animals are oriented with anterior to left and dorsal to top. Scale bars = 10µm.

Given the importance of PI3K-Akt signaling in cell migration in other contexts, we sought to independently confirm that PI3K signaling is not required for SM migration. F1 generation *age-1(m333)* homozygotes from heterozygous mothers are reported to be egg-laying deficient^52^, which is a classic phenotypic outcome of SM migration defects^29^. Since the dauer phenotype of *age-1(m333)* null mutants occurs in the F2 generation, we examined SM positioning in F1 progeny of balanced *age-1(m333)/*mnC1 animals, which are expected to be 1/3 homozygous and 2/3 heterozygous for the *age-1(m333)* mutation (mnC1 homozygotes are Dpy Unc). Interestingly, we observed an SM differentiation defect that included ectopic SMs in 5/69 animals (Fig. S3). However, we observed no SM migration defects in animals with normal M lineage cell fates (0/64). These findings confirmed PI3K signaling is not required for SM migration while *age-1* has earlier functions in M lineage differentiation, likely through cell non-autonomous activity. To test the possibility that ERK and PI3K act redundantly in SM migration, we simultaneously depleted AGE-1::mNG::AID and mNG::AID::MPK-1 in the M lineage. SMs migrated to the correct position in the majority of animals, although we observed slight displacements in 9/64 cells (Fig. 2B,C). We also observed thinner SMs in AGE-1; MPK-1 depleted animals (Fig. 2C), which may have contributed to mispositioning in a small number of animals. We next evaluated the consequences of depleting the mTOR homolog LET-363, a protein that acts downstream of PI3K-Akt and promotes cell migration in contexts including cancer metastasis and T cell migration^53^. M lineage-specific depletion of mNG::AID::mTOR/LET-363 led to a striking phenotype where M lineage cells were entirely absent in many animals at the L3 stage. In animals where M lineage cells were visible, they were drastically smaller than normal, and SMs failed to migrate (Fig. 2D). To test the extent to which mTOR/LET-363 directly regulates SM migration, we delayed K-NAA addition until after the SMs were born, but several hours before the onset of FGF-dependent migration^31^. We did not observe migration defects with timed mTOR/LET-363 depletion, although SMs were still qualitatively smaller than control animals (Fig, 2D). Overall, our results suggest that PI3K, AKT, and mTOR signaling are not cell-autonomously required for SM migration, and that the PI3K-Akt and ERK pathways do not act redundantly in this process.

FGFs can also regulate cell migration through activation of Phospholipase-C gamma (PLCγ), which can in turn signal through Ca^2+^, diacylglycerol, and Protein Kinase C (PKC) family proteins^54–56^ and can regulate chemotaxis^57^. The single *C. elegans* PLCγ homolog, PLC-3, acts downstream of EGFR/LET-23 in gonadal sheath cells and the ALA neuron^58,59^, indicating that *C. elegans* RTKs are similarly capable of signaling through PLCγ. PLC-3 null mutants are sterile and thus *plc-3* would not have been identified in genetic screens for egg-laying deficient animals. To test functions of PLCγ in SM migration, we imaged SM positioning in homozygous *plc-3(tm1340)* offspring of heterozygous *plc-3(tm1340)/*+ parents. SM positioning was unaltered in *plc-3* null animals (Fig. 2E), indicating this pathway does not play an essential role in SM migration.

### Extracellular FGF/EGL-17 orients SOS-1 localization in migrating cells

To understand how migrating SMs navigate towards an FGF source, we next sought to determine the extent to which GRB2/SEM-5, SOS/SOS-1, and Ras/LET-60 act instructively to orient migrating cells versus permissively to allow migration. We first examined subcellular localization of the endogenously tagged proteins during the FGF-dependent phase of SM migration^28,31^. SOS and GRB2 homologs are cytosolic proteins that are recruited to the cell membrane by activated growth factor receptors, whereas Ras proteins are membrane-anchored through prenyl lipid modification of a C-terminal CAAX domain^60,61^. Strikingly, we found that mNG::AID::SOS-1 punctae localized to the cell membrane along the anterior and ventral edges of migrating SMs (Fig. 3A, B), which face the somatic gonad and ventral midline cells that express FGF/EGL-17 at this stage^31^. Time-lapse imaging revealed that mNG::AID::SOS-1 localized to dynamic punctae within active leading-edge protrusions in migrating SMs (Fig. 3C; Video 1), suggesting these locations are sites of receptor activation. GRB2/SEM-5::mScarlet-I::AID localized to sparse punctae at the cell membrane and faintly in the cytoplasm of migrating cells (Fig. 3A), but the protein was expressed at low levels and photobleached rapidly. Ras/mScarlet-I::LET-60 localized primarily to the plasma membrane without obvious anteroposterior polarization and was also visible in larger intracellular punctae (Fig. 3A). To assess the possibility that SOS-1 translates the direction of an FGF source into a polarized intracellular signal, we first tested the extent to which SOS-1 localization in SMs responds to FGF signaling. Polarized localization of mNG::AID::SOS-1 was lost in the FGFR mutant *egl-15(n1458),* indicating that of SOS-1 polarization in the SM depends on FGFR/EGL-15(5A) (Fig. 3D, F). To test the extent to which SOS-1 localization provides an intracellular readout of the position of an extracellular FGF source, we deleted the FGF homolog (*egl-17*) required for SM migration and then misexpressed *FGF/egl-17* in a small cluster of posterior cells using a fragment of the *egl-20* promoter to reorient SM migration towards the tail^31^. In *FGF/egl-17(Δ); Pegl-20>FGF/egl-17* animals, mNG::AID::SOS-1 punctae localized to the posterior ends of SMs as they migrated towards the tail (Fig. 3E, F), consistent with an instructive role for SOS-1 in directing SM migration towards FGF-expressing cells.

**Figure 3.**
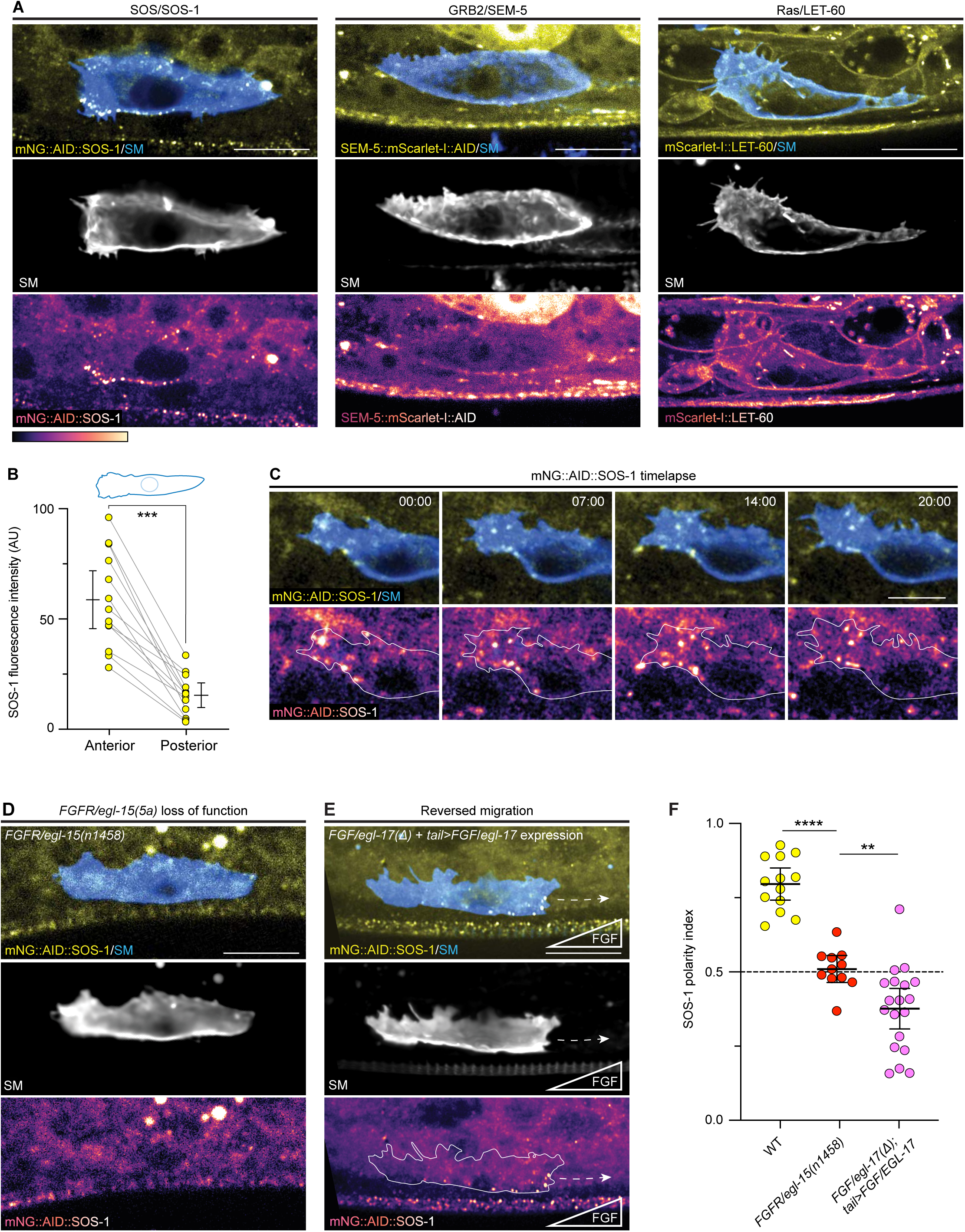
Extracellular FGF orients SOS-1 to the leading edge of migrating SMs. **(A)** Representative, live spinning disk confocal images showing endogenous mNG::AID::SOS-1, GRB-2/SEM-5::mScarlet-I::AID, or mScarlet-I::Ras/LET-60 localization during SM migration. (**B**) Quantification of mNG::AID::SOS-1 fluorescence intensity at the leading (anterior) and trailing (posterior) edges of migrating SMs. Each point represents a signal cell, and measurements from the same cell are connected by a gray line. Bars show mean and 95% confidence intervals. ***, P=0.0002, Wilcoxon matched-pairs signed rank test (two tailed). **(C)** Individual frames from a time-lapse image series showing mNG::AID::SOS-1 localization in a migrating SM. mNG::AID::SOS-1 punctae are prominent at the front of the cell and in leading edge protrusions as the cell migrates. Images correspond to frames from Video 1, and time is in noted in minutes:seconds. **(D)** Representative, live spinning disk confocal images showing endogenous mNG::AID::SOS-1 localization during SM migration in an *FGFR/egl-15(n1458)* mutant background. Note the absence of polarized mNG::AID::SOS-1 punctae at the cell membrane. **(E)** Representative, live spinning disk confocal images showing mNG::AID::SOS-1 localization during SM migration in a background where FGF/EGL-17 is misexpressed near the tail to cause SMs to migrate posteriorly. mNG::AID::SOS-1 punctae localize to the posterior end of the SM as it migrates towards a posterior FGF source. (**F**) Comparisons of mNG::AID::SOS-1 polarization in wild-type (WT), *FGFR/egl-15(n1458)*, and *FGF/EGL-17(1′); Pegl-20>FGF/egl-17* SMs. Each point represents an individual cell, and bars show the mean and 95% confidence intervals for each genotype. **, P=0.0046; ****, P<0.0001, Kolmogorov-Smirnov test. Single channel images of endogenously tagged proteins were colored using the mpl-magma look-up table in Fiji. Color represents fluorescence intensity with white as most intense and purple as least intense. All images are oriented with anterior to left and dorsal to top. Circular objects visible in both fluorescence channels are autofluorescent gut granules. Scale bars = 10µm.

### SOS-1 provides a directive cue for SM migration

As SOS-1 was required cell-autonomously for SM migration, and subcellular localization of mNG::AID::SOS-1 reflected the direction of an FGF source, we sought to test the possibility that polarized SOS activity is essential to orient migrating SMs. Alternatively, SOS-1 may be permissively required for SM migration but not responsible for the ability of SMs to navigate towards an FGF source. To test if asymmetric SOS activity is required for directed SM migration, we forced uniform SOS activity around the cell membrane using a SOS catalytic domain^62^ (SOS^cat^) fused to 2x mTurquoise2 (mT2) and the pleckstrin homology (PH) domain of *plc1δ1*, which associates with the inner leaflet of the plasma membrane^63^ (Fig. 4A). Expressing SOS^cat^::2x mT2::PH in the M lineage arrested SM migration in a wild-type background, although the phenotype was less penetrant than depletion of SOS-1 or FGFR (Fig. 4B, E). Intriguingly, SOS^cat^::2x mT2::PH expression also resulted in dorsal displacement of 16% of SMs, which is approximately triple the incidence of dorsoventral defects seen in *FGFR/egl-15(n1458)* animals (Fig. 4C; Table S1). *Phlh-8>*SOS^cat^::2x mT2::PH also prevented SMs from migrating towards an ectopic, posterior FGF source in *FGF*/*egl-17* null animals with *FGF/egl-17* expressed near the tail (Fig. 4C, E), further validating that asymmetric SOS activity is required for SMs to migrate towards an FGF source, regardless of direction.

**Figure 4.**
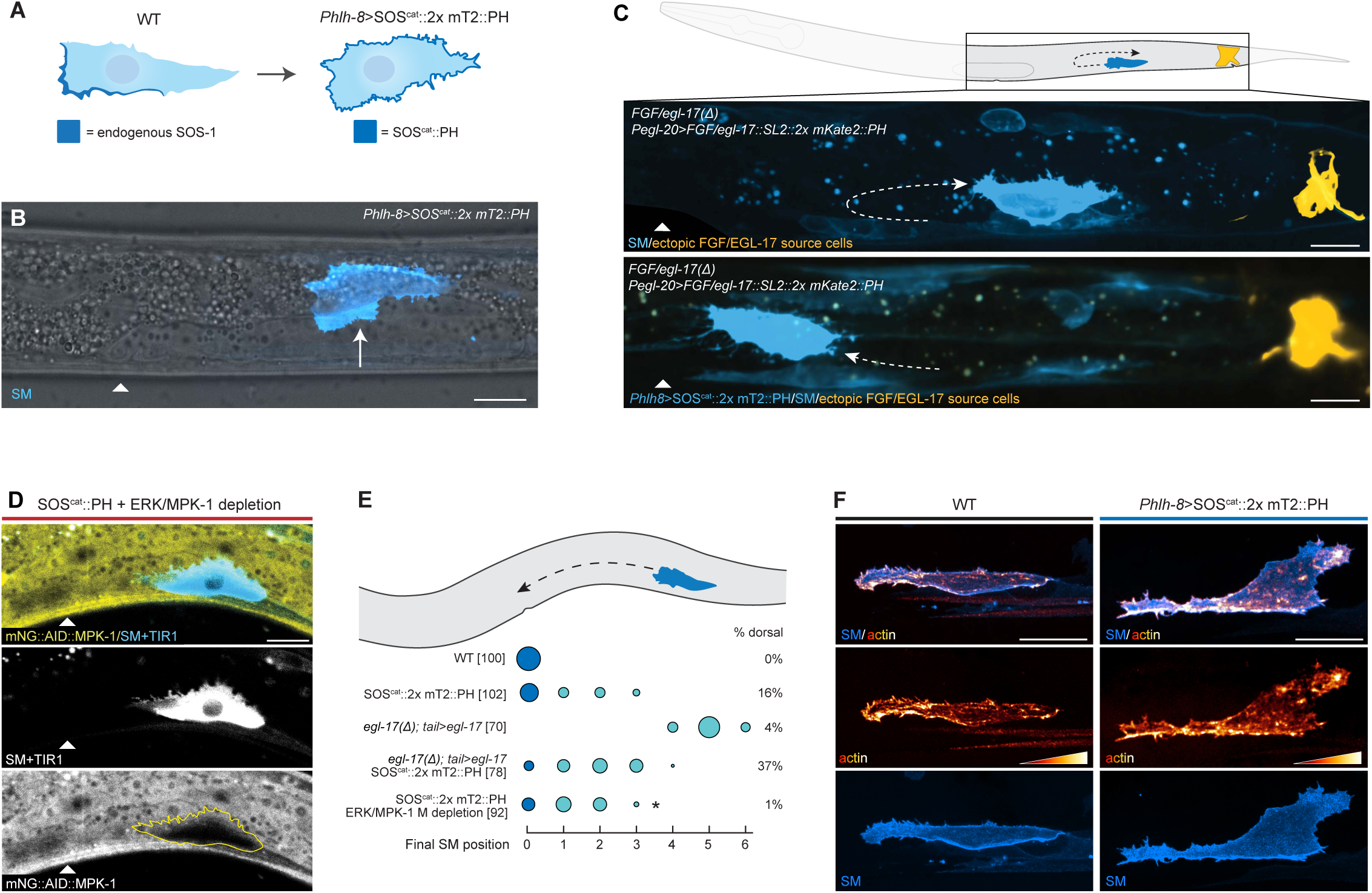
Spatially uniform SOS activity in SMs disrupts cell migration and cytoskeletal organization. **(A)** Schematic of native SOS-1 localization in a migrating SM (see Figure 3A) contrasted with spatially uniform SOS activity caused by membrane-associated *Phlh8*>SOS^cat^::2x mT2::PH. **(B)** Representative, live spinning disk confocal image showing migration defect caused by *Phlh8*>SOS^cat^::2x mT2::PH in a wild-type background. The SM (arrow) failed to migrate and is dorsally displaced. **(C)** Representative, live spinning disk confocal images of SM positions at the end of migration in *egl-17(1′); Pegl-20>FGF/egl-17* and *Phlh8*>SOS^cat^::2x mT2::PH; *egl-17(1′); Pegl-20>FGF/egl-17* animals. Expressing *FGF/egl-17* in the tail normally reorients SMs to migrate posteriorly in an *egl-17(1′)* background. This migration is suppressed by *Phlh8*>SOS^cat^::2x mT2::PH. Dashed white arrow denotes the migratory path taken by the SM. **(D)** Representative, live spinning disk confocal images showing SM positioning in a *Phlh8*>SOS^cat^::2x mT2::PH; mNG::AID::ERK/MPK-1 animal where endogenous ERK/MPK-1 was depleted in the M lineage. Depleting ERK/MPK-1 suppressed the dorsoventral defects caused by membrane-associated SOS^cat^ and enhanced the anteroposterior migration defect. **(E)** Dot plot summarizing SM positioning in animals expressing SOS^cat^::2x mT2::PH in the M lineage in different genetic backgrounds. Area of each dot is proportional to the frequency of cells stopping at that position. *, P<0.05 compared to *Phlh-8>SOS^cat^::2x mT2::PH*, Fisher’s exact test. Numbers of SMs analyzed for each strain are shown in brackets. Source data available in Table S1. (**F**) Representative, live spinning disk confocal images showing actin organization in migrating SMs visualized by *2x mKate2::moesin actin binding domain* in wild-type and *Phlh8*>SOS^cat^::2x mT2::PH animals. A glow look-up table, where white represents the highest fluorescence intensity and red the lowest, was used to visualize actin localization. White triangles in B-D indicate the normal endpoint of SM migration. Circular objects visible in both fluorescence channels are autofluorescent gut granules. All images are oriented with anterior to left and dorsal to top. Scale bars = 10µm.

To rule out the possibility that SOS^cat^::2x mT2::PH arrests SM migration through hyperactivating ERK, we used auxin-inducible degradation to deplete endogenous mNG::AID::ERK/MPK-1 in SMs in *Phlh-8*>SOS^cat^::2x mT2::PH animals. Intriguingly, ERK/MPK-1 depletion in SMs suppressed the dorsal migration defects caused by SOS^cat^::2x mT2::PH but enhanced the anterior migration defects (Fig. 4D, E). This result suggests that increased ERK activity in SOS^cat^::2x mT2::PH SMs can partially rescue their ability to migrate anteriorly, although ERK/MPK-1 is not required in the SMs for their normal migration. Additionally, this result indicates that elevated ERK activity affects dorsoventral SM positioning separately from anterior migration. To investigate how uniform SOS^cat^ localization impaired SM migration, we visualized actin in *Phlh-8*>SOS^cat^::2x mT2::PH animals using 2x mKate2 fused to a Moesin actin-binding domain^64^. In many types of migrating cells, including SMs, actin is enriched in protrusions at the leading edge^31,65^. However, in *Phlh-8*> SOS^cat^::2x mT2::PH animals, we observed misdirected, actin-rich protrusions around the periphery of the cell near the point where migration typically failed (52/68 cells; Fig. 4F). These findings suggest that uniform SOS^cat^ activity around the cell membrane disrupts cell migratory behavior by altering cytoskeletal dynamics.

Because GRB2 typically links SOS to activated growth factor receptors^60^, we next tested the extent to which subcellular GRB2/SEM-5 polarization is also required for SM migration. In an attempt to uniformly target GRB2/SEM-5 to the cell membrane without disrupting its function, we used the *hlh-8* promoter to express a modified version of SEM-5::mNG with its SH2 domain replaced with the *plc1δ1* PH domain (Fig. S4). This transgene disrupted migration in approximately 15% of SMs, which may be attributable to its only weak membrane localization (Fig. S4) compared to SOS^cat^::2x mT2::PH. In *Phlh-8*>SEM-5::PH::mNG animals, approximately half of cells with anterior migration defects also exhibited dorsal displacement, similar to the increased frequency of dorsal migration defects with uniform SOS^cat^ localization.

### Activated Ras/LET-60 acts permissively in SM migration

Next, we investigated the extent to which Ras/LET-60 functions cell-autonomously as a permissive versus directive signal for FGF-directed SM migration. A permissive role for *Ras/let-60* was previously inferred based on normal SM positioning in *let-60(n1046[G13E])* gain-of-function mutants and the ability of *n1046[G13E]* to rescue SM migration in the absence of FGF signaling^33^. To determine the effect of constitutively active Ras in the context of cell type-specific gain of function, we first confirmed that M lineage-specific *Phlh-8>let-60(G13E)* expression did not affect SM migration in a wild-type background and partially rescued SM migration defects in an *egl-15(n1458)* background (Fig. S5; Table S1). To assess the possibility that stronger Ras/LET-60 gain of function may interfere with SM migration, we then expressed *let-60(G12V)* in the M lineage in a wild-type background. While substitutions at G13 permit some GTP hydrolysis, G12V substitution is expected cause stronger gain of function by locking Ras/LET-60 in its GTP-bound active state^66,67^. Similar to *let-60(G13E)* misexpression, SMs expressing *let-60(G12V)* migrated to the correct position in the majority of animals. Interestingly, *Phlh-8>let-60(G12V)* animals exhibited a low-penetrance defect in M lineage differentiation that led to ectopic SMs (4/72), which was not observed in *let-60(n1046)* or *Phlh-8>let-60(G13E)* animals. Of *Phlh-8>let-60(G12V)* animals with normal numbers of SMs at the early L3 stage, we observed slight undermigration for 3/68 cells (Fig. 5A, F; Table S1). To test the extent to which constitutively activated Ras/LET-60 might affect SM migration through mechanisms distinct from the ERK-independent role of Ras/LET-60 in normal SM migration, we depleted mNG::AID::ERK/MPK-1 in the M lineage in *Phlh-8>let-60(G12V)* animals. M lineage-specific ERK/MPK-1 depletion in this background did not suppress the incidence of M lineage differentiation defects (8/132), which suggests this phenotype does not require Ras-ERK signaling. However, ERK/MPK-1 depletion led to a striking overmigration phenotype where many cells migrated anteriorly past their normal endpoint (45/124; Fig. 5B, F), while a smaller number undermigrated (14/124; Fig. 5F; Table S1). These findings suggest that activated Ras/LET-60(G12V) can promote anterior SM migration cell autonomously through an ERK-independent mechanism, which is counteracted by ERK/MPK-1 to facilitate normal SM positioning when Ras/LET-60 is abnormally activated.

**Figure 5.**
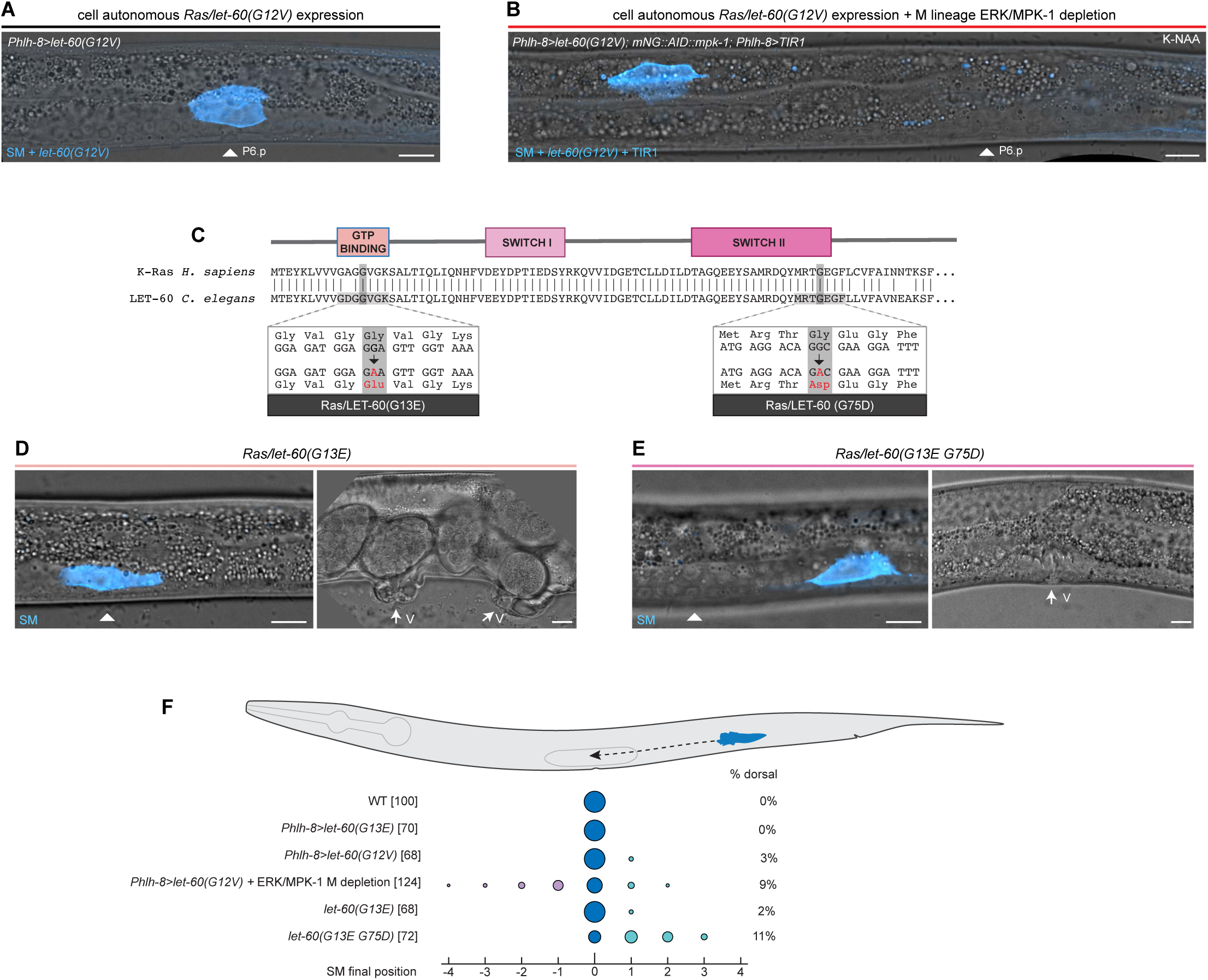
Cell type-specific Ras gain of function and an intragenic revertant mutation suggest permissive roles for Ras/LET-60 in SM migration and distinct Ras signal transduction mechanisms in SMs and VPCs. (**A**) Representative image of normally positioned SM in a *Phlh-8>let-60(G12V)* animal. (**B**) Example of overmigrated and dorsally positioned SM in a *Phlh-8>let-60(G12V)* animal with mNG::AID::ERK/MPK-1 depleted in the M lineage. **(C)** Schematic diagram showing alignment of human K-Ras and *C. elegans* LET-60 with the G13E (gain-of-function) and G75D (intragenic revertant) mutations emphasized. **(D)** Representative, live images of *Ras/let-60(G13E)* mutant animals showing normal SM positioning at the L3 stage (left) and a multivulval phenotype in a young adult (right). (**E**) Representative, live images of genome-edited *Ras/let-60(G13E G75D)* animals showing SM undermigration at the L3 stage and normal vulval induction at the L4 stage. See Table S2 for analyses of lethality, vulval induction, and SM migration in *let-60(n1046ku75)*. **(F)** Dot plot summarizing SM migration endpoints in strains used to dissect roles for activated *Ras*/*let-60* signaling in SM migration. Area of each dot is proportional to the frequency of cells stopping at that position. Numbers of SMs analyzed for each strain are shown in brackets. Source data available in Table S1. White triangles indicate the normal endpoint of migration in A, B, D, and E. v with arrow denotes vulva in D and E. All images are oriented with anterior to left and dorsal to top. Scale bars = 10µm.

### The role for Ras/LET-60 in SM migration is genetically separable from canonical Ras-ERK-dependent developmental processes

As Ras/LET-60 is required cell-autonomously for SM migration, but overactivation confers a minimal migratory phenotype in a wild-type background, we hypothesized that SM migration may be genetically separable from other developmental processes that require Ras/LET-60. We turned to unpublished intragenic revertant mutations in *let-60(n1046gf)* that were identified in a screen for suppressors of activated *Ras/let-60.* Mutations identified in this manner defined much of the signal transduction pathway downstream of Ras in *C. elegans*^9^. One intragenic revertant, *let-60(n1046ku75)* suppressed the multivulval phenotype of *let-60(n1046)* and selectively impaired SM migration relative to other developmental processes. SM migration defects were observed in 93% of *let-60(n1046ku75)* animals with only minimal effects on larval survival and vulval induction (Table S2). *ku75* is a point mutation in *let-60(n1046)* that causes a Glycine>Aspartic acid substitution at amino acid position 75 (G75D) in the switch II region of Ras/LET-60 (Fig. 5C; Fig. S6), which is predicted to interfere with binding to SOS-1 (Fig. S7; Video S1). To eliminate the possibility that linked mutations affecting other genes might underlie the SM migration phenotype in *let-60(n1046ku75)* animals, we used co-CRISPR to generate a new *let-60(G13E)* allele in an unmutagenized background followed by a second co-CRISPR genome edit to generate *let-60(G13E G75D)* animals. Similar to the intragenic revertant mutation, CRISPR-edited *let-60(G13E G75D)* suppressed the multivulval phenotype caused by *let-60(G13E)* (Fig. 5D, E) without noticeable larval lethality or early developmental abnormalities. We observed SM migration defects in the majority of genome edited *let-60(G13E G75D)* animals (Fig. 5E, F), confirming that the combination of G13E and G75D substitutions more severely disrupts *Ras/let-60* activity in SM migration compared to its essential, ERK-dependent functions in other aspects of development.

### Branched actin regulators control SM protrusive dynamics but are dispensable for FGF-directed migration

Given our findings that homologs of GRB2, SOS, and Ras were required cell-autonomously for SM migration, we next sought to identify downstream proteins linking these signal transduction components to the cytoskeletal remodeling required for directed cell migration. Because SMs migrate with actin-rich, lamellar protrusions at the leading edge^31^, we focused on cytoskeletal regulatory proteins with known roles in crawling cell motility^65,68–70^. We generated biologically functional, endogenous fluorescent protein::AID-tagged alleles for homologs of ARP2 (ARX-2), SCAR/WAVE (WVE-1), WASP (WSP-1), Rac (CED-10), RhoG (MIG-2), and ROCK (LET-502). We also tagged the homolog of the non-receptor tyrosine kinase Abl (ABL-1), which mediates Rac-GEF activity of vertebrate SOS proteins^71,72^. All of these knock-in strains had superficially normal development except for mScarlet-I::AID::CED-10, which exhibited slow growth and frequent defects in germline and somatic gonad development. For SM migration experiments, we depleted Rac/CED-10 and the related RhoG/MIG-2 simultaneously as they often act redundantly or in parallel, and individual *ced-10* or *mig-2* mutants are not reported to have SM migration phenotypes^69,73^. We found that depleting ARX-2, WVE-1, WVE-1 plus WSP-1, or CED-10 plus MIG-2 in the M lineage resulted in M lineage defects including abnormal cell division patterns and fates, which complicated identifying roles in SM migration (Fig. S8). To circumvent earlier defects in M lineage development, we treated synchronized larvae with K-NAA after SMs were born to deplete ARX-2, WVE-1, WVE-1 plus WSP-1, and CED-10 plus MIG-2 during SM migration without perturbing earlier processes. We found that SMs invariably migrated to their correct position centered over the dorsal uterine cells and P6.p when ARX-2::mNG::AID, WVE-1::mNG::AID, or WVE-1::mNG::AID plus WSP-1::mNG::AID were depleted after the SMs were specified (Fig. 6A, B; Table S1). We confirmed that TIR levels were not limiting by repeating timed ARX-2::mNG::AID depletion in a *Phlh-8>flp-D5; Prpl-28>FLP-on TIR1(F79G)* background (Fig. 6A, B; Table S1). While SMs migrated to the correct position after depleting these branched actin regulators, we observed abnormal membrane morphologies (Fig. S8). The majority of SMs were also positioned normally after simultaneous, timed depletion of the Rac and RhoG homologs mScarlet-I::AID::CED-10 and mNG::AID::MIG-2 (Fig. 6A, B), but we observed mild migration defects in a minority of SMs that indicate CED-10 and MIG-2 contribute to their final positioning (9/76; Fig. 6B; Table S1). However, the presence of somatic gonad defects in this background (Fig. S8) prevented drawing firm conclusions on cell-autonomous requirements for CED-10 and MIG-2 in SM migration given that the gonad plays essential roles in SM migration guidance. We did not observe abnormalities in M lineage development or SM positioning after continuously depleting ENA/VASP/UNC-34::mNG::AID^74^, mNG::AID::ROCK/LET-502 or ABL-1::mNG::AID in the M lineage (Fig. 6A, B; Table S1). SMs were also normally positioned in *abl-1(ok171)* null mutant animals (Fig. 6B; Fig. S9; Table S1), confirming that ABL-1 does not mediate the essential function of SOS-1 in SM migration.

**Figure 6.**
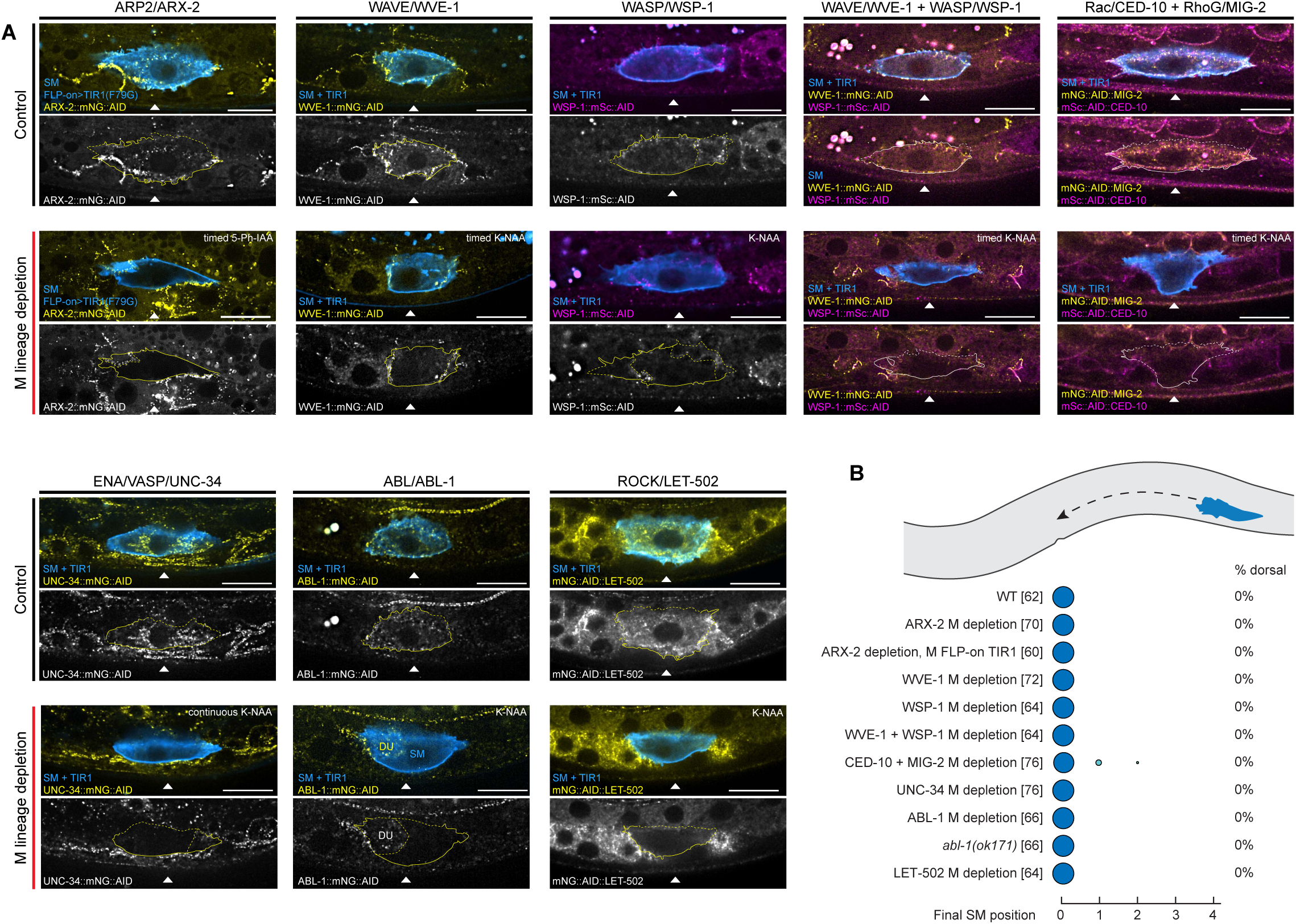
Key cytoskeletal regulators are not required cell autonomously for SM positioning. **(A)** Live, spinning disk confocal images showing representative SM positioning at the end of migration and tagged endogenous protein localization in control and M lineage depletion animals. To circumvent early defects in M lineage development, ARX-2, WVE-1, WVE-1 plus WSP-1, and CED-10 plus MIG-2 were depleted after the SMs were born. See Figure S8 for examples of abnormal M lineage development caused by earlier depletion of these proteins. WSP-1, UNC-34, ABL-1, and LET-502 depletion did not affect earlier M lineage development. SMs are outlined in images where the SM membrane marker channel (blue) is not shown. Dashed sections of the outline indicate regions where cells above or below the SM are visible in the same focal plane or where the SM membrane is not in focus. ARX-2::mNG::AID animals expressing FLP-on>TIR1(F79G) in the M lineage were treated with 5-Ph-IAA to trigger depletion. All other strains express unmodified TIR1 in the M lineage and were treated with K-NAA to trigger depletion. White triangles indicate the normal endpoint of migration. DU denotes dorsal uterine cell. All images are oriented with anterior to left and dorsal to top. Circular objects visible in all fluorescence channels are autofluorescent gut granules. Scale bars = 10µm. (**B**) Dot plot summarizing SM migration endpoints after M-lineage specific depletion of endogenously tagged proteins or in mutant strains. Area of each dot is proportional to the frequency of cells stopping at that position. Data plotted for CED-10 plus MIG-2, ARX-2, WVE-1, and WVE-1 plus WSP-1 depletion are from animals with timed depletion and scored at the 1 SM stage. Numbers of SMs analyzed for each strain are shown in brackets. Source data available in Table S1.

We next used live imaging of migrating SMs to elucidate roles for candidate cytoskeletal regulators in shaping SM protrusive architectures and migratory behaviors. Lamellar protrusions with branched actin networks at the leading edge play essential roles in cell motility in many contexts^2,75,76^. However, some cell types that typically migrate using lamellipodial crawling can migrate directionally in the absence of key branched actin regulators by altering their modes of motility^75,77,78^. While depleting candidate cytoskeletal regulatory proteins in the M lineage did not substantially affect the final position of SMs (see Fig. 6), live imaging of migrating SMs at earlier stages showed that ARX-2, WVE-1 plus WSP-1, or CED-10 plus MIG-2 depletion caused strikingly abnormal protrusive morphologies (Fig. 7A-D). SMs normally migrate with a broad, lamellar protrusion at the anterior and short microspikes or filopodia. However, ARX-2-depleted SMs migrated with aberrant leading-edge morphologies that including a single elongated protrusion that tapered to a point (27/49; Fig. 7A-D) or multiple short filopodia (14/49; Fig. 6C, D). We also observed cells that appeared to be bifurcated or multifurcated as they migrated over the somatic gonad (8/49; Fig. 7C, D). Because WVE-1 and WSP-1 can act partially redundantly in cell migration^79^, we assayed SM migratory morphologies after combined WVE-1 plus WSP-1 depletion. Similar to ARX-2 depletion, this experiment caused abnormal SM migratory morphologies including cells that migrated with a single narrow, elongated protrusion (15/21; Fig. 7A-D), multiple short filopodia (5/21; Fig. 7C, D), or bifurcation (1/21; Fig. 7C, D) consistent with deeply conserved roles for WAVE and WASP in regulating lamellipodial cell crawling^80^. CED-10 plus MIG-2 depletion caused a distinct SM migratory phenotype characterized by a majority of cells that migrated with multiple irregularly shaped, rounded pseudopods and short filopodia (16/19; Fig. 7A-D) along with a minority of cells that migrated with a single narrow protrusion (3/19; Fig. 7C, D). In addition to abnormal leading edge morphologies, we also observed trailing edge defects including membrane bubbling, thin elongated cell tails, and larger trailing cell fragments after ARX-2, WVE-1 plus WSP-1, or CED-10 plus MIG-2 depletion (Fig. S10). Cells depleted for ENA/VASP/UNC-34, ROCK/LET-502 (Fig. 7A, B, D), or ABL-1 (Fig. S9) continued to migrate with a lamellipodium at the leading edge and did not display obvious migratory defects, although these proteins may play more minor roles in SM dynamics. Our combined results demonstrated that several conserved regulators of branched actin assembly are essential for normal protrusive dynamics in migrating SMs. Yet, none of the tested proteins were essential for FGF-directed migration, which highlights considerable redundancy between cytoskeletal regulators and the ability of cells *in vivo* to adapt their migratory modes when actin regulators are depleted. Importantly, SM protrusions remained anteriorly directed in all experimental conditions, suggesting that migration guidance does not strictly require direct interactions between FGF signal transduction components and the cytoskeletal regulatory proteins tested.

**Figure 7.**
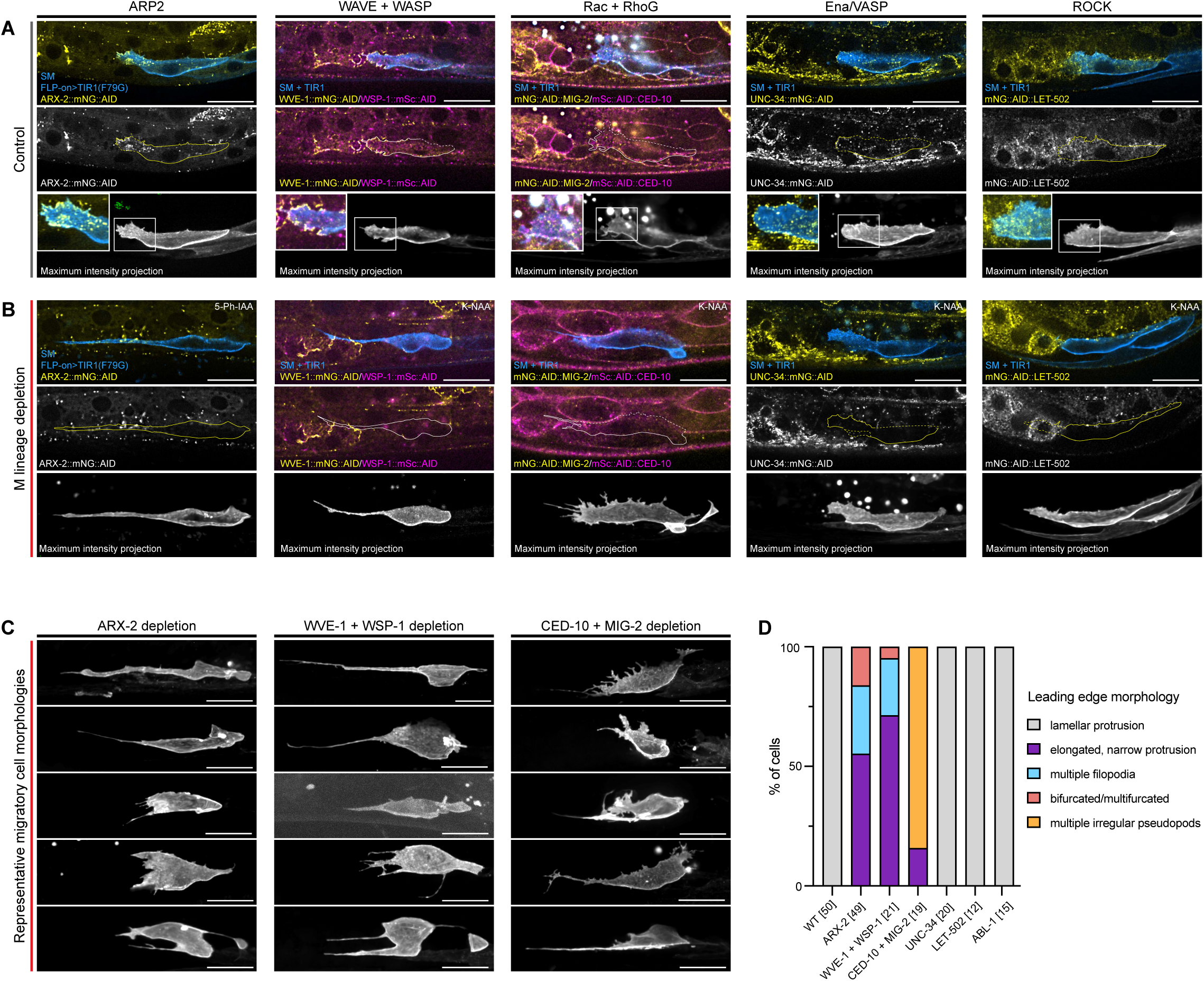
Branched actin regulators control leading edge protrusive dynamics during SM migration. (**A**, **B**) Live, spinning disk confocal images showing membrane morphology in migrating SMs and tagged endogenous protein localization in control (**A**) and M lineage depletion (**B**) animals. Images show single z-slices with the leading edge in focus for the membrane and endogenously tagged protein channels along with a maximum intensity projection to depict cellular morphology. SMs are outlined in images where the SM membrane marker channel (blue) is not shown. Dashed sections of the outline indicate regions where cells above or below the SM are visible in the same focal plane or where the SM membrane is not in focus. ARX-2::mNG::AID animals expressing FLP-on>TIR1(F79G) in the M lineage were treated with 5-Ph-IAA to trigger depletion. All other strains express unmodified TIR1 in the M lineage and were treated with K-NAA to trigger depletion. Images shown in B represent the most common cellular phenotype for each strain. (**C**) Maximum intensity projections depicting the spectrum of SM membrane morphologies caused by depleting branched actin regulators during SM migration. All images are oriented with anterior to left and dorsal to top. Scale bars = 10µm. (**D**) Quantification of leading edge morphologies in migrating SMs after cell-autonomous depletion of endogenously tagged cytoskeletal regulatory proteins. Numbers of SMs analyzed for each strain are shown in brackets.

## Discussion

RTKs regulate cell migration in many contexts during animal development, and aberrant RTK activity drives cancer and other disease states^5,81,82^. Yet, elucidating the underlying cellular and molecular mechanisms that orient migrating cells downstream of RTKs in living animals has been challenged by pleiotropy and a lack of systematic approaches to dissect functions in cell migration without perturbing other critical developmental or physiological processes. Here, we used cell type-specific depletion of 17 endogenously tagged proteins, along with gain-of-function and mutant analyses, to determine functions for key signal transduction and cytoskeletal regulatory proteins in migration of *C. elegans* postembryonic mesoderm progenitors. Our results provide a mechanism by which migrating cells orient towards an extracellular growth factor source and raise new questions about how RTK signal transduction directs cell migration in living animals. Surprisingly, we found that homologs of FGFR, GRB2, SOS, and Ras were required for SM migration towards an FGF source, but typical downstream effectors in the ERK, PI3K-Akt-mTOR, and PLCγ signaling pathways were not required cell-autonomously. Instead, our work demonstrated that FGFR-dependent, localized SOS activity orients migrating SMs towards an FGF source independent of canonical RTK effectors. We found that SOS is localized at the leading edge of migrating SMs and provides an intracellular readout of the direction of an FGF source. Uniform SOS^cat^ localization at the cell membrane disrupted polarized protrusive behaviors in migrating SMs and prematurely arrested SM migration. The function of SOS^cat^ did not depend on ERK in this context, and simultaneous ERK/MPK-1 depletion in fact enhanced migration defects in SOS^cat^::2x mT2::PH cells. As FGFs and other growth factors provide positional information in developing animals and regulate cell migration in multiple systems, our findings may be broadly applicable towards understanding the cell biological mechanisms by which FGFs and RTKs orchestrate critical aspects of animal development.

Ras and Rac family proteins are thought to be the primary targets activated by SOS^8,9,83^. This hypothesis is supported by the finding that Ras/LET-60 depletion results in SM migration phenotypes similar to SOS-1 depletion. While we found clear evidence that SOS transduces an instructive directional cue for FGF-directed SM migration, our results are consistent with a model where constitutive activation of Ras/LET-60 is largely permissive for anterior migration. Endogenous Ras/LET-60 protein is not obviously polarized in migrating SMs, *let-60(G13E)* gain-of-function mutants do not have defects in SM migration, and expressing activated *Ras/let-60(G13E)* or *(G12V)* in migrating SMs had minimal effects on cell migration in a wild-type background. However, it is possible that Ras/LET-60 activity may be polarized even if the protein is not. It is also possible that LET-60(G13E) and/or LET-60(G12V) remain sensitive to SOS-1 activity and may therefore be regulated in a polarized manner in migrating SMs. In the future, improved Ras biosensors and methods for locally activating Ras without manipulating Ras-GEFs could provide valuable insights into the mechanisms by which Ras/LET-60 mediates SM migration. Different roles for ERK/MPK-1 downstream of wild-type Ras/LET-60 versus constitutively activated alleles also complicate inferring the mechanisms used in normal cell migration based on Ras gain-of-function conditions. The finding that Ras/LET-60(G12V) promotes overmigration cell-autonomously when ERK/MPK-1 is depleted suggests that constitutive Ras activation can promote both ERK-independent anterior migration and a second ERK-dependent process that enforces normal SM positioning in a wild-type background. In contrast, in cells expressing a membrane-associated SOS catalytic domain, we found that ERK/MPK-1 depletion enhanced SM undermigration, which indicates that SOS^cat^::PH and Ras/LET-60(G12V) activate distinct cellular responses that impact migration through different mechanisms. Our results suggesting that SOS-1 instructs SM orientation while activated Ras/LET-60 acts permissively raise the question of what other effectors may act downstream of SOS-1 to execute its instructive role. Although SOS proteins can directly activate Rac in some contexts^8^, *Rac/ced-10* mutants do not have defects in SM positioning, and SMs depleted for Rac/CED-10 and RhoG/MIG-2 were still capable of migrating to the correct destination. Identifying additional proteins that interact with SOS-1 during RTK-mediated migration will also be crucial for deepening our understanding of directional cell movement.

Our investigation also revealed a novel genetic interaction between two mutations in *Ras/let-60*, G13E and G75D, that offers insights into cell type-specific signal transduction mechanisms. Ras family proteins are highly conserved in animals and play crucial roles in fundamental cellular processes including survival, proliferation, and migration^7,9,84^. Strong *Ras* gain-of-function mutations are a major driver of tumorigenesis and metastasis while other types of mutations result in a spectrum of diseases called RASopathies^7,84,85^. The intragenic revertant allele *Ras/let-60(G13E G75D)* described here demonstrates that the ERK-dependent, essential roles for *Ras/let-60* in larval viability and VPC fate specification are genetically separable from its function in SM migration. The mechanism by which the *Ras/let-60(G13E G75D)* allele differentially affects SM migration is unknown but could involve interactions with GEFs, modulators such as Ras-GAPs, or downstream effectors. Substitutions at amino acid 75 are expected to result in reduction of function based on saturation mutagenesis assays^86^, and the classical *let-60(n2021)* mutant is a G75S substitution that functions as a moderate loss of function allele^34^. Intriguingly, amino acid 75 is located in the Ras switch II domain, which plays a crucial role in tight binding of Ras to SOS^87,88^, and the G75D substitution is predicted to affect interactions between Ras and the allosteric and catalytic sites of SOS. It is possible that the combination of G13E and G75D substitutions results in an activated Ras protein that is capable of constitutive signaling but is spatially dysregulated within cells by virtue of an impaired ability to bind SOS. In this scenario, *Ras/let-60(G13E G75D)* may retain sufficient function in contexts where SOS-dependent subcellular localization of Ras activity is not critical but fail to function in contexts where the ability to bind SOS is essential. While the underlying biochemical mechanism is speculative, the phenotypes observed in *Ras/let-60(G13E G75D)* animals reveal a novel genetic interaction by which an activating mutation interacts with a putative loss-of-function substitution in the switch II region of *Ras/let-60* to differentially affect downstream responses.

Developmental roles for essential cytoskeletal regulators are often challenging to dissect in living animals due to lethality and pleiotropic effects. Thus, our understanding of how these proteins function in cell migration comes largely from experiments in cultured cells, with associated caveats^65,75^. Intriguingly, we found that cell type-specific depletion of key cytoskeletal regulators affected SM protrusive dynamics but did not substantially impact the intrinsic ability of SMs to migrate or navigate to the correct destination. These findings parallel experiments demonstrating some cell types that typically migrate with lamellar protrusions can functionally compensate for loss of ARP2/3 complex activity by altering their mode of migration^75,77,78,89^. We found that depleting ARX-2 or WVE-1 plus WSP-1 led to similar cell migratory phenotypes, which showed that assembly of branched actin networks is required for lamellipodia formation in SMs. However, SMs compensated for the loss of branched actin polymerization by migrating using an elongated, pointed protrusion or multiple filopodia. In the future, it will be intriguing to identify the cytoskeletal regulators that mediate SM migration in the absence of ARP2/3 complex activity and assess their functions in normal migration. We found that homologs of the related small GTPases Rac and RhoG are together essential for normal SM migratory behaviors, but their depletion conferred a distinct cellular phenotype characterized by numerous short and irregularly shaped pseudopods. While Rac family GTPases are key activators of the WAVE regulatory complex^90–92^, the unique phenotype caused by CED-10 plus MIG-2 depletion suggests these proteins perform additional functions in regulating cytoskeletal architecture in migrating SMs. The SM bifurcations and cell fragments observed in some cases were also reminiscent of the loss of cell integrity observed in migrating distal tip cells when Rac activity is compromised^93^.

### An endogenously tagged protein toolkit to dissect RTK signaling and cell dynamics *in vivo*

Our systematic approach to depleting endogenous signal transduction and regulatory proteins in specific cell types was a critical factor in dissecting how an RTK regulates cell migration in a living animal. In doing so, we generated a toolkit of endogenously tagged proteins that provides new tools to elucidate cellular and developmental mechanisms *in vivo*. These include key proteins involved in SOS, Ras, ERK, PI3K, Akt, and mTOR signaling along with cytoskeletal regulators important for many aspects of cell physiology and cellular behavior^4,94–96^. Notably, this toolkit includes functional mScarlet-I and AID-tagged alleles for Ras/LET-60, which is a protein of intense interest due to its biomedical significance that has been recalcitrant to endogenous tagging without disrupting its function^45^. Although incomplete protein depletion may contribute to negative phenotypes using AID-tagged endogenous proteins, comparisons to mutants and/or the presence of depletion phenotypes suggests these tools are capable of ablating or severely reducing tagged protein functions. With some promoters, TIR1 levels can be limiting for degrading AID-tagged proteins. However, out of the endogenous proteins that we targeted in SMs, we observed strong depletion phenotypes for the three genes with highest expression in SMs based on L2 scRNAseq data^47^ (*egl-15*, *let-60*, and *sos-1*; Figure S11). Because RTK signaling and cytoskeletal remodeling have critical functions in many developmental and homeostatic processes^5,65,81^, the endogenously tagged proteins generated in this study provide a powerful toolkit for researchers exploring a wide range of biological questions. Together, our findings shed light on how RTK signaling drives directed cell migration in living animals and highlight the diversity and intricacy of signal transduction and cell migratory mechanisms that mediate animal development.

## Resource availability

### Lead contact

For further inquiries or requests for obtaining reagents and resources generated in this study, please contact. Ariel Pani at.

### Materials availability

*C. elegans* strains generated in this study will be deposited at the CGC (https://cgc.umn.edu/) upon manuscript acceptance. Plasmids and other reagents will be shared on request.

### Data and code availability

Source data used to analyze SM positioning are summarized in Table S1. Other raw data will be made available upon request. No code was generated in this study.

## Acknowledgements

pWZ234 was a gift of David Matus. We also thank Min Han for supporting the genetic screen that originally identified *let-60(n1046ku75)* and Arun Dutta for their feedback on the manuscript. Some strains were provided by the CGC, which is funded by the NIH Office of Research Infrastructure Programs (P40 OD010440). This work was funded by National Institutes of Health grants R35GM142880 (AMP), R35GM136315 (MVS), and R35GM144237 (DJR) and fellowship F31HD112152 (TVG).

## Author contributions

Conceptualization, TVG and AMP; methodology, TVG., LL, JIM, NRR, MCL, DJR, MVS, AMP; formal analysis, TVG, AMP; investigation, TVG, MVS, LL, AMP; data curation, TVG, AMP; writing – original draft, TVG; writing – review & editing, TVG, MVS, DJR, AMP; visualization, TVG, AMP; project administration, AMP; funding acquisition, TVG, MVS, DJR, AMP.

## Declaration of interests

The authors declare no competing interests.

## STAR Methods

### Experimental model and study participant details

All experiments used *Caenorhabditis elegans* hermaphrodites. *C. elegans* were grown on Nematode Growth Medium (NGM) plates seeded with *Escherichia coli* OP50^97^. Worms were maintained and experiments were performed at room temperature (22 °C). *C. elegans* strains used in this study are listed in the Key Resources table.

### Method details

#### Plasmid construction

Repair templates for genome engineering were cloned constructed using Gibson assembly and Cas9 + sgRNA plasmids were cloned using site-directed mutagenesis as described elsewhere^98,99^. Sequences used for repair template design were downloaded from Wormbase^100^, and plasmids were designed using ApE^101^. Primers and double stranded synthetic DNA fragments (gBlocks) were ordered from Integrated DNA Technologies (IDT). Constructs were transformed into chemically competent *E. coli* 5-alpha cells (NEB C2987). Plasmid DNA was extracted using either Qiagen (Qiagen 27104) or Invitrogen PureLink HQ (Invitrogen K210010) miniprep kits. Plasmids were verified by Sanger sequencing (GENEWIZ) and/or whole-plasmid nanopore sequencing (Plasmidsaurus) before use. To generate the sem-5::PH(internal)::mNG construct (pTG421), the internal SH2 domain (amino acids ITRNDAEVLLKKPTVRDGHLVRQCESSPGEFSISVRFQDSVQHFK) where replaced with a PH domain from *Rattus norvegicus plc1δ1* that is common used for plasma membrane localization of fluorescent proteins in *C. elegans*^102^. To enhance expression efficiency, the DNA sequences coding for *egl-17*, the SOS catalytic domain^62^, and *let-60(G13E)* were codon-optimized using the *C. elegans* codon adapter tool^103^ (https://worm.mpi-cbg.de/codons/cgi-bin/optimize.py). We used a germline codon optimized^104^ sequence for the *mScarlet-I::let-60* knock-in. A comprehensive list of all plasmids generated or used in this study, including the appropriate plasmid backbones and vectors used for cloning, is provided in Reagents and Resources. Additional details on plasmid construction are available upon request. Promoter sequences used are provided in Supplemental File 1.

#### Genome engineering and transgenesis

Fluorescent protein::AID knock-in strains were generated using Cas9-triggered homologous recombination with a self-excising selection cassette^98^ using an injection technique with a paintbrush as previously described^105^. Fluorescent protein insertion sites were selected based on protein domain function and structure where possible. Guide RNA target sites were selected based on proximity to the desired insertion site and predicted off-target sites using ChopChop^106^ (https://chopchop.cbu.uib.no/). When the guide sequence was present in the repair template itself, we introduced synonymous mutations to prevent cleavage of the repair template by Cas9. Guide sequences and insertion sites are detailed in Supplemental File 1. Due to the compact structure of Ras/LET-60, we found it was necessary to use a 30 amino acid flexible linker to generate biologically functional knock-in alleles. Due to low efficiency of the guide RNA targeting the *Ras/let-60* N-terminus, we inserted a known, high-efficiency gRNA site derived from *dpy-10* at the N-terminus of endogenous *Ras/let-60* using co-CRISPR. This allele was viable as a homozygote and confirmed by sequencing. We then used this efficient guide RNA site to engineer N-terminal *Ras/let-60* knock-ins using co-CRISPR. We were unable to recover homozygous mScarlet-I::AID::Ras/LET-60 knock-in animals, suggesting this fusion was not functional. However, we were able to separately generate functional mScarlet-I::Ras/LET-60 and AID::Ras/LET-60 alleles that allowed for visualization and depletion of the endogenous protein in complementary strains. Microinjection mixes for PCR fragment-based knock-ins were templates were prepared as previously described^107^, and crRNA and ssODN sequences are provided in Reagents and Resources table and Supplemental File 1. The PI3K/AGE-1::mNG::AID knock-in allele was generated by inserting mNG::AID before the WIF (Wortmannin-Insensitive Fragment) motif to preserve protein function (see Supplemental File 1). Because the SEC truncated *age-1* in the initial insertion strain, we heat-shocked heterozygous *age-1::mNG^SEC^AID* animals to excise the SEC and isolated heterozygous *age-1::mNG::AID/WT* animals by picking fluorescent animals with wild-type movement under a fluorescence microscope. APL127 was then derived from homozygous self-progeny of a heterozygous SEC-excised animal. APL306 was constructed by crossing APL127 and APL173. The SEC could not be excised from the *Phlh-8>sem-5::PH::mNG SEC* transgene in the parent strain of APL481 despite repeated heat shock attempts. To accurately score SM positioning in this strain, we eliminated the rolling phenotype caused by the SEC by knocking out the synthetic *sqt-1(d)* allele using a crRNA specific to the codon-optimized *sqt-1* sequence used in the SEC and a single stranded oligonucleotide repair template (Supplemental File 1). SpCas9 protein and TracrRNA were then co-injected, following the standard Co-CRISPR protocol^108^, and non-rolling worms were selected in the next generation. All endogenously tagged strains used in this study appeared superficially wild-type, with the exception of APL496 (*mNG::AID::RhoG/mig-2 + mScarlet-I::AID::Rac/ced-10*). While the SMs typically migrated correctly in this strain, we observed frequent defects in germline and somatic gonad development (Fig. S8) along with occasional abnormal SM divisions after migration, indicating that this combination of tags causes partial loss of function. The M-lineage *Prpl-28> FLP-on TIR1(F79G)::T2A::2x mKate2::DHB* strain was constructed by inserting a single copy *Phlh-8>flp-D5* transgene into strain DQM1113^50^. A comprehensive list of all strains, endogenous protein tags, linker sequences, guide RNA targets, and specific insertion locations can be found in the Reagents and Resources table and/or Supplemental File 1.

Strains APL430, APL499, and APL885 were generated using co-CRISPR with single-stranded oligodeoxynucleotide repair templates. Repair templates and guide RNAs were designed following established protocols, with *rol-6* used as a co-injection marker^108^. SpCas9 protein, single stranded DNA repair templates, tracrRNA, and crRNAs were ordered from Integrated DNA Technologies. Sequences are available in Supplemental File 1. The *FGFR/egl-15(ljf54)* mutant was isolated by screening for egg laying deficient animals in the F2 generation. The *Ras/let-60(G13E)* mutant *let-60(ljf52)* was isolated by screening for the semidominant multivulval phenotype caused by Ras/LET-60 gain of function. The *Ras/let-60(G13E G75D)* allele *let-60(ljf53)* was generated by co-CRISPR in the *let-60(ljf52)* background. *let-60(ljf53)* was isolated by screening for F2 animals where the multivulval phenotype was suppressed. Oligonucleotide repair template and crRNA sequences are provided in Supplemental File 1. Genomic edits were sequence-verified by sequencing PCR products amplified from genomic DNA of homozygous animals.

All transgenic strains were generated by single copy insertions at safe harbor sites using Cas9-triggered homologous recombination with the SEC approach^98^. Transgenes were inserted near the ttTi4348 insertion site on Chr I or a site we established for efficient single copy insertions on ChrIV^31,109^. Guide RNA and homology arm sequences used for transgenes are provided in Supplemental File 1.

#### Endogenously tagged protein depletion

AID-tagged proteins in strains expressing TIR1 were depleted by treating with the water soluble, synthetic auxin 1-Naphthaleneacetic Acid Potassium Salt (K-NAA)^38^. To deplete AID-tagged proteins, we treated entire plates of worms by adding 400 µL of a 100 mM K-NAA solution for a final concentration of 4mM per plate. Plates were then allowed to dry with the lid ajar. The K-NAA stock solution was prepared in nuclease free water, stored at 4°C, and replaced every 3-4 weeks. Paired control plates were treated with nuclease free water was used for the control plates. To deplete endogenously tagged proteins in strains expressing TIR1(F79G), we treated with the auxin derivative 5-Ph-IAA at a final concentration of 1mM per plate^110^. 5-Ph-IAA was reconstituted in DMSO and diluted in water to allow for even spreading on plates. Control plates utilized an equivalent concentration of DMSO diluted in water. For timed protein depletion experiments, we used bleach synchronization^97^ to produce plates of synchronized L1 stage animals. Prior to depletion, we confirmed developmental staging in a subsample of animals on each plate using spinning disk confocal microscopy. Staging was determined based on external anatomy, vulval morphology, and M lineage development^111^. Once developmental stages were verified, K-NAA or 5-Ph-IAA was immediately added to the plates. Unless otherwise noted, all depletions were performed using a single copy TIR1::F2A::2x mTurquoise2::PH transgene driven by the *hlh-8* promoter.

We took several approaches to validate depletion phenotypes for target proteins whose depletion did not result in SM migration phenotypes. To confirm that our AID-tagged alleles were capable of ablating protein function, we also used whole-body depletion with a ubiquitously expressed TIR1 transgene. For all strains described here, ubiquitous depletion phenocopied the known null mutant phenotypes, indicating these alleles are capable of eliminating the known functions of the targeted proteins. For genes with viable null mutations, we examined M lineage phenotypes and SM migration in null mutants (*age-1(m333), ksr-1(n2526), abl-1(ok171)*).

Because the SMs are thin and wrap closely over the somatic gonad, it is often impossible to capture confocal images that show tagged endogenous protein expression in the SM without including the surfaces of underlying uterine cells. Fluorescence from these cells readily bleeds through into the same image plane as the SM (Fig. S12) but can be recognized by its appearance in the complete z-stack.

#### Identification of let-60(n1046ku75)

The *let-60(n1046ku75)* mutant was isolated in a mutagenesis screen to identify factors acting downstream of Ras/LET-60 in vulval development. Briefly, *let-60(n1046gf)* hermaphrodites were mutagenized with 50 mM ethyl methanesulfonate (EMS) and allowed to self-fertilize for two generations. F2 grand progeny were screened for non-multivulval animals to identify suppressors of activated Ras/LET-60. Candidate suppressor strains were backcrossed to the parent *let-60(n1046gf)* strain, and mutations were placed on a genetic map using phenotypic mapping strains^43,112^. To identify the molecular lesion responsible for *ku75, let-60* coding sequences were PCR amplified from the parental *let-60(n1046gf)* strain and *ku75* and directly sequenced.

#### Microscopy and image processing

For live imaging, multiple worms from the same strain or experimental manipulation were anesthetized in a drop (∼4 µL) of 0.2 mM levamisole in M9 buffer on a 170 ± 5 μm glass coverslip and allowed to rest for several minutes before being mounted on 2-5% agarose pads^113^. Worms were imaged or scored for SM positioning within 1 hour of mounting. All images were acquired by live imaging using either a Yokogawa CSU-X1 spinning disk confocal equipped with a Hamamatsu ORCA Fusion BT CMOS camera, or a Yokogawa CSU-W1 SoRA spinning disk confocal equipped with a Hamamatsu ORCA Quest qCMOS camera. Both images systems were mounted on Nikon Ti2 inverted microscope stands with perfect focus. Imaging was performed using excitation lasers at 445 nm, 514 nm, 561 nm, or 594 nm depending on the fluorescent protein. For the CSU-W1 SoRA system, imaging was performed with either a Nikon SR Plan Apo IR 60X/1.27 water immersion objective or a Nikon Apo TIRF 100X/1.49 oil immersion objective. The CSU-X1 system utilized a Nikon Apo TIRF 100X/1.49 oil objective, a Nikon Apo TIRF 60X/1.49 oil objective, or a Nikon S Fluor 40X/1.30 oil objective. Images were acquired using Nikon NIS-Elements software (Versions AR 5.42.06 and AR 5.42.04) with z-step sizes, camera exposure times, and laser intensities adjusted based on fluorescence intensities of the tagged proteins. Images of endogenously tagged proteins were deconvolved, and images of plasma membrane markers were denoised, using Nikon NIS Elements software. For figure preparation, we adjusted brightness and contrast using FIJI ^114^. The mpl-magma look up table in Fiji was used to better visualize protein localization for some images and in Video 1. Some animals were computationally straightened using Fiji to more clearly depict SM positioning.

#### Quantification and statistical analysis

To compare SM positioning at the endpoint of migration between strains, we scored SM positioning at L3 developmental stages when SMs have normally completed their migration but prior to their initial division. To ensure accurate scoring in animals with SM migration failures, we used VPC, gonad, and body wall muscle development to establish developmental timing. For normally positioned and undermigrated SMs, positions were scored based on established landmarks including the positions of the vulval precursor cells and M lineage-derived body wall muscles that are visible using the same transgene used to visualize SM positions (Fig. S1). The SM position was scored based on the visual center of each cell (two SMs per animal), and scored on a scale from 0 (representing a cell centered correctly over P6.p) to 7 (the most posterior position observed) to reflect the range of migration phenotypes observed. Any position other than 0 represents a migration defect. For overmigrated SMs, we divided the region between P6.p and the posterior end of the pharynx into five evenly sized quintiles. At least 30 animals (60 SMs) were included for each experimental condition with no data points excluded from analysis post-hoc. Ectopic SMs in *age-1(m333)* mutants, *Phlh-8>let-60(G12V)* transgenics, and after depleting tagged endogenous proteins were scored using cell shape, position, and/or relative levels of *Phlh-*8-driven transgene expression to classify cells as SMs or other M lineage cell types. At the L3 stage, SMs and other M lineage-derived cell types can be distinguished by reduced expression of *Phlh-8*>driven transgenes in body wall muscles and coelomocytes versus increased expression in SMs. Statistical analyses to compare frequency distributions of SM positions were conducted using Fisher’s exact test (two-tailed) calculated in Microsoft Excel (Version 16.89.1). For manipulations that affected dorsoventral SM positioning, we used a standard contingency table and chi-squared test to compare the percentages of dorsoventral defects between strains. To quantify SOS-1 localization in migrating SMs, we first made maximum intensity projections of confocal z-slices to visualize the anterior and posterior ends of the SMs, which were often in different z-planes. We cropped these projections to the size of the SM and used Fiji to threshold the top 10% of pixel values to reduce background. To compare SOS-1 levels at the leading and trailing edges in Fig. 3B, we measured the mNG::AID::SOS-1 fluorescence intensity in the leading region of the SM anterior to the nucleus compared to the region posterior to the nucleus using Fiji. In Fig. 3F, we calculated the SOS-1 polarity index from thresholded images by dividing the fluorescence intensity of the anterior region of the cell by the sum of the anterior and posterior regions of the cell. For all fluorescence intensity measurements, regions of interest were drawn based on a simultaneously acquired plasma membrane marker channel. Wilcoxon matched-pairs signed rank and Kolmogorov-Smirnov tests were performed using Graphpad Prism 10.5.0. Numbers of cells or animals analyzed and details of statistical tests are provided in individual figure legends. P-values were considered significant at p<0.05 with * representing p<0.05, ** representing **, p<0.01; *** representing p<0.001, and **** representing p<0.0001 in figures. Error bars represent 95% confidence intervals.

## Figure and Video Legends

**Video 1. mNG::AID::SOS-1 localizes to dynamic punctae in leading edge protrusions during SM migration.** Time-lapse imaging of endogenously tagged mNG::AID::SOS-1 in a migrating SM. Video corresponds to still images in Figure 4B. Anterior is to left and dorsal to top. Images were acquired at 1 minute intervals.

## Supplemental Information

**Document S1.** Includes Figures S1 – S12, Tables S1 and S2, and Supplemental File 1

**Video S1**. Related to Figure S7.

## Key Resources Table

| REAGENT or RESOURCE | SOURCE | IDENTIFIER |
| --- | --- | --- |
| Chemicals, peptides, and recombinant proteins |  |  |
| Potassium 1-Naphthaleneacetate (K-NAA) | Fisher Scientific | N000625G |
| 2-(5-phenyl-1H-indol-3-yl)acetic acid (5-Ph-IAA) | MedChemExpress | HY-134653 |
| Hygromycin B (50 mg/mL) (Gibco) | Fisher Scientific | 10-687-010 |
| **Experimental models: Organisms/strains** |  |  |
| *C. elegans*: wild-type | CGC | N2 |
| *C. elegans*: *ljfSi2[Phlh-8>mKate2::Dme moesin ABD::F2A::2x mTurquoise2::PH::3xHA::tbb-2 3'UTR loxN ttTi4348] I* | Gibney and Pani, 2025 | APL5 |
| *C. elegans*: *ljfSi3[Phlh-8>TIR1::F2A::2x mTurquoise2::PH::3xHA::tbb-2 3'UTR loxN ttTi4348] I* | This paper | APL34 |
| *C. elegans*: *ljfSi10[Phlh-8>2xmTurquoise2::PH::3xHA::tbb-2 3'UTR loxN ttTi5605] II* | This paper | APL40 |
| *C. elegans*: *ljfSi3[Phlh-8>TIR1::F2A::2x mTurquoise2::PH::3xHA::tbb-2 3'UTR loxN ttTi4348] I*; *egl-15(ljf34[egl-15::mNG::3xFlag::AID]) X* | This paper | APL59 |
| *C. elegans*: *ljfSi3[Phlh-8>TIR1::SL2::2x mTurquoise2::PH::3xHA::tbb-2 3'UTR loxN]*, *I*; *age-1(ljf11[age-1::mNG::3xFlag::AID])*, *II* | This paper | APL127 |
| *C. elegans*: *ljfSi3[Phlh-8>TIR1::F2A::2x mTurquoise2::PH::3xHA::tbb-2 3'UTR loxN ttTi4348] I*; *mpk-1(ljf38[mNG::3xFlag::AID::mpk-1]) III* | This paper | APL173 |
| *C. elegans*: *ljfSi3[Phlh-8>TIR1::F2A::2x mTurquoise2::PH::3xHA::tbb-2 3'UTR loxN ttTi4348] I*; *sos-1(ljf346[mNG::3xFlag::AID::sos-1]) V* | This paper | APL182 |
| *C. elegans*: *ljfSi30[Phlh-8>TIR1::F2A::2x mTurquoise2::PH::3xHA::tbb-2 3'UTR lox511i] III*; *egl-15(n1458) X* | This paper | APL197 |
| *C. elegans*: *ljfSi49 [Phlh-8>sos(cat):: 2x mTurquoise2::PH::3xHA::tbb-2 3'UTR loxN] I* | This paper | APL217 |
| *C. elegans*: *ljfSi51 [Phlh-8>let-60(G13E)::sl2::2x mTurquoise2::PH::tbb-2 3'UTR loxN] I* | This paper | APL236 |
| *C. elegans*: *ljfSi51 [Phlh-8>let-60(G13E)::sl2::2x mTurquoise2::PH::tbb-2 3'UTR loxN] I*; *egl-15(n1458) X* | This paper | APL237 |
| *C. elegans*: *ljfSi3[Phlh-8>TIR1::F2A::2x mTurquoise2::PH::3xHA::tbb-2 3'UTR loxN ttTi4348] I*; *sem-5(ljf35[sem-5::mScarlet::AID^lox511i SEC (mutated) lox511i]) X* | This paper | APL240 |
| *C. elegans*: *ljfSi10[Phlh-8>2x mTurquoise2::PH::tbb-2 3'UTR loxN ttTi5605] II*; *ljfSi32[Pegl-20(-1261 – 610)::pes-10>egl-17::mNG::SL2::2x mKate2::PH::tbb-2 3'UTR lox511i Chr IV]) IV*; *egl-17(ljf14[deletion + mNG^3xFlag]) X* | Gibney and Pani, 2025 | APL242 |
| *C. elegans*: *ljfSi3[Phlh-8>TIR1::F2A::2x mTurquoise2::PH::3xHA::tbb-2 3'UTR loxN ttTi4348] I*; *let-60(re302re359[AID::d10::let-60]) IV* | This paper | APL266 |
| *C. elegans*: *ljfSi23[Phlh-8>mScarlet-I::PH::3xHA::tbb-2 3'UTR loxN] I*; *egl-15(n1458) X* | This paper | APL281 |
| *C. elegans*: *ljfSi3[Phlh-8>TIR1::F2A::2x mTurquoise2::PH::3xHA::tbb-2 3'UTR loxN ttTi4348] I*; *abl-1(ljf46[abl-1::mNG::3xFlag::AID]) X* | This paper | APL294 |
| *C. elegans*: *ljfSi3[Phlh-8>TIR1::F2A::2x mTurquoise2::PH::3xHA::tbb-2 3'UTR loxN ttTi4348] I*; *let-60(re302re326[mScarlet-I[GLO]]::2xHA::3x linker::d10::let-60]) IV* | This paper | APL302 |
| *C. elegans*: *ljfSi3*[*Phlh-8*>*TIR1::SL2::2x mTurquoise2::PH::3xHA loxN*], *I*; *age-1*(*ljf11*[*age-1::mNG::3xFlag::AID*]), *II*; *mpk-1*(*ljf38*[*mNG*[*C1*]<sup>^</sup>*loxP*<sup>^</sup>*3xFlag::AID::mpk-1*]) *III* | This paper | APL306 |
| *C. elegans*: HML1035: *cshls128* [*rps-28p::TIR1::T2A::mCherry::his-11* + *Cbr-unc-119(+)*] *II*; *mpk-1*(*ljf39*[*mNG::3xFlag::AID::mpk-1*]) *III*; *ieSi58* [*eft-3p::degron::GFP::unc-54 3'UTR* + *Cbr-unc-119(+)*] *IV* | This paper | APL341 |
| *C. elegans*: *ljfSi3*[*Phlh-8*>*TIR1::F2A::2x mTurquoise2::PH::3xHA::tbb-2 3'UTR loxN ttTi4348*] *I*; *wsp-1*(*ljf49*[*wsp-1::mScarlet::AID*<sup>^</sup>*lox511*]) *IV* | This paper | APL357 |
| *C. elegans*: *ljfSi3*[*Phlh-8*>*TIR1::F2A::2x mTurquoise2::PH::3xHA::tbb-2 3'UTR loxN ttTi4348*] *I*; *ksr-1*(*ljf347*[*mNG::3xFlag::AID::ksr-1*]) *X* | This paper | APL410 |
| *C. elegans*: *ljfSi2*[*Phlh-8*>*mKate2::moesin::F2A::2x mTurquoise2::PH::3xHA::tbb-2 3'UTR loxN ttTi4348*] *I*; *plc-3*(*tm1340*)/*mIn1* [*mls14 dpy-10(e128)*] *II* | This paper | APL416 |
| *C. elegans*: *jfls10*[*Phlh-8*>*2xmTurquoise::PH::3xHA::tbb-2 3'UTR loxN ttTi5605*] *II*; *let-60* (*ljf52*[*let-60(G13E)*]) *IV* | This paper | APL430 |
| *C. elegans*: *ljfSi3*[*Phlh-8*>*TIR1::F2A::2x mTurquoise2::PH::3xHA::tbb-2 3'UTR loxN ttTi4348*] *I*; *arx-2*(*ljf45*[*arx-2::mNG::3xFlag::AID*]) *V* | This paper | APL452 |
| *C. elegans*: *ljfSi53* [*Phlh-8*>*sem-5::PH(internal)::mNG::tbb-2 3'UTR lox511i*] *Chr IV* | This paper | APL481 |
| *C. elegans*: *ljfSi3*[*Phlh-8*>*TIR1::F2A::2x mTurquoise2::PH::3xHA::tbb-2 3'UTR loxN ttTi4348*] *I*; *ced-10*(*ljf44*[*mScarlet::AID*<sup>^</sup>*lox511*<sup>^</sup>*ced-10*]) *IV*; *mig-2*(*ljf43*[*mNG::3xFlag::AID::mig-2*]) *X* | This paper | APL496 |
| *C. elegans*: *ljfSi10*[*Phlh-8*>*2xmTurquoise::PH::3xHA::tbb-2 3'UTR loxN ttTi5605*] *II*; *let-60* (*ljf53*[*let-60(G13E G75D)*]) *IV* | This paper | APL499 |
| *C. elegans*: *ljfSi2*[*Phlh-8*>*mKate2::Dme moesin ABD::F2A::2x mTurquoise2::PH::3xHA::tbb-2 3'UTR loxN ttTi4348*] *I*; *ljfSi50* [*Phlh-8*>*sos(cat)::2x mTurquoise2::PH::tbb-2 3'UTR lox511i*] *IV* | This paper | APL531 |
| *C. elegans*: *ljfSi3*[*Phlh-8*>*TIR1::SL2::2x mTurquoise2::PH::3xHA loxN*], *I*; *let-363*(*ljf41*[*mNG::3xFlag::AID::let-363*]) *I* | This paper | APL533 |
| *C. elegans*: JDW225: *eft-3p::TIR1::F2A::mTagBFP2::AID*<sup>*</sup>*::NLS::tbb-2 3'UTR* (*I:-5.32*); *age-1*(*ljf42*[*age-1::mNG::3xFlag::AID*]), *II* | This paper | APL556 |
| *C. elegans*: *ljfSi3*[*Phlh-8*>*TIR1::F2A::2x mTurquoise2::PH::3xHA::tbb-2 3'UTR loxN ttTi4348*] *I*; *wve-1*(*ljf48*[*wve-1::mNG::3xFlag::AID*]) *I*; *wsp-1*(*ljf49*[*wsp-1::mScarlet::AID*]) *IV* | This paper | APL564 |
| *C. elegans*: *ljfSi43*[*Phlh-8*>*TIR1::SL2::2x mTurquoise2::PH::3xHA::tbb-2 3'UTR loxN*], *I*; *akt-1*(*ljf40*[*mNG::3xFlag::AID::akt-1*]) *V*; *akt-2*(*ok393*) *X* | This paper | APL663 |
| *C. elegans*: *ljfSi44* [*Phlh-8*>*TIR1::SL2::2x mTurquoise2::PH::3xHA::tbb-2 3'UTR loxN*] *I*; *age-1*(*m333*)/*mnC1* [*dpy-10(e128) unc-52(e444)*] *II* | This paper | APL668 |
| *C. elegans*: *ljfSi46* [*Phlh-8*>*TIR1::SL2::2x mTurquoise2::PH::3xHA::tbb-2 3'UTR loxN*] *I*; *abl-1*(*ok171*) *X* | This paper | APL672 |
| *C. elegans*: *ljfSi27*[*Phlh-8*>*TIR1::F2A::2x mTurquoise2::PH::3xHA::tbb-2 3'UTR lox511i*] *III*; *ksr-1*(*n2526*) *X* | This paper | APL727 |
| *C. elegans*: *ljfSi3*[*Phlh-8*>*TIR1::F2A::2x mTurquoise2::PH::3xHA::tbb-2 3'UTR loxN ttTi4348*] *I*; *let-502*(*ljf47*[*mNG::3xFlag::AID::let-502*]) *I* | This paper | APL737 |
| *C. elegans*: *let-502*(*ljf47*[*mNG::3xFlag::AID::let-502*]) *I*; *ljfSi61* [*Peft-3*>*TIR1::tbb-2 3'UTR lox511i*] *IV* | This paper | APL738 |
| *C. elegans*: *ljfSi47* [*Phlh-8>TIR1::SL2::2x mTurquoise2::PH::3xHA::tbb-2 3'UTR loxN*] I; *ljfSi48* [*Pegl-20(-1261 – 610)::pes-10>egl-17::SL2::2x mKate2::PH::tbb-2 3'UTR lox511i*] IV; *sos-1*(*ljf51*[*mNG::3xFlag::AID::sos-1*]) V; *egl17*(*ljf14*[*egl-17 deletion + mNG^3xFlag*] X | This paper | APL771 |
| *C. elegans*: *ljfSi30*[*Phlh-8>TIR1::F2A::2x mTurquoise2::PH::3xHA::tbb-2 3'UTR lox511i*] III; *sos-1*(*ljf56*[*mNG::3xFlag::AID::sos-1*]) V; *egl-15*(*n1458*) X | This paper | APL779 |
| *C. elegans*: *ljfSi3*[*Phlh-8>TIR1::SL2::2x mTurquoise2::PH::3xHA loxN*] I; *age-1*(*ljf11*[*age-1::mNG::3xFlag::AID*]), II; *mpk-1*(*ljf38*[*mNG::3xFlag::AID::mpk-1*]) III; *ljfSi45* [*Peft-3>TIR1::tbb-2 3'UTR lox511i*] IV | This paper | APL785 |
| *C. elegans*: *ljfSi3*[*Phlh-8>TIR1::F2A::2x mTurquoise2::PH::3xHA::tbb-2 3'UTR loxN ttTi4348*] I; *wve-1*(*ljf48*[*wve-1::mNG::3xFlag::AID*]) I | This paper | APL826 |
| *C. elegans*: *ljfSi3*[*Phlh-8>TIR1::F2A::2x mTurquoise2::PH::3xHA::tbb-2 3'UTR loxN ttTi4348*] I; *mpk-1*(*ljf38*[*mNG::3xFlag::AID::mpk-1*]) III; *ljfSi54* [*Phlh-8>let-60(G13E)::sl2::2x mTurquoise2::PH::tbb-2 3'UTR lox511i*] Chr IV; *egl-15*(*ljf54*[*egl-15 5a-*]) X | This paper | APL885 |
| *C. elegans*: *ljfSi3*[*Phlh-8>TIR1::F2A::2x mTurquoise2::PH::3xHA::tbb-2 3'UTR loxN ttTi4348*] I; *bmdSi297*[*Prpl-28>FRT-stop-FRT tir1(F79G)::T2A::DHB::2x mKate2*] II; *ljfSi22* [*Phlh-8>flp-D5*] IV; *arx-2*(*ljf114*[*arx-2::mNG::3xFlag::AID*]) V | This paper, Xiao *et al.*, 2023 | APL974 |
| *C. elegans*: *bmdSi297*[*Prpl-28>FRT-stop-FRT tir1(F79G)::T2A::DHB::2x mKate2*] II; *mpk-1*(*ljf107*[*mNG::AID::mpk-1*]) III; *ljfSi22* [*Phlh-8>flp-D5*] IV | This paper, Xiao *et al.*, 2023 | APL976 |
| *C. elegans*: *ljfSi76* [*Phlh-8>let-60(G12V)::sl2::2x mTurquoise2::PH::3xHA::tbb-2 3'UTR lox511i*] IV | This paper | APL987 |
| *C. elegans*: *ljfSi3*[*Phlh-8>TIR1::F2A::2x mTurquoise2::PH::3xHA::tbb-2 3'UTR loxN ttTi4348*] I; *mpk-1*(*ljf38*[*mNG::3xFlag::AID::mpk-1*]) III; *ljfSi78* [*Phlh-8>let-60(G12V)::sl2::2x mTurquoise2::PH::3xHA::tbb-2 3'UTR lox511i*] IV | This paper | APL1020 |
| *C. elegans*: *ljfSi3*[*Phlh-8>TIR1::F2A::2x mTurquoise2::PH::3xHA::tbb-2 3'UTR loxN ttTi4348*] I; *unc-34*(*ljf3*[*unc-34::mNG::3xFlag::AID*]), V | This paper, Costa *et al.*, 2023 | APL1087 |
| *C. elegans*: *age-1*(*m333*)/*mnC1* [*dpy-10(e128) unc-52(e444)*] II | CGC | DR722 |
| *C. elegans*: *bmdSi297* [*loxN::rpl-28p::FRT3::STOP::FRT3::TIR1(F79G)::T2A::DHB::2xmKate2*] II | Xiao *et al.*, 2023 | DQM1113 |
| *C. elegans*: *let-60(re302[d10::let-60])* IV | This paper | DV3749 |
| *C. elegans*: *cshIs128* [*rps-28p::TIR1::T2A::mCherry::his-11 + Cbr-unc-119(+)*] II. *ieSi58* [*eft-3p::degron::GFP::unc-54 3'UTR + Cbr-unc-119(+)*] IV | CGC | HML1035 |
| *C. elegans*: *eft-3p::TIR1::F2A::mTagBFP2::AID::NLS::tbb-2 3'UTR* (I:-5.32) | CGC; Ashley *et al.*, 2021 | JDW225 |
| *C. elegans*: *ksr-1(n2526)* X | CGC | MT8677 |
| *C. elegans*: *plc-3(tm1340)/mln1* [*mls14 dpy-10(e128)*] II | CGC | PS4886 |
| *C. elegans*: *akt-2(ok393)* X | CGC | VC204 |
| *C. elegans*: *abl-1(ok171)* X | CGC | XR1 |
| **Recombinant DNA** |  |  |
| Plasmid: *Peft-3>Cas9 + PU6>Chr I (ttTi4348)* sgRNA | Pani & Goldstein, 2018 | pAP082 |
| Plasmid: *Phlh-8>2x mTurquoise2::PH::3xHA::tbb-2 3'UTR loxN SEC loxN, Chr II* | Gibney and Pani, 2025 | pAP138 |
| Plasmid: *PhlH-8>mScarlet-I[C1]_PH_3xHA_tbb-2 3'UTR ttTi4348 loxN SEC loxN* | This paper | pAP145 |
| Plasmid: *Peft-3>Cas9 + PU6>Chr II (ttTi5605) sgRNA* | Dickinson *et al.*, 2013 | pDD122 |
| Plasmid: *Peft-3>Cas9 + PU6>empty sgRNA for cloning* | Dickinson *et al.*, 2013 | pDD162 |
| Plasmid: *mTurquoise2[C1]^SEC^2xHA backbone for cloning homologous repair templates* | Dickinson *et al.*, 2013 | pDD315 |
| Plasmid: *Peft-3>Cas9 + PU6>egl-15 C-terminus sgRNA* | This paper | pTG008 |
| Plasmid: *egl-15::mNG^loxp::SEC::loxp^3xFlag::AID homologous repair template* | This paper | pTG010 |
| Plasmid: *PhlH-8>sos(cat)::2x mTurquoise2::PH::3xHA:tbb-2 3'UTR loxN SEC loxN, Chr I* | This paper | pTG109 |
| Plasmid: *Peft-3>Cas9 + PU6>Chr IV sgRNA* | Gibney and Pani, 2025 | pTG110 |
| Plasmid: *Peft-3>Cas9 + PU6>mpk-1 N-terminus sgRNA* | This paper | pTG141 |
| Plasmid: *mNG^loxp::SEC::loxp^3xFlag::AID::sos-1 homologous repair template* | This paper | pTG143 |
| Plasmid: *Peft-3>Cas9 + PU6>age-1 C-terminus sgRNA* | This paper | pTG146 |
| Plasmid: *Peft-3>Cas9 + PU6>sem-5 C-terminus sgRNA* | This paper | pTG149 |
| Plasmid: *age-1::mNG^loxp::SEC::loxp^3xFlag::AID homologous repair template* | This paper | pTG152 |
| Plasmid: *Peft-3>Cas9 + PU6>mpk-1 N-terminus sgRNA* | This paper | pTG173 |
| Plasmid: *mNG^loxp::SEC::loxp^3xFlag::AID::mpk-1 homologous repair template* | This paper | pTG174 |
| Plasmid: *PhlH-8>let-60(G13E)::sl2::2x mTurquoise2::PH::tbb-2 3'UTR loxN SEC loxN, Chr I* | This paper | pTG224 |
| Plasmid: *Peft-3>Cas9 + PU6>akt-1 N-terminus sgRNA* | This paper | pTG230 |
| Plasmid: *abl-1::mNG^loxp::SEC::loxp^3xFlag::AID homologous repair template* | This paper | pTG239 |
| Plasmid: *mNG^loxp::SEC::loxp^3xFlag::AID::akt-1 homologous repair template* | This paper | pTG247 |
| Plasmid: *Peft-3>Cas9 + PU6>abl-1 C-terminus sgRNA* | This paper | pTG274 |
| Plasmid: *Pegl-20(-1261 – 610)::pes-10>egl-17::mNG::SL2::2x mKate2::PH::tbb-2 3'UTR lox 511i SEC lox 511i, Chr IV* | Gibney and Pani, 2025 | pTG282 |
| Plasmid: *sem-5::mScarlet::AID^lox511I::SEC::lox511I^ homologous repair template* | This paper | pTG283 |
| Plasmid: *Peft-3>Cas9 + PU6>ced-10 N-terminus sgRNA* | This paper | pTG304 |
| Plasmid: *wsp-1::mScarlet::AID^lox511I::SEC::lox511I homologous repair template* | This paper | pTG312 |
| Plasmid: *Peft-3>Cas9 + PU6>wsp-1 C-terminus sgRNA* | This paper | pTG313 |
| Plasmid: *mNG^loxp::SEC::loxp^3xFlag::AID::let-502 homologous repair template* | This paper | pTG342 |
| Plasmid: *Peft-3>Cas9 + PU6>arx-2 C-terminus sgRNA* | This paper | pTG344 |
| Plasmid: *Peft-3>Cas9 + PU6>let-502 N-terminus sgRNA* | This paper | pTG349 |
| Plasmid: *mNG^loxp::SEC::loxp^3xFlag::AID::mig-2 homologous repair template* | This paper | pTG377 |
| Plasmid: *PhlH-8>let-60(G13E)::sl2::2x mTurquoise2::PH::tbb-2 3'UTR lox 511i SEC lox 511i, Chr IV* | This paper | pTG392 |
| Plasmid: *Peft-3>Cas9 + PU6>mig-2 N-terminus sgRNA* | This paper | pTG397 |
| Plasmid: *mScarlet::AID^lox511I::SEC::lox511I^ced-10* homologous repair template | This paper | pTG409 |
| Plasmid: *Peft-3>Cas9 + PU6>ksr-1* N-terminus sgRNA | This paper | pTG411 |
| Plasmid: *mNG^loxP::SEC::loxP^3xFlag::AID::ksr-1* homologous repair template | This paper | pTG413 |
| Plasmid: *Peft-3>Cas9 + PU6>wve-1* C-terminus sgRNA | This paper | pTG415 |
| Plasmid: *Phlh-8>sem-5::PH(internal)::mNG::tbb-2* 3'UTR *lox 511i SEC lox 511i, Chr IV* | This paper | pTG421 |
| Plasmid: *Pegl-20(-1261 – 610)::pes-10>egl-17::SL2::2x mKate2::PH::tbb-2* 3'UTR *lox 511i SEC lox 511i, Chr IV* | This paper | pTG422 |
| Plasmid: *Phlh-8>sos(cat)::2x mTurquoise2::PH::tbb-2* 3'UTR *lox511I SEC lox511I, ChrIV* | This paper | pTG427 |
| Plasmid: *arx-2::mNG^loxP::SEC::loxP^3xFlag::AID* homologous repair template | This paper | pTG429 |
| Plasmid: *Peft-3>Cas9 + PU6>let-363* N-terminus sgRNA | This paper | pTG445 |
| Plasmid: *wve-1::mNG^loxP::SEC::loxP^3xFlag::AID* homologous repair template | This paper | pTG452 |
| Plasmid: *mNG^loxP::SEC::loxP^3xFlag::AID::let-363* homologous repair template | This paper | pTG453 |
| Plasmid: *Peft-3>tir1::tbb-2* 3'UTR *lox511I SEC lox511I, ChrIV* | This paper | pTG565 |
| Plasmid: *mNG[C1]^SEC^3xFlag^AID* backbone for cloning homologous repair templates | Gift of Oliver Hobert lab | pUA77 |
| Plasmid: *Phlh-8>TIR1::F2A::2x mTurquoise2::PH::3xHA::tbb-2* 3'UTR *loxN SEC loxN, Chr I* | This paper | pWZ234 |
| **Software and algorithms** |  |  |
| ApE - A plasmid Editor (Version 2.0.61) | Davis & Jorgensen, 2022 |  |
| *C. elegans* codon adapter | Redemann *et al.*, 2011 |  |
| Fiji/ImageJ2 (Version: 2.14.0/1.54f) | Schindelin *et al.*, 2012 |  |
| Illustrator CC (Version 26.0.2) | Adobe Systems |  |
| Microsoft Excel (Version 16.89.1) | Microsoft |  |
| Nikon NIS-Elements (Versions AR 5.42.06 and AR 5.42.04) | Nikon Instruments Inc. |  |
| Prism (Version 10.5.0) | GraphPad Software |  |
| **Other** |  |  |
| NEBuilder HiFi DNA Assembly Cloning Kit | New England Biolabs | E5520S |
| PureLink HQ Mini Plasmid DNA Purification Kit | Invitrogen | K210001 |
| Q5 High-Fidelity 2X Master Mix | New England Biolabs | M0492L |
| Q5 Site-Directed Mutagenesis Kit | New England Biolabs | E0554S |

## Notes

### Competing Interest Statement

The authors have declared no competing interest.

### Summary of Updates

This version of the manuscript has been revised after peer review.

