## Supplemental Information for "FGF-dependent, polarized SOS activity orchestrates directed migration of *C. elegans* muscle progenitors independently of canonical effectors *in vivo*"

__**Document S1:**__

**Figures S1 – S12**

**Table S1. Source data for SM positioning.**

**Table S2. An intragenic revertant of let-60(n1046gf) causes strong SM migration defects**

**Supplemental File 1: Genome editing and molecular cloning information**

**Video S1 Legend**

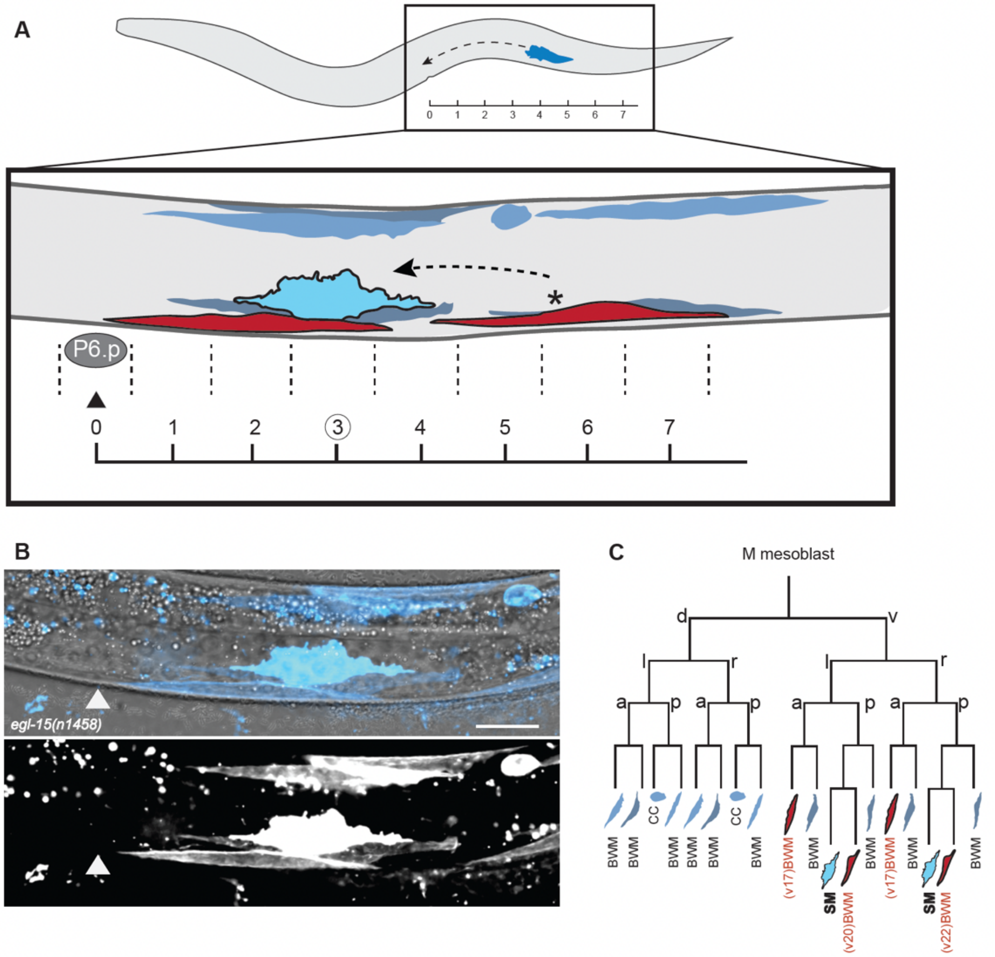


**Figure S1. Diagram of the methodology for scoring SM positions using M lineage-derived body wall muscles as reference points. (A)** Diagram of positions used for scoring SM positions relative to the M lineage-derived body wall muscle cells on the ventral side. Image shows average cell positioning in an *FGFR/egl-15(n1458)* mutant with impaired SM migration. For simplicity, only the left side of the worm is shown. Cells were scored based on where the cell body was positioned relative to the M-derived body wall muscle cells. We referred to the most anterior, M-derived, ventral body wall muscle cell as v17, whose anterior limit overlaps P6.p. For cells migrating posteriorly to category 4, the v20 or v22 cell was used as a reference (the most anterior of the two posterior ventral body wall cells on either the left or right respectively). The grey asterisk indicates the normal birthplace of SMs, with the curved dashed grey arrow indicating the distance the SMs normally travel even in the absence of FGF (EGL-17) or FGFR (EGL-15). Black triangle indicates normal endpoint of migration, above the vulval precursor cell (P6.p). **(B)** Example image of SM at a stopping position score of 3 in an *FGFR/egl-15(n1458)* background. White triangle indicates the normal endpoint of migration. **(C)** Diagram of M lineage divisions that from late L1 to the late L2 stage. CC= Coelomocyte, BWM = Body Wall Muscle, SM = Sex Myoblast, d= dorsal, v = ventral, l = left, r = right, a = anterior, p = posterior. BWM in red indicates cells that were used as references to score SM positions (v17, v20, v22). Scale bar = 10μm.

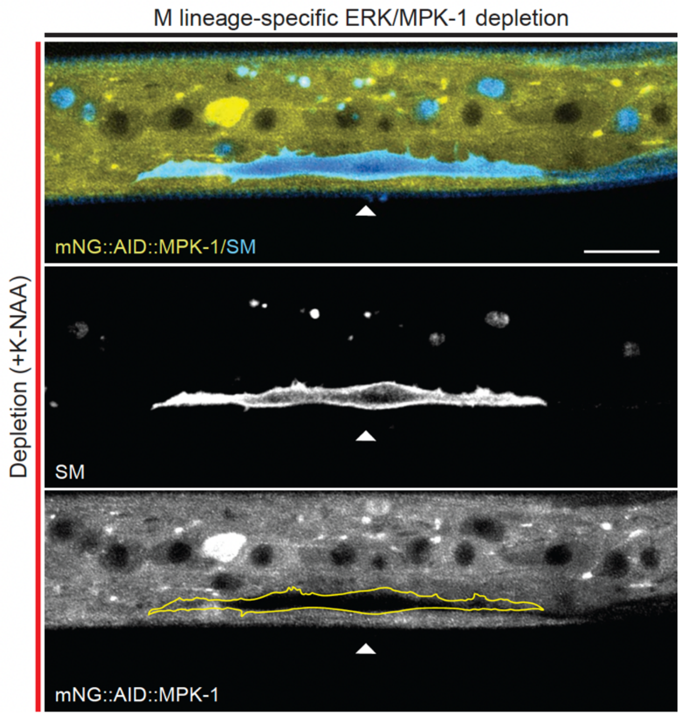


**Figure S2. Example of elongated SM morphology after cell-autonomous ERK/MPK-1 depletion.** Images show SM membrane and endogenously tagged mNG::AID::MPK-1 after K-NAA treatment. Note the SM has migrated to the correct position, but is unusually long and thin. White triangles indicate the normal endpoint of SM migration. Circular objects visible in both fluorescence channels are autofluorescent gut granules. All images are oriented with anterior to left and dorsal to top. Scale bars = 10μm.

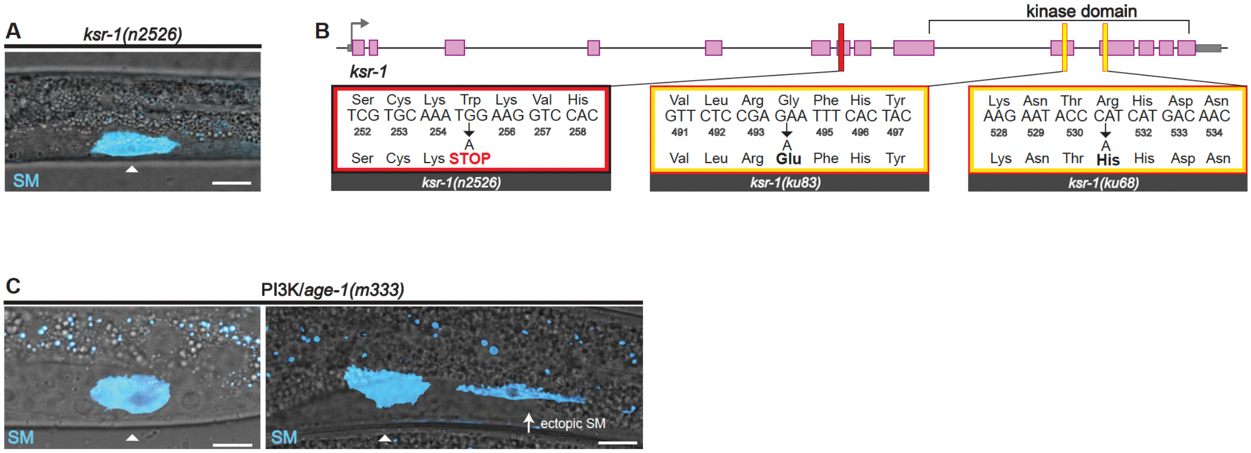


**Figure S3. SM positioning in *ksr-1(n2526)* and *age-1(m333)* mutant animals.** (**A**) Representative image of normally positioned SM in a *ksr-1(n2526)* animal. 60/60 SMs migrated to the correct position in this strain. (**B**) Diagram of the *ksr-1* genomic locus and the *n2526, ku83,* and *ku68* mutations. (**C**) SM positioning in *age-1(m333)* animals with normal M lineage differentiation (left) or an ectopic SM (right, arrow). White triangles indicate the normal endpoint of SM migration. Circular objects visible in visible in the membrane fluorescence channel are autofluorescent gut granules. All images are oriented with anterior to left and dorsal to top. Scale bars = 10µm.

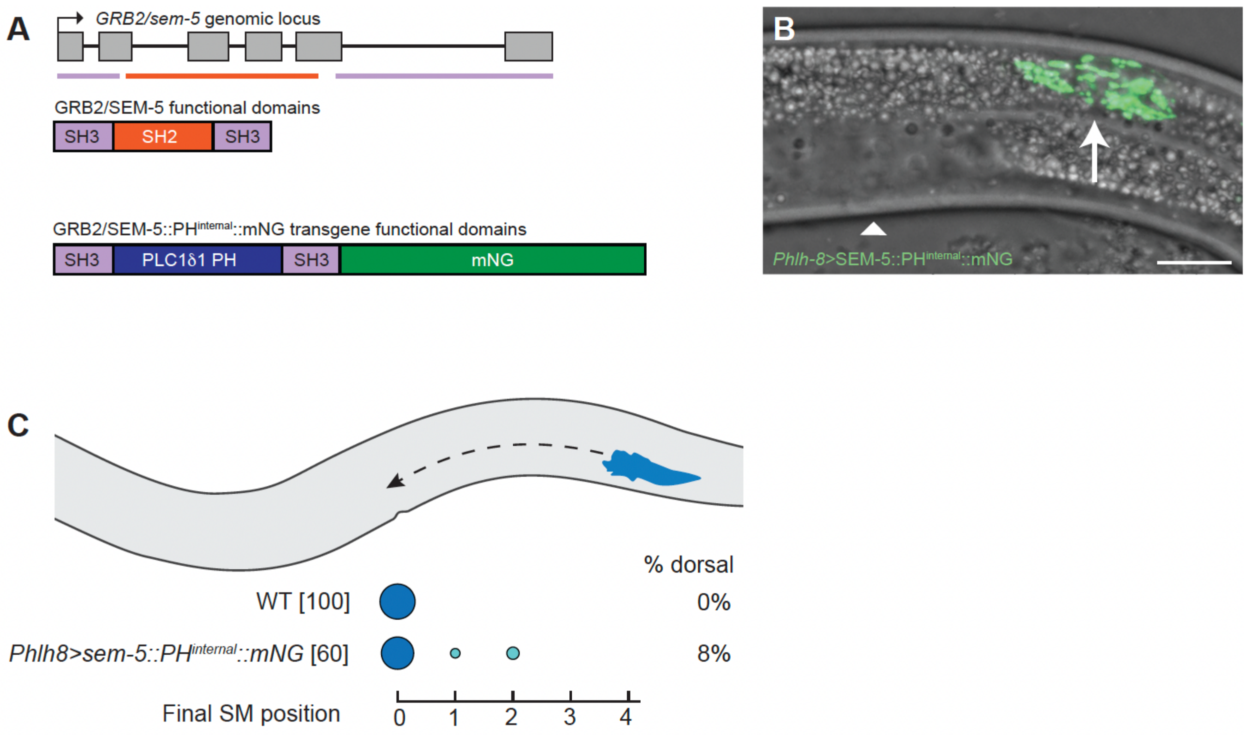


**Figure S4. Membrane-associated GRB2/SEM-5 activity disrupts SM migration. (A)** Schematic of the *GRB2/sem-5* locus, functional protein domains, and the engineered GRB2/SEM-5::PH::mNG construct. To promote constitutive, uniform activity, we replaced the SH2 domain the PH domain from *plc1δ1*. **(B)** Live spinning disk confocal image showing SEM-5::PH$^{internal}$::mNG localization. Note the non-uniform localization of the fusion protein and prominent internalized signal, which differs from the SOS$^{cat}$::2x mT2::PH fusion. White triangle indicates the normal endpoint of migration. Images is oriented with anterior to left and dorsal to top. Scale bar = 10µm. **(C)** Dot plot summarizing SM migration endpoints. Area of each dot is proportional to the frequency of cells stopping at that position. Number of SMs analyzed for each strain are shown in brackets. Source data available in Table S1.

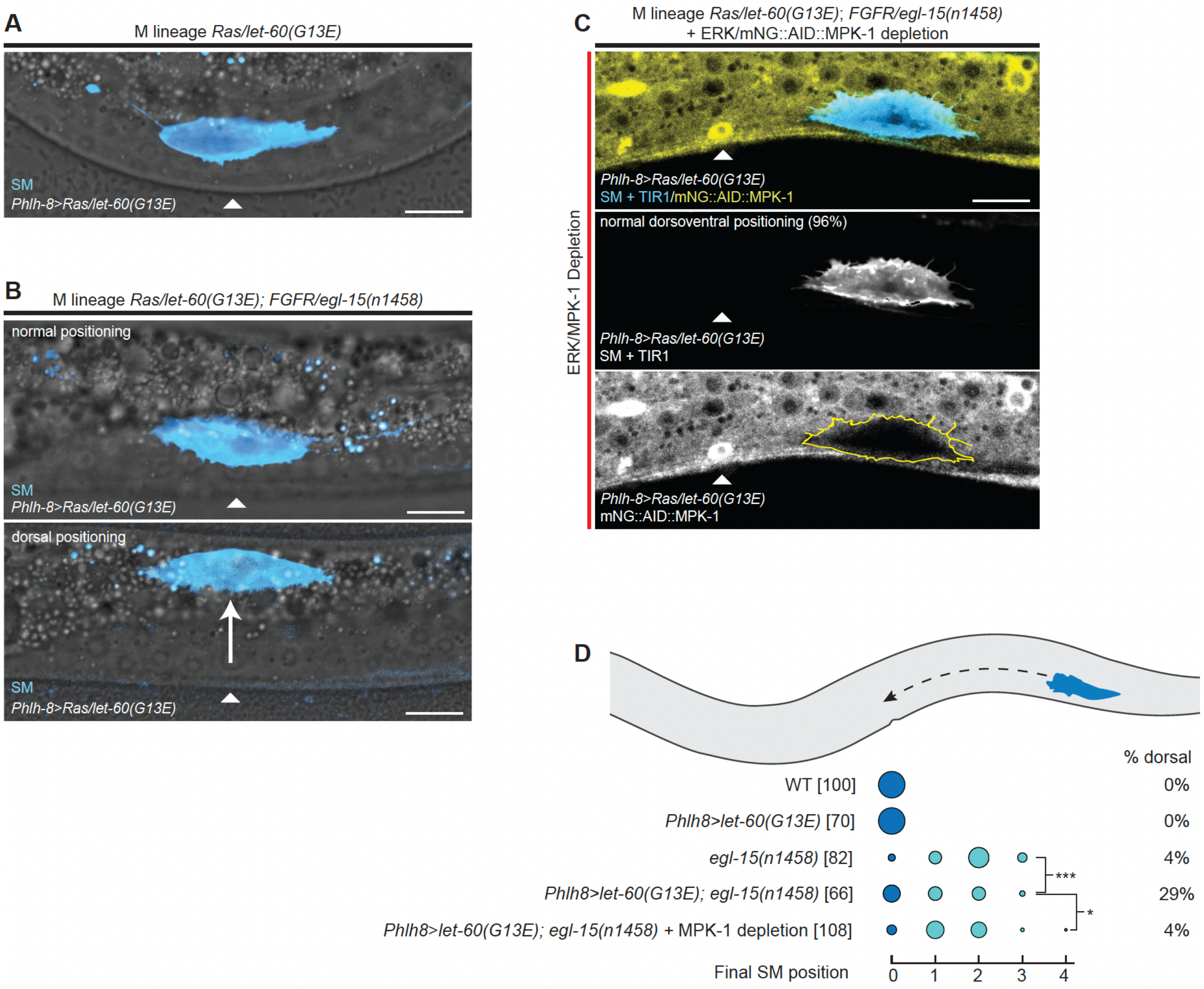


**Figure S5. ERK/MPK-1 depletion suppresses dorsal and enhances anterior migration defects in a *Phlh8>Ras/let-60(G13E); FGFR/egl-15(n1458)* background.** (**A**) Live spinning disk confocal image showing SM positioning in an animal with activated *Ras/let-60(G13E)* expressed in the M lineage. **(B)** Examples of *Phlh8>Ras/let-60(G13E); FGFR/egl-15(n1458)* animals with normally positioned SM (top image) or dorsally misplaced SM (bottom image). **(C)** SM positioning and mNG::AID::ERK/MPK-1 fluorescence in a *Phlh8>Ras/let-60(G13E); FGFR/egl-15(n1458)* animal with mNG::AID::ERK/MPK-1 depleted in the M lineage. **(D)** Dot plot summarizing SM migration endpoints. Area of each dot is proportional to the frequency of cells stopping at that position. Number of SMs analyzed for each strain are shown in brackets. Asterisks indicate statistical significance for indicated comparisons (Fisher's exact test, *, $P<0.05$; ***, $P<0.0005$). Source data available in Table S1. White triangles indicate the normal endpoint of migration. All images are oriented with anterior to left and dorsal to top. Scale bars = 10µm.

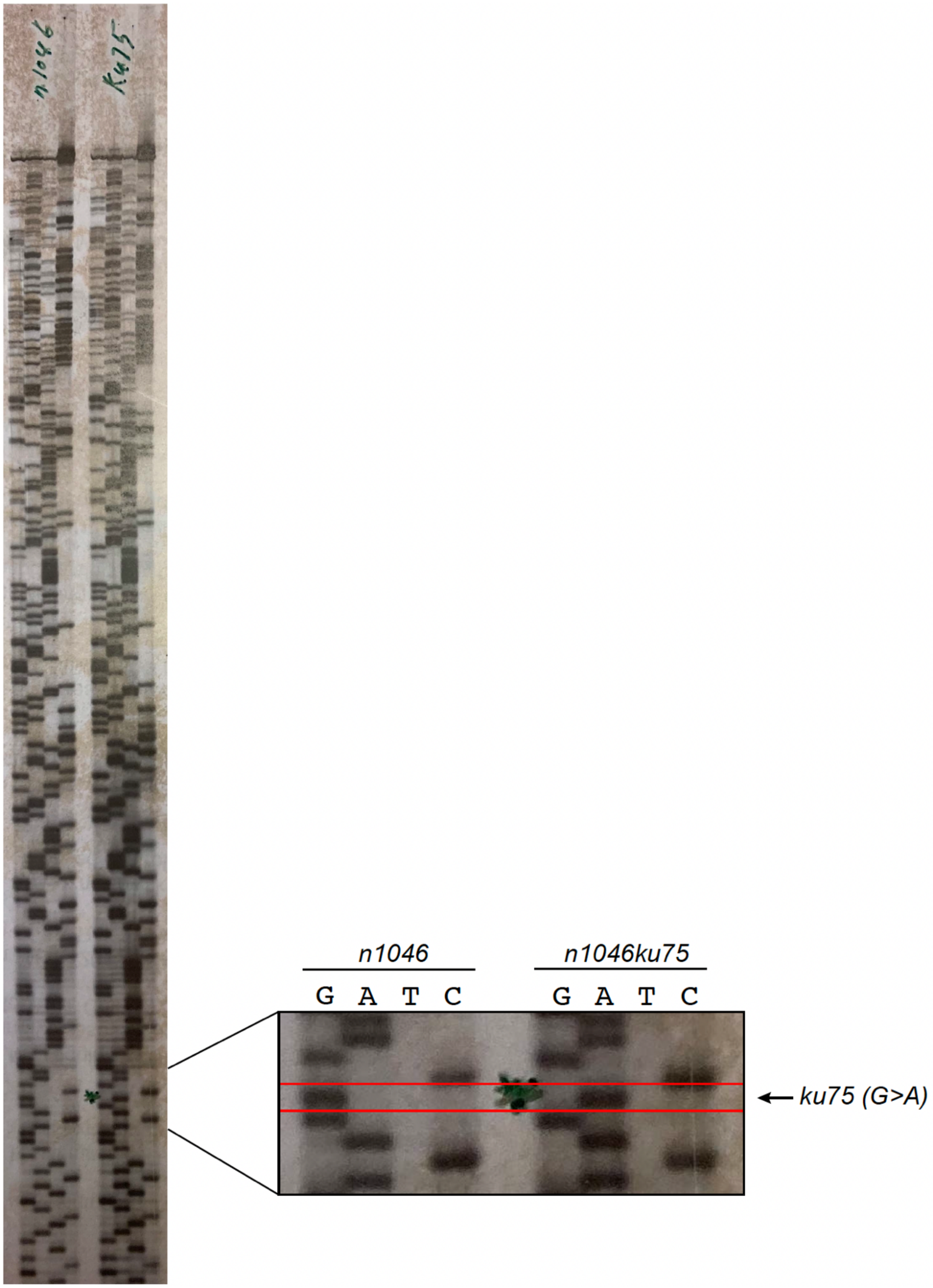


**Figure S6. Identification of the molecular lesion responsible for *let-60(n1046ku75)*.** Sanger sequencing gel showing DNA sequences of *let-60* PCR products in *let-60(n1046)* and *let-60(n1046ku75)* animals. *ku75* is G>A nucleotide substitution that leads to a glycine to aspartic acid substitution at amino acid position 75.

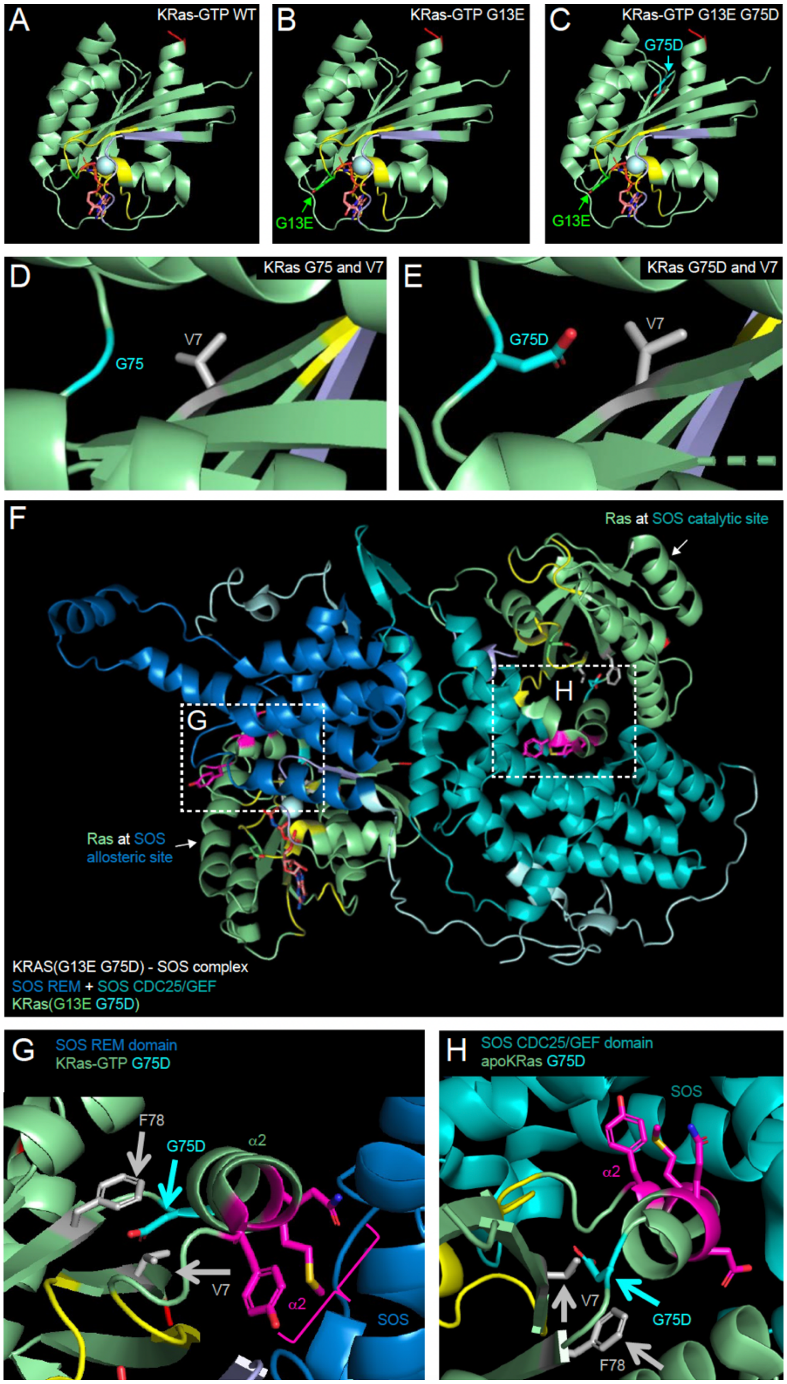


**Figure S7: Ras/LET-60(G75D) substitution is predicted to perturb interactions between Ras/LET-60 and SOS-1.** Ras/LET-60(G75D) is predicted to displace the Ras/LET-60 α helix 2 that interacts with SOS proteins, thus perturbing predicted interactions. **A**-**E**) PyMOL visualization of a crystal structure (PDB: 8EBZ) of human KRas G13D with GMPPNP non-hydrolyzable GTP analog (without the C-terminal hypervariable (HVR)+CAAX membrane-targeting sequences). Color coding. Small GTPase = pale green, nucleotide = salmon, basic charges = dark blue, acidic charges = red, $Mg^{2+}$ = pale cyan, G1-5 boxes that coordinate nucleotide binding = yellow, core effector binding loop = lavender, residue 13 = bright green, residue 75 = lt. blue, residue 7 = gray, last residue of the GTPase domain = red and is oriented upwards where it would normally extend to the HVR+CAAX sequences and the lipid bilayer. (**A**, **B**) Without the R-group to illustrate Gly13 vs. G13E. The structure (PDB: 8EBZ) is for KRas G13D, but Glutamate13 was substituted for Aspartate here to illustrate the *C. elegans let-60(n1046*gf[G13E]*)* mutant. The Asp13 changes causes mild constitutive loading of KRas, and Glu13 is expected to be similar. (**C**) Position of the G75D mutant change caused by the *Ras/let-60(n1046 ku75 [G13E G75D])* intragenic revertant mutation. (D-E) Zoomed in views contrasting wild-type Gly75 and mutant G75D. (**D**) The single H+ atom of the Gly75 side chain (not depicted) does not conflict with Val7, shown in gray (F78 not highlighted). (**E**) The larger and acidic G75D side chain is predicted to conflict with V7 in the hydrophobic core. This and steric interference are expected to displace the neighboring $\alpha$ helix 2. (**F-H**) PyMOL visualization of predicted interactions between KRas(G13E G75D) and SOS based on the crystal structure of a human ternary KRas(G13D)-SOS complex (PDB: 7KFZ). Color coding: Human SOS1: REM = blue, GEF = teal, other sequences pale cyan. KRas(G13D): GTPase = pale green, GTP = salmon, $Mg^{2+}$ = pale cyan, G boxes coordinating nucleotide binding = yellow, basic charges = dark blue, acidic charges = red, N- and C-terminal residues = red, alpha helix 2 residues oriented toward SOS = pink. (**F**) View of two KRas molecules bound to the SOS allosteric and catalytic sites. Dashed boxes show locations of zoomed views in G and H. (**G**) Zoomed view of predicted interaction between KRas-GTP(G75D) and the REM domain of SOS1 at its allosteric Ras-binding site. Gly75, altered by the G75D mutation, is shown as mutant Aspartate that interferes with Val7 and Phe78 with side chain. Dark blue = basic, red = acidic, yellow = sulfur. (**H**) Zoomed view of predicted interaction between apoKRas(G75) and the CDC25/GEF domain of SOS1 at its catalytic site.

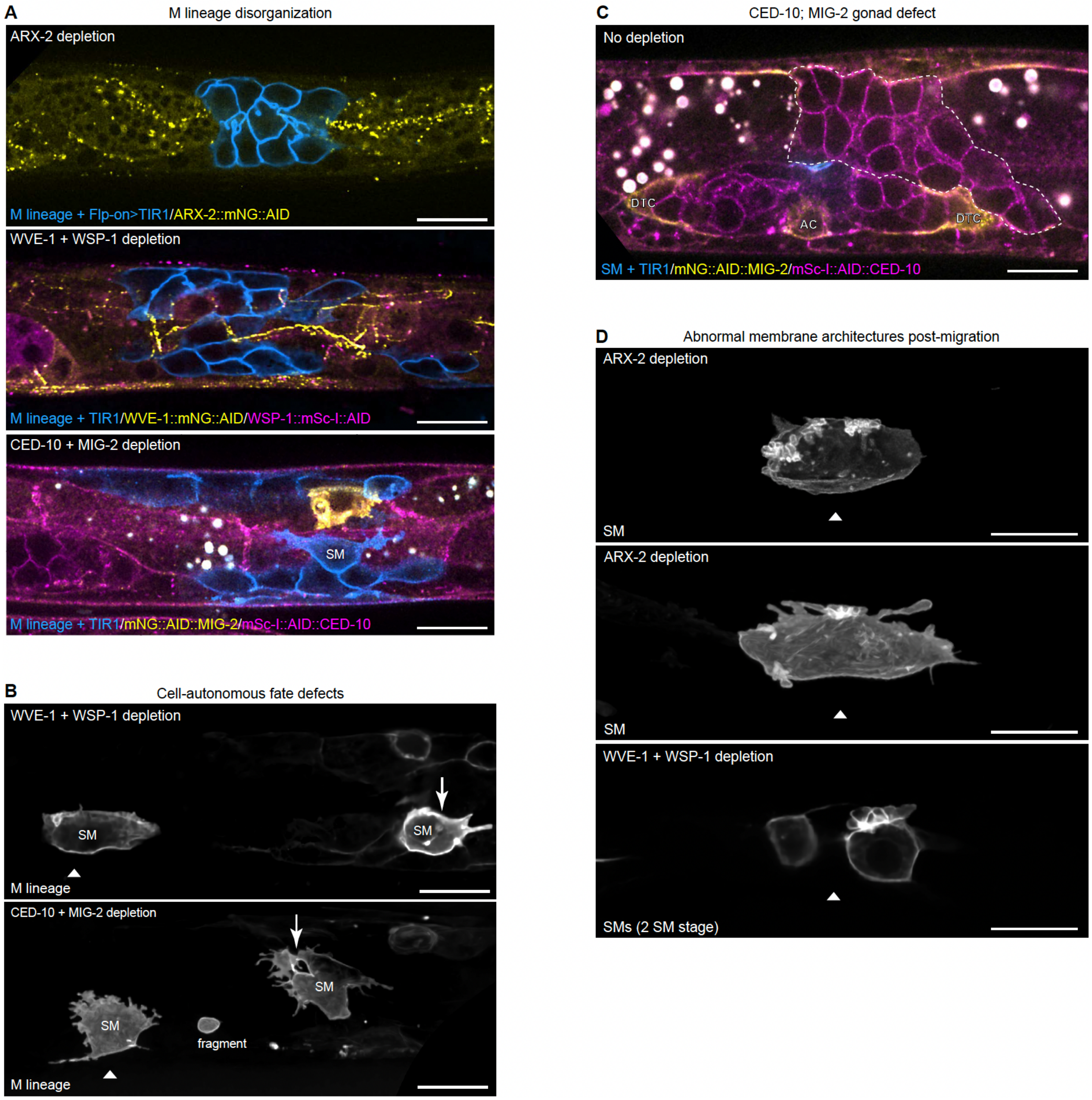


**Figure S8. Examples of M lineage defects caused by cytoskeletal regulator depletion, gonad defects caused by CED-10 tagging, and abnormal post-migratory SM membrane architectures after ARX-2 or WVE-1 plus WSP-1 depletion.** (**A**) Examples of defects in early M lineage development caused by depleting ARX-2, WVE-1 plus WSP-1, or CED-10 plus MIG-2 continuously in the M lineage. Animals expressing TIR1(F79) were raised from hatching on 5-Ph-IAA-treated plates. Animals expressing unmodified TIR1 were raised on K-NAA-treated plates. (**B**) Examples of ectopic SMs, which likely represent defects in M lineage cell fates, caused by continuous depletion of WVE-1 plus WSP-1 or CED-10 plus MIG-2. (**C**) Example of developmental defects in a mScarlet-I::AID::CED-10; mNG::AID::MIG-2 animal. Abnormally located gonad cells are indicated with a dashed outline. Abbreviations: AC, anchor cell; DTC, distal tip cell. (**D**) Examples of abnormal membrane morphologies in post-migratory SMs caused by depleting ARX-2 or WVE-1 plus WSP-1. White triangles indicate the normal endpoint of SM migration. All images are oriented with anterior to left and dorsal to top. Scale bars = 10μm.

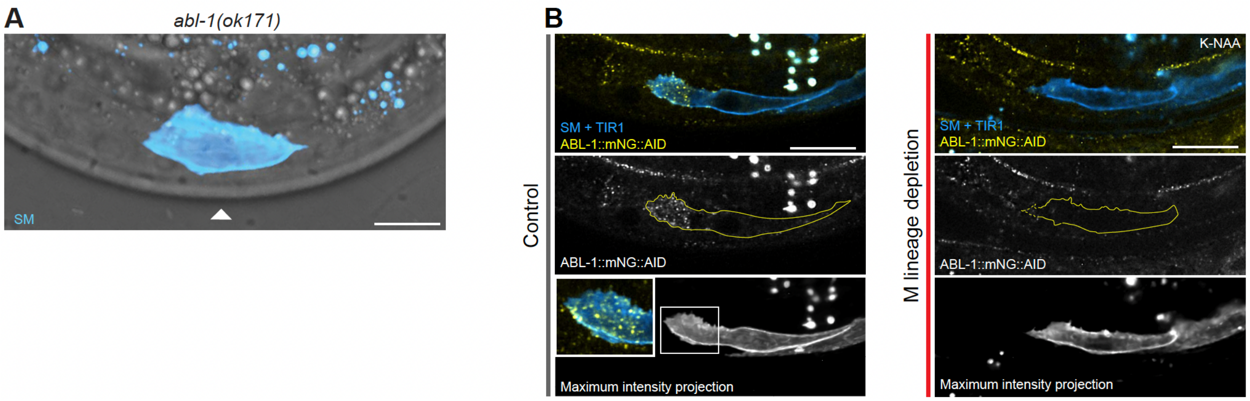


**Figure S9. ABL-1 is not required for normal SM positioning or lamellar protrusions during migration.** (**A**) Live image of representative SM positioning in an *abl-1(ok171)* mutant animal (66/66 SMs normally positioned). (**B**) Live, spinning disk confocal images showing membrane morphology in migrating SMs and tagged endogenous ABL-1::mNG::AID localization in control (left) and K-NAA-treated (right) animals. Images show single z-slices with the leading edge in focus for the membrane and endogenously tagged protein channels along with a maximum intensity projection to depict cellular morphology. SMs are outlined in images where the SM membrane marker channel (blue) is not shown. Dashed sections of the outline indicate regions where cells above or below the SM are visible in the same focal plane or where the SM membrane is not in focus. White triangle indicate the normal endpoint of SM migration in A. All images are oriented with anterior to left and dorsal to top. Scale bars = 10μm.

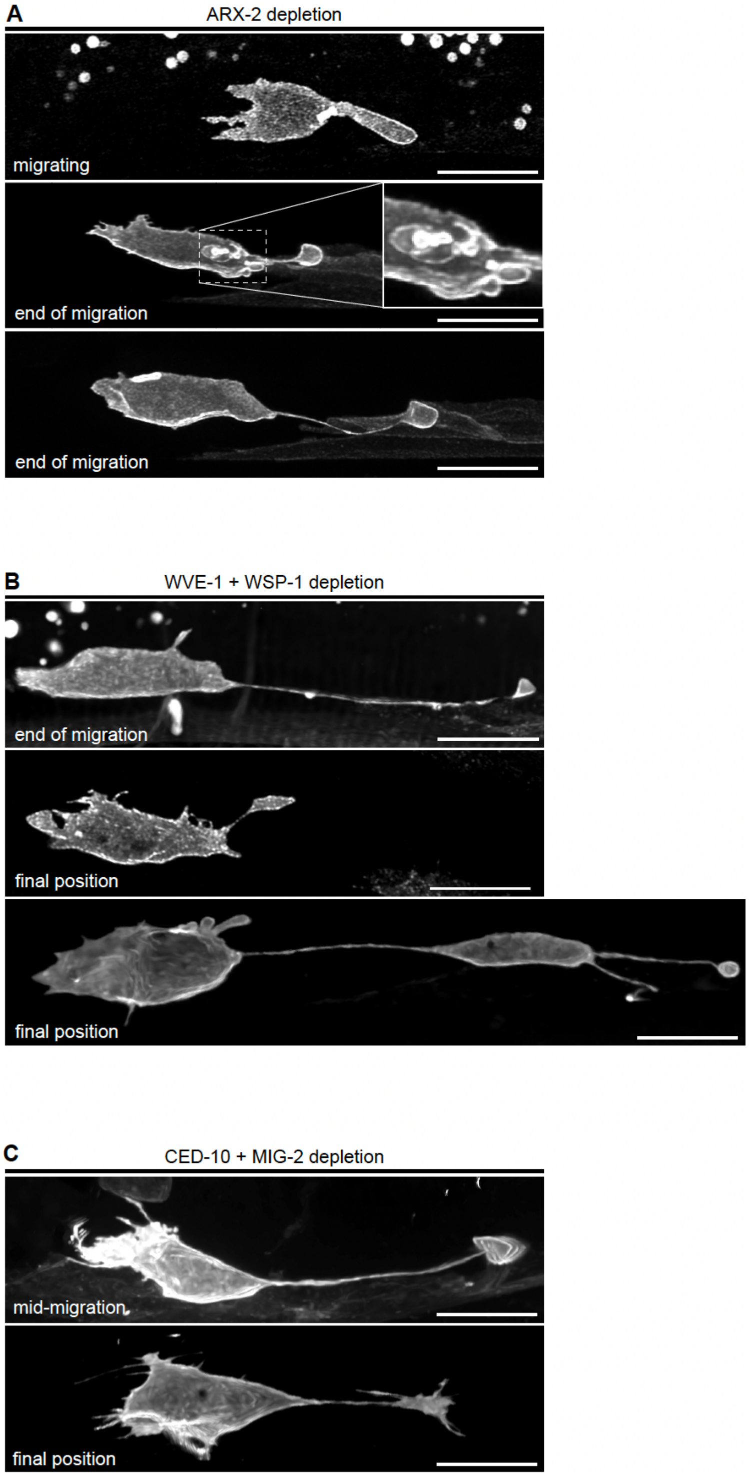


**Figure S10. Examples of abnormal SM trailing edge morphologies caused by depleting cytoskeletal regulators in the M lineage.** Maximum intensity projections of plasma membrane markers showing examples of abnormal trailing edge structures observed after depleting ARX-2 (**A**), WVE-1 plus WSP-1 (**B**) or CED-10 + MIG-2 (**C**). Note the presence of a thin filament connected to a trailing patch of membrane after depleting multiple actin regulators. Images are oriented with anterior to left and dorsal to top. Scale bars = 10μm.

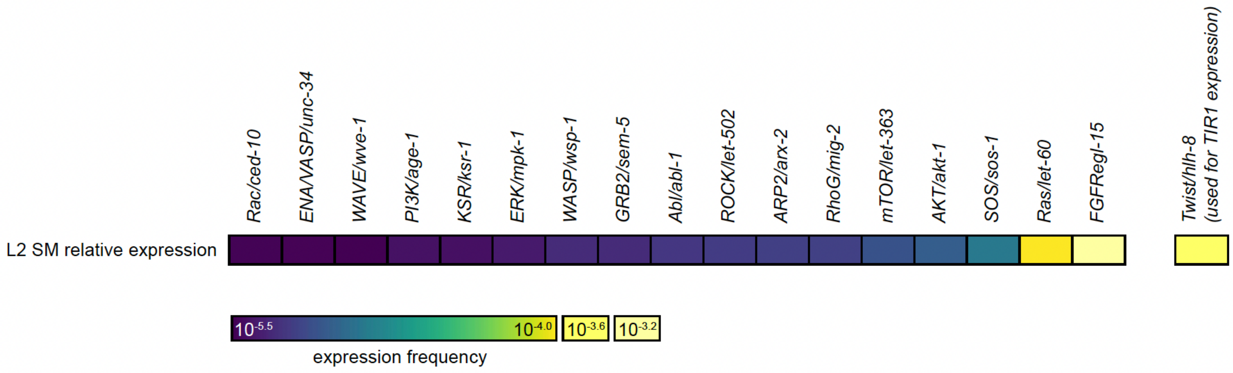


**Figure S11. Relative expression of targeted candidates genes in SMs in L2 scRNAseq data. (A)** Relative gene expression frequency[44] for the seventeen genes targeted by auxin-inducible degradation and the *hlh-8* gene whose promoter was used for transgenic TIR1 expression in most experiments. Note that targeting the three highest-expressed genes resulting in SM migration defects.

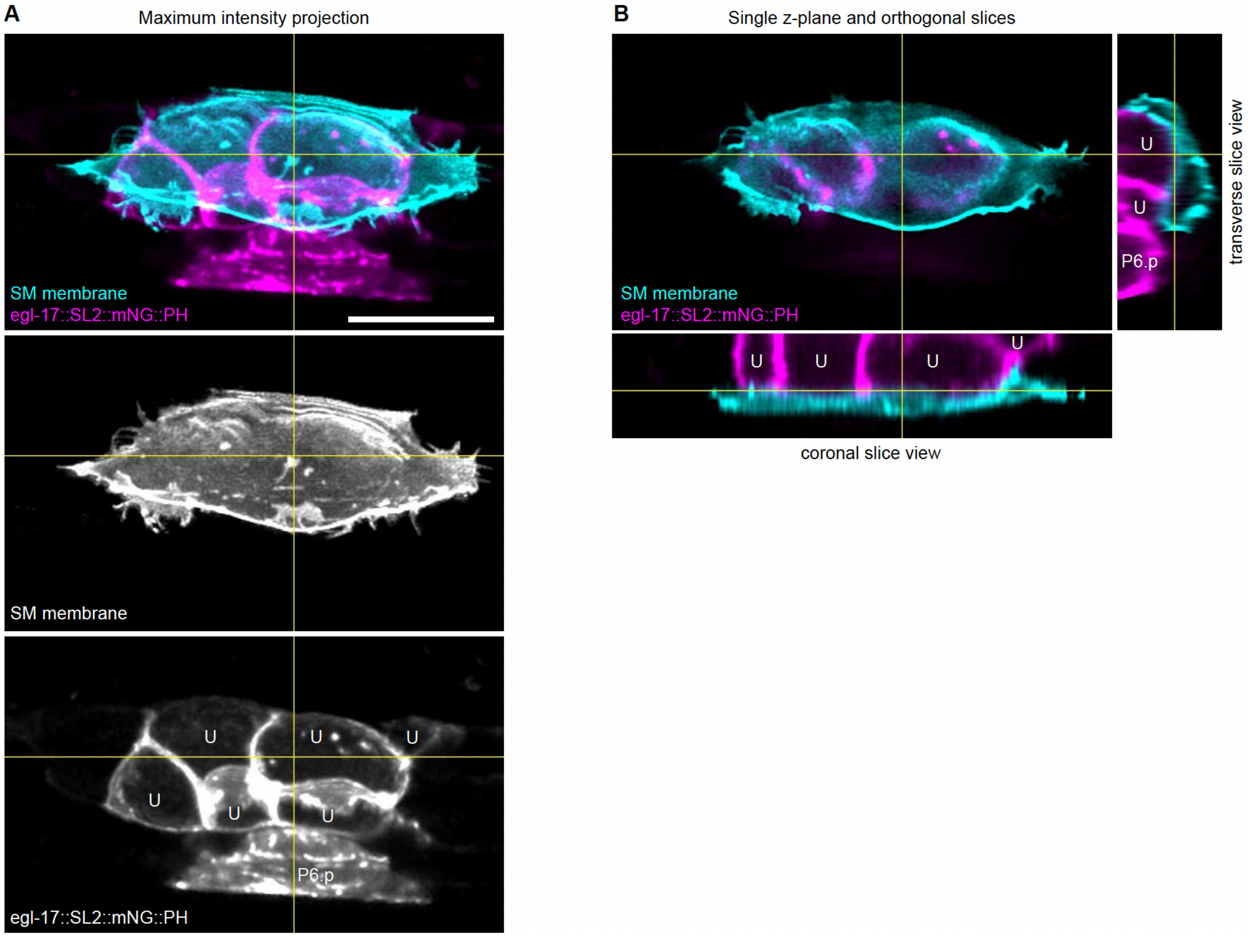


**Figure S12. Fluorescence from underlying uterine cells is visible in the same confocal z-plane as the SM.** Maximum intensity projection (**A**) and single z-plane with orthogonal slices (**B**) showing SM plasma membrane (cyan) and plasma membranes of underlying uterine cells (U) and P6.p. The SM was visualized with Phlh-8>2x mT2::PH, and underlying cells were visualized using egl-17::SL2::mNG::PH. Note that magenta fluorescence from uterine cells in **B** is visible in the same plane as the center of the SM. Animal is oriented with anterior to left and dorsal to top. Scale bar = 10μm.

**Video S1. Ras/LET-60(G75D) substitution is predicted to perturb interactions between Ras/LET-60 and SOS-1.** Video shows rotated views of predicted interactions between KRas(G13E G75D) and SOS based on the crystal structure of a human ternary KRas(G13D)-SOS complex (PDB: 7KFZ). See Figure S7 for annotations.

**Supplemental Table 1: Source data for final SM positioning**

| | | | AP position | | | | | | | | | | | | | |
|---|---|---|---|---|---|---|---|---|---|---|---|---|---|---|---|---|
| | | | Overmigration | | | | Normal | undermigration | | | | | | | Average | |
| Strain | Treatment | n | -4 | -3 | -2 | -1 | 0 | 1 | 2 | 3 | 4 | 5 | 6 | 7 | position | % Dorsal |
| WT (N2) | KNAA | 62 | 0.0% | 0.0% | 0.0% | 0.0% | 100.0% | 0.0% | 0.0% | 0.0% | 0.0% | 0.0% | 0.0% | 0.0% | 0.0 | 0.0% |
| | Control | 100 | 0.0% | 0.0% | 0.0% | 0.0% | 100.0% | 0.0% | 0.0% | 0.0% | 0.0% | 0.0% | 0.0% | 0.0% | 0.0 | 0.0% |
| *egl-15(n1458)* | KNAA | 76 | 0.0% | 0.0% | 0.0% | 0.0% | 6.6% | 22.4% | 57.9% | 11.8% | 1.3% | 0.0% | 0.0% | 0.0% | 1.8 | 5.3% |
| | Control | 82 | 0.0% | 0.0% | 0.0% | 0.0% | 3.7% | 22.0% | 54.9% | 19.5% | 0.0% | 0.0% | 0.0% | 0.0% | 1.9 | 3.7% |
| *Phlh-8>TIR-1::F2A::2x mT2::PH (N2 background)* | KNAA | 74 | 0.0% | 0.0% | 0.0% | 0.0% | 100.0% | 0.0% | 0.0% | 0.0% | 0.0% | 0.0% | 0.0% | 0.0% | 0.0 | 0.0% |
| | Control | 60 | 0.0% | 0.0% | 0.0% | 0.0% | 100.0% | 0.0% | 0.0% | 0.0% | 0.0% | 0.0% | 0.0% | 0.0% | 0.0 | 0.0% |
| *egl-15::mNG::AID; Phlh-8>TIR-1::F2A::2x mT2::PH* | KNAA | 100 | 0.0% | 0.0% | 0.0% | 0.0% | 3.0% | 32.0% | 48.0% | 17.0% | 0.0% | 0.0% | 0.0% | 0.0% | 1.8 | 3.0% |
| | Control | 90 | 0.0% | 0.0% | 0.0% | 0.0% | 100.0% | 0.0% | 0.0% | 0.0% | 0.0% | 0.0% | 0.0% | 0.0% | 0.0 | 0.0% |
| *sem-5::mScarlet-I::AID; Phlh-8>TIR-1::F2A::2x mT2::PH* | KNAA | 76 | 0.0% | 0.0% | 0.0% | 0.0% | 7.9% | 30.3% | 35.5% | 25.0% | 1.3% | 0.0% | 0.0% | 0.0% | 1.8 | 2.6% |
| | Control | 60 | 0.0% | 0.0% | 0.0% | 0.0% | 100.0% | 0.0% | 0.0% | 0.0% | 0.0% | 0.0% | 0.0% | 0.0% | 0.0 | 0.0% |
| *mNG::AID::sos-1; Phlh-8>TIR-1::F2A::2x mT2::PH* | KNAA | 74 | 0.0% | 0.0% | 0.0% | 0.0% | 9.5% | 44.6% | 36.5% | 9.5% | 0.0% | 0.0% | 0.0% | 0.0% | 1.5 | 8.1% |
| | Control | 80 | 0.0% | 0.0% | 0.0% | 0.0% | 100.0% | 0.0% | 0.0% | 0.0% | 0.0% | 0.0% | 0.0% | 0.0% | 0.0 | 0.0% |
| *AID::let-60; Phlh-8>TIR-1::F2A::2x mT2::PH* | KNAA | 90 | 0.0% | 0.0% | 0.0% | 0.0% | 22.2% | 45.6% | 21.1% | 11.1% | 0.0% | 0.0% | 0.0% | 0.0% | 1.2 | 2.2% |
| | Control | 60 | 0.0% | 0.0% | 0.0% | 0.0% | 100.0% | 0.0% | 0.0% | 0.0% | 0.0% | 0.0% | 0.0% | 0.0% | 0.0 | 0.0% |
| *mNG::AID::ksr-1; Phlh-8>TIR-1::F2A::2x mT2::PH* | KNAA | 72 | 0.0% | 0.0% | 0.0% | 0.0% | 0.0% | 0.0% | 0.0% | 0.0% | 0.0% | 0.0% | 0.0% | 0.0% | 0.0 | 0.0% |
| | Control | 60 | 0.0% | 0.0% | 0.0% | 0.0% | 100.0% | 0.0% | 0.0% | 0.0% | 0.0% | 0.0% | 0.0% | 0.0% | 0.0 | 0.0% |
| *ksr-1(n25261 [60])* | None | 60 | 0.0% | 0.0% | 0.0% | 0.0% | 100.0% | 0.0% | 0.0% | 0.0% | 0.0% | 0.0% | 0.0% | 0.0% | 0.0 | 0.0% |
| *mNG::AID:::mpk-1; Phlh-8>TIR-1::F2A::2x mT2::PH* | KNAA | 66 | 0.0% | 0.0% | 0.0% | 0.0% | 100.0% | 0.0% | 0.0% | 0.0% | 0.0% | 0.0% | 0.0% | 0.0% | 0.0 | 0.0% |
| | Control | 60 | 0.0% | 0.0% | 0.0% | 0.0% | 100.0% | 0.0% | 0.0% | 0.0% | 0.0% | 0.0% | 0.0% | 0.0% | 0.0 | 0.0% |
| *mNG::AID:::mpk-1; Prpl-28>TIR-1::P2A::mCherry::his-11* | KNAA | 64 | 0.0% | 0.0% | 0.0% | 16.0% | 39.0% | 25.0% | 20.0% | 0.0% | 0.0% | 0.0% | 0.0% | 0.0% | 0.5 | 3.0% |
| | KNAA - Timed* | 84 | 0.0% | 0.0% | 0.0% | 2.0% | 94.0% | 4.0% | 0.0% | 0.0% | 0.0% | 0.0% | 0.0% | 0.0% | 0.0 | 0.0% |
| | Control | 60 | 0.0% | 0.0% | 0.0% | 0.0% | 100.0% | 0.0% | 0.0% | 0.0% | 0.0% | 0.0% | 0.0% | 0.0% | 0.0 | 0.0% |
| *mNG::AID:::mpk-1; Prpl-28>FLP-ON>TIR1(F79G)::T2A::2x mKate2::DHB; Phlh-8>flp-D5* | 5-Ph-IAA | 80 | 0.0% | 0.0% | 0.0% | 0.0% | 100.0% | 0.0% | 0.0% | 0.0% | 0.0% | 0.0% | 0.0% | 0.0% | 0.0 | 0.0% |
| | Control | 60 | 0.0% | 0.0% | 0.0% | 0.0% | 100.0% | 0.0% | 0.0% | 0.0% | 0.0% | 0.0% | 0.0% | 0.0% | 0.0 | 0.0% |
| *age-1::mNG::AID; Phlh-8>TIR-1::F2A::2x mT2::PH* | KNAA | 70 | 0.0% | 0.0% | 0.0% | 0.0% | 100.0% | 0.0% | 0.0% | 0.0% | 0.0% | 0.0% | 0.0% | 0.0% | 0.0 | 0.0% |
| | Control | 66 | 0.0% | 0.0% | 0.0% | 0.0% | 100.0% | 0.0% | 0.0% | 0.0% | 0.0% | 0.0% | 0.0% | 0.0% | 0.0 | 0.0% |
| *age-1(m333)*** | None | 64 | 0.0% | 0.0% | 0.0% | 0.0% | 100.0% | 0.0% | 0.0% | 0.0% | 0.0% | 0.0% | 0.0% | 0.0% | 0.0 | 0.0% |
| *akt-1::mNG::AID; akt-2(ok393); Phlh-8>TIR-1::F2A::2x mT2::PH* | KNAA | 66 | 0.0% | 0.0% | 0.0% | 0.0% | 100.0% | 0.0% | 0.0% | 0.0% | 0.0% | 0.0% | 0.0% | 0.0% | 0.0 | 0.0% |
| | Control | 68 | 0.0% | 0.0% | 0.0% | 0.0% | 100.0% | 0.0% | 0.0% | 0.0% | 0.0% | 0.0% | 0.0% | 0.0% | 0.0 | 0.0% |
| *mNG::AID::let-363; Phlh-8>TIR-1::F2A::2x mT2::PH**** | KNAA - Timed* | 62 | 0.0% | 0.0% | 0.0% | 0.0% | 96.8% | 3.2% | 0.0% | 0.0% | 0.0% | 0.0% | 0.0% | 0.0% | 0.0 | 0.0% |
| | Control | 80 | 0.0% | 0.0% | 0.0% | 0.0% | 100.0% | 0.0% | 0.0% | 0.0% | 0.0% | 0.0% | 0.0% | 0.0% | 0.0 | 0.0% |
| *age-1::mNG::AID; mNG::AID::mpk-1; Phlh-8>TIR-1::F2A::2x mT2::PH* | KNAA | 64 | 0.0% | 0.0% | 0.0% | 0.0% | 90.6% | 9.4% | 0.0% | 0.0% | 0.0% | 0.0% | 0.0% | 0.0% | 0.1 | 0.0% |
| | Control | 60 | 0.0% | 0.0% | 0.0% | 0.0% | 100.0% | 0.0% | 0.0% | 0.0% | 0.0% | 0.0% | 0.0% | 0.0% | 0.0 | 0.0% |
| *PLCγ/plc-3(tm1340)* | None | 60 | 0.0% | 0.0% | 0.0% | 0.0% | 100.0% | 0.0% | 0.0% | 0.0% | 0.0% | 0.0% | 0.0% | 0.0% | 0.0 | 0.0% |
| *mScarlet-I::AID::ced-10; mNG::AID::mig-2; Phlh-8>TIR-1::F2A::2x mT2::PH***** | KNAA - Timed* | 76 | 0.0% | 0.0% | 0.0% | 0.0% | 88.2% | 10.5% | 1.3% | 0.0% | 0.0% | 0.0% | 0.0% | 0.0% | 0.2 | 0.0% |
| | Control | 66 | 0.0% | 0.0% | 0.0% | 0.0% | 100.0% | 0.0% | 0.0% | 0.0% | 0.0% | 0.0% | 0.0% | 0.0% | 0.0 | 0.0% |
| *arx-2::mNG::AID; Phlh-8>TIR-1::F2A::2x mT2::PH****** | KNAA - Timed* | 70 | 0.0% | 0.0% | 0.0% | 0.0% | 100.0% | 0.0% | 0.0% | 0.0% | 0.0% | 0.0% | 0.0% | 0.0% | 0.0 | 0.0% |
| | Control | 62 | 0.0% | 0.0% | 0.0% | 0.0% | 100.0% | 0.0% | 0.0% | 0.0% | 0.0% | 0.0% | 0.0% | 0.0% | 0.0 | 0.0% |
| *arx-2::mNG::AID; Prpl-28>FLP-ON>TIR1(F79G)::T2A::2x mKate2::DHB; Phlh-8>flp-D5* | 5-Ph-IAA - Timed* | 60 | 0.0% | 0.0% | 0.0% | 0.0% | 100.0% | 0.0% | 0.0% | 0.0% | 0.0% | 0.0% | 0.0% | 0.0% | 0.0 | 0.0% |
| | Control | 62 | 0.0% | 0.0% | 0.0% | 0.0% | 100.0% | 0.0% | 0.0% | 0.0% | 0.0% | 0.0% | 0.0% | 0.0% | 0.0 | 0.0% |
| *abl-1::mNG::AID; Phlh-8>TIR-1::F2A::2x mT2::PH* | KNAA | 66 | 0.0% | 0.0% | 0.0% | 0.0% | 100.0% | 0.0% | 0.0% | 0.0% | 0.0% | 0.0% | 0.0% | 0.0% | 0.0 | 0.0% |
| | Control | 60 | 0.0% | 0.0% | 0.0% | 0.0% | 100.0% | 0.0% | 0.0% | 0.0% | 0.0% | 0.0% | 0.0% | 0.0% | 0.0 | 0.0% |
| *abl-1(ok171)* | None | 66 | 0.0% | 0.0% | 0.0% | 0.0% | 100.0% | 0.0% | 0.0% | 0.0% | 0.0% | 0.0% | 0.0% | 0.0% | 0.0 | 0.0% |
| *mNG::AID::let-502; Phlh-8>TIR-1::F2A::2x mT2::PH* | KNAA | 64 | 0.0% | 0.0% | 0.0% | 0.0% | 100.0% | 0.0% | 0.0% | 0.0% | 0.0% | 0.0% | 0.0% | 0.0% | 0.0 | 0.0% |
| | Control | 60 | 0.0% | 0.0% | 0.0% | 0.0% | 100.0% | 0.0% | 0.0% | 0.0% | 0.0% | 0.0% | 0.0% | 0.0% | 0.0 | 0.0% |
| *wve-1::mNG::AID; Phlh-8>TIR-1::F2A::2x mT2::PH******* | KNAA - Timed* | 72 | 0.0% | 0.0% | 0.0% | 0.0% | 100.0% | 0.0% | 0.0% | 0.0% | 0.0% | 0.0% | 0.0% | 0.0% | 0.0 | 0.0% |
| | Control | 60 | 0.0% | 0.0% | 0.0% | 0.0% | 100.0% | 0.0% | 0.0% | 0.0% | 0.0% | 0.0% | 0.0% | 0.0% | 0.0 | 0.0% |
| *wsp-1::mScarlet-I::AID; Phlh-8>TIR-1::F2A::2x mT2::PH* | KNAA | 64 | 0.0% | 0.0% | 0.0% | 0.0% | 100.0% | 0.0% | 0.0% | 0.0% | 0.0% | 0.0% | 0.0% | 0.0% | 0.0 | 0.0% |
| | Control | 60 | 0.0% | 0.0% | 0.0% | 0.0% | 100.0% | 0.0% | 0.0% | 0.0% | 0.0% | 0.0% | 0.0% | 0.0% | 0.0 | 0.0% |
| *wve-1::mNG::AID; wsp-1::mScarlet-I::AID; Phlh-8>TIR-1::F2A::2x mT2::PH******** | KNAA - Timed* | 64 | 0.0% | 0.0% | 0.0% | 0.0% | 100.0% | 0.0% | 0.0% | 0.0% | 0.0% | 0.0% | 0.0% | 0.0% | 0.0 | 0.0% |
| | Control | 60 | 0.0% | 0.0% | 0.0% | 0.0% | 100.0% | 0.0% | 0.0% | 0.0% | 0.0% | 0.0% | 0.0% | 0.0% | 0.0 | 0.0% |
| *unc-34::mNG::AID; Phlh-8>TIR-1::F2A::2x mT2::PH* | KNAA | 76 | 0.0% | 0.0% | 0.0% | 0.0% | 100.0% | 0.0% | 0.0% | 0.0% | 0.0% | 0.0% | 0.0% | 0.0% | 0.0 | 0.0% |
| | Control | 70 | 0.0% | 0.0% | 0.0% | 0.0% | 100.0% | 0.0% | 0.0% | 0.0% | 0.0% | 0.0% | 0.0% | 0.0% | 0.0 | 0.0% |
| *Phlh-8>SOScat::2x mTurquoise::PH* | None | 102 | 0.0% | 0.0% | 0.0% | 0.0% | 56.9% | 17.6% | 17.6% | 7.8% | 0.0% | 0.0% | 0.0% | 0.0% | 0.8 | 15.7% |
| *Pegl-20>FGF/egl-17; egl-17((ljf14[deletion + mNG^3xFlag])* | None | 70 | 0.0% | 0.0% | 0.0% | 0.0% | 0.0% | 0.0% | 0.0% | 0.0% | 17.1% | 70.0% | 12.9% | 0.0% | 5.0 | 4.3% |
| *Phlh-8>SOScat::2x mTurquoise::PH; mNG::AID::mpk-1; Phlh-8>TIR-1::F2A::2x mT2::PH* | KNAA | 92 | 0.0% | 0.0% | 0.0% | 0.0% | 28.3% | 38.0% | 29.0% | 4.3% | 0.0% | 0.0% | 0.0% | 0.0% | 1.1 | 1.1% |
| | Control | 138 | 0.0% | 0.0% | 0.0% | 0.0% | 59.4% | 13.8% | 21.7% | 4.3% | 0.7% | 0.0% | 0.0% | 0.0% | 0.7 | 21.7% |
| *Phlh-8>SOScat::2x mTurquoise::PH; Pegl-20>FGF/egl-17; egl-17((ljf14[deletion + mNG^3xFlag])* | None | 78 | 0.0% | 0.0% | 0.0% | 0.0% | 15.4% | 25.6% | 35.9% | 21.8% | 1.3% | 0.0% | 0.0% | 0.0% | 1.7 | 37.2% |
| *Phlh-8>sem-5::mNG::PH********* | None | 60 | 0.0% | 0.0% | 0.0% | 0.0% | 85.0% | 6.7% | 8.3% | 0.0% | 0.0% | 0.0% | 0.0% | 0.0% | 0.2 | 8.3% |
| *Ras/let-60(G13E)* | None | 68 | 0.0% | 0.0% | 0.0% | 0.0% | 95.6% | 4.4% | 0.0% | 0.0% | 0.0% | 0.0% | 0.0% | 0.0% | 0.0 | 1.5% |
| *Ras/let-60(G13E G75D)* | None | 72 | 0.0% | 0.0% | 0.0% | 0.0% | 34.7% | 33.3% | 23.6% | 8.3% | 0.0% | 0.0% | 0.0% | 0.0% | 1.1 | 11.1% |
| *Phlh-8>let-60(G13E)* | None | 70 | 0.0% | 0.0% | 0.0% | 0.0% | 100.0% | 0.0% | 0.0% | 0.0% | 0.0% | 0.0% | 0.0% | 0.0% | 0.0 | 0.0% |
| *Phlh-8>let-60(G13E); egl-15(n1458)* | None | 66 | 0.0% | 0.0% | 0.0% | 0.0% | 40.9% | 30.3% | 24.2% | 4.5% | 0.0% | 0.0% | 0.0% | 0.0% | 0.9 | 28.8% |
| *Phlh-8>let-60(G13E); egl-15(n1458); mNG::AID:::mpk-1; Phlh-8>TIR-1::F2A::2x mT2::PH* | KNAA | 108 | 0.0% | 0.0% | 0.0% | 0.0% | 15.7% | 46.3% | 34.3% | 1.9% | 0.9% | 0.0% | 0.0% | 0.0% | 1.2 | 3.7% |
| | Control | 62 | 0.0% | 0.0% | 0.0% | 0.0% | 41.9% | 38.7% | 17.7% | 1.6% | 0.0% | 0.0% | 0.0% | 0.0% | 0.8 | 19.4% |
| *Phlh-8>let-60(G12V)* | None | 68 | 0.0% | 0.0% | 0.0% | 0.0% | 95.6% | 4.4% | 0.0% | 0.0% | 0.0% | 0.0% | 0.0% | 0.0% | 0.0 | 2.9% |
| *Phlh-8>let-60(G12V); mNG::AID:::mpk-1; Phlh-8>TIR-1::F2A::2x mT2::PH* | KNAA | 124 | 1.6% | 3.2% | 8.9% | 22.6% | 52.4% | 8.9% | 2.4% | 0.0% | 0.0% | 0.0% | 0.0% | 0.0% | -0.1 | 8.9% |
| | Control | 118 | 0.0% | 1.7% | 0.8% | 6.8% | 89.0% | 1.7% | 0.0% | 0.0% | 0.0% | 0.0% | 0.0% | 0.0% | -0.4 | 11.9% |

*KNAA-Staged = Adult worms bleached 2 days prior to stage worms and treated with auxin at the moment the SMs were born to alleviate observed problems in m lineage defects that occurred prior to SM migration (see materials and methods)

**Superficially wild-type F1 progeny of *age-1(m333)/mnC1 [dpy-10(e128); unc-52(e444)]* animals were scored for SM migration defects. 33% of these animals are expected to be *age-1(m333)* homozygotes. SM positioning was not scored in 5 additional worms due to M lineage differentiation defects resulting in an abnormal SM number (see Supplemental Figure 2)

***When on continuous auxin, SM's always failed to properly form (30 worms scored). Instead, the m lineage usually divded less times, and cells were very small or completely absent by the late L3 stage

****When on continuous auxin, (30 worms scored), only 4 looked 'normal', 2 had a dorsal/ventral division instead of a anterior-posterior division for the first division of an SM, 9 appeared to have both SMs on the same side of the worm (7 left, 2 right), 4 had too many SMs potentially due to differnetiation issues in the m lineage, and 11 appeared to be missing a cell on one side, potentially due to differentiation issues.

*****When on continuous auxin, (38 worms scored), 20 worms appeared 'normal', 10 had a missing cell on one side likely due to differentiation issues in the m lineage, 4 had both cells on the same side, and 4 had one cell on the dorsal side of the worm instead of ventral.

******When on continuous auxin, (33 worms scored), 29 appeared WT. 2 worms had both SMs on the same side, implying WAVE depletion earlier in te m lineage could impact cell division. In addition, 1 worm had an SM missing, and another had 1 SM that was displaced (left side, scored as a 2) and was dorsal

*******When on continuous auxin, (38 worms scored), 14 appeared WT, 1 had both SMs missing, 2 had WT positioning but one cell was dorsal, 10 had both SMs on the right side of the worm, with 5 of those showing 1 SM being displaced, indicating earlier issues with division/differentiation during m lineage development, 9 had the left SM missing entirely with 2 of thsoe having a displaced right SM, and 2 worms had a left or right cell being displaced around position 2.

********Of cells that are displaced, 55.6% are dorsal

| Supplemental Table 2. An intragenic revertant of *let-60(n1046gf)* with strong SM migration defects | | | | | |
|---|---|---|---|---|---|
| **Genotype** | **Mutation** | **% lethal (n)** | **% Egl (n)** | **% Vul (n)** | **% SM mig (n)** |
| *let-60(n1046ku75)* | *let-60(G13E G75D)* | 2 (156) | 82 (152) | 12 (25) | 93 (72) |

Vul = underinduced (fewer than 22 vulval nuclei). SM mig = SM nearest to a VPC other than P6.p.
*n1046* = G13E
*ku75 = G75D*

### Supplemental File 1: Genome editing and molecular cloning information

| Genomic edits and insertions generated by co-CRISPR | | |
|---|---|---|
| **Target/purpose** | **crRNA (5' – 3')** | **ssODN/dsDNA repair template (5' – 3')** |
| *dpy-10* for co-CRISPR | GCUACCAUAGGCACCACGAG | CACTTGAACTTCAATACGGCAAGATGAGAATGACTGGAAACCGTACCGCAT<br>GCGGTGCCTATGGTAGCGGAGCTTCACATGGCTTCAGACCAACAGCCTAT |
| *egl-15(5a)* loss of function | AACAGAAAAUUGUUCGAGCC | atgaataatgaacagaaaattgttcgagccaggcacAGCATCGGGAGCCTC<br>CCTAGGGATTACAAGGATGACGATGACAAGAGAAGCATCGGGAGCCTCAGG<br>AGCATTctgagattaagacagatcaatcatgctttgggtagg<br>Purple: insertion causing loss of function |
| *let-60(G13E)* | AAGCUUGUGGUAGUUGGAGA | ATGACGGAGTACAAGCTTGTGGTAGTTGGAGATGGAGAAGTTGGTAAATCA<br>GCACTCACCATTCAACTCATCC |
| *let-60(G75D)* | CGAUGCGUGAUCAGUACAUG | AATATTCGGCGATGCGTGATCAGTACATGAGGACAGACGAAGGATTTCTGT<br>TGGTTTTCGCCGTCAACGAGGC |
| Codon-optimized *sqt-1* from SEC * | CCAUCUACUCGGAGGUUGA | TTATGGTATGCTTCATGACCATGTCAACCATCTACTAGATGAcTAGTCGGA<br>GGTTGATGGATTCAGAGAGAAGCTTGATAC |
| *let-60* to generate *let-60(re302[d10:let-60])* | AAUGACGGAGUACAAGCUUG | catttttccatattcaactatgcgtctttttttcagaaaagggtaATGACGG<br>AGTACAAGGGAAACCGTA**CCGCTCGTGGTGCCTATGGTAG**CGGAGCTTCAA<br>TGACGGAGTACAAGCTTGTAGTAGTTGGAGATGGAGGAGTTGGTAAATCAG<br>CACTCACCATTCAACTC<br>**Bold:** *dpy-10* guide sequence + PAM (antisense) |
| *dpy-10* to insert *AID* into *let-60(re302[d10:let-60])* | GCUACCAUAGGCACCACGAG | taATGACGGAGTACAAGGGAAACCGTACCGCTCagGGAATGCCAAAGGACC<br>CAGCTAAGCCACCAGCTAAGGCTCAAGTTGTTGGATGGCCACCAGTTCGTT<br>CTTACCGTAAGAACGTTATGGTTTCTTGCCAAAAGTCTTCTGGAGGACCAG<br>AGGCTGCTGCCTTCGTCAAGGGAGCATCGGGAGCCTCAGGAGCATCGcGTG<br>GTGCCTATGGTAGCGGAGCTTCAATGACGGAG<br>Cyan highlight: AID |
| *dpy-10* to insert *mScarlet-I* into *let-60(re302[d10:let-60])* | GCUACCAUAGGCACCACGAG | TGACGGAGTACAAGGGAAACCGTACCGCTCGTGGTGCCTATGGTAGCATGG<br>TCTCCAAAGGAGAAGCTGTGATCAAGGAGTTCATGCGCTTCAAGGTTCACA<br>TGGAAGGAAGCATGAATGGTCACGAGTTCGAAATCGAAGGAGAAGGGGAGG<br>GCCGCCCGTACGAGGGAACTCAGACCGCCAAGCTGAAGGTCACCAAGGGAG<br>gtaagtttgtgataatccaatttcaattcgcaatggtcatcgttttttcag<br>GACCACTTCCATTCTCATGGGATATTCTCTCCCCACAATTCATGTACGGCT<br>CCCGTGCTTTCATCAAACACCCAGCCGACATTCCAGATTACTACAAGCAAT<br>CTTTCCCAGAAGGCTTCAAGTGGGAGCGTGTGATGAACTTCGAGGATGGTG<br>GAGCAGTTACAGTAACTCAGGACACTTCTCTCGAGgtaagtttttactccg<br>cttttaacaatggttgtttgacatcatttttcagGATGGCACTTTGATCT<br>ACAAGGTCAAGCTCCGTGGTACCAATTTCCCACCAGATGGACCAGTTATGC<br>AGAAGAAGACGATGGGATGGGAGGCTTCCACCGAACGATTGTACCCAGAAG<br>ATGGAGTTCTCAAGGGAGACATCAAAATGGCTCTTCGCCTCAAGGACGGAG<br>gtaagtttgatgaaacggtttcgtcttatatacactaatggtacttttcag<br>GACGTTACCTGGCGGATTTCAAGACCACATACAAGGCAAAGAAGCCAGTTC<br>AAATGCCAGGAGCATATAACGTTGACCGCAAGCTTGATATTACTTCCCATA<br>ATGAAGACTACACAGTTGTAGAACAATACGAACGATCCGAGGGACGTCATT<br>CGACCGGAGGAATGGACGAGCTGTACGGAGCCGGATCTTATCCATACGATG<br>TCCCAGATTACGCTTACCCATATGACGTTCCAGACTATGCCGGTGGCGGTG<br>GATCGGGAGGAGGAGGTTCGGGTGGCGGAGGCAGTGGAGGTGGCGGAAGTG<br>GCGGCGGTGGTTCAGGAGGAGGCGGATCCGGAGCTTCAATGACGGAGTACA<br>AGCTTGTAGTAGT<br>Red: mScarlet-1 (GLO)<br>Blue: 2xHA<br>Grey highlight: 30X linker |

* This edit inserts a stop codon at the 5' end of the coding optimized *sqt-1(d)* gene in the self-excising cassette repair template used to generate endogenous protein knock-ins by the SEC method (Dickinson et al., *Genetics*, 2015).

| Genomic insertions generated by SEC | | |
|---|---|---|
| **Gene/Location** | **Insertion Site** | **Guide target sequence (5' – 3')** |
| *abl-1* | FDQIMRLVDR^STOP | GATCCACCAGCCTCATGATT |
| *age-1* | AFNGSWSTKT^NWLFHAVKHY | GTGGAAGAGCCAATTCGTTT |
| *akt-1* | M^SMTSLSTKSR | TGTCATCGACATTCTTTCAC |
| *arx-2* | RCMAKLGIKA^STOP | GGGTGGTATTGCAAGATGTA |
| *ced-10* | VGKTCLLISY^TTNAFPGEYI | GGGAAATGCGTTTGTGGTGT |
| *egl-15* | APVNLPSEPQ^HTICDDYESN | TCGTCGCAAATTGTGTGTTG |
| *ksr-1* | M^MMQTQVASRA | TCCCGCACGTGATGCAACTT |
| *let-363* | MLQ^QHGISFQMNA | GAAAACTAATTCCGTGTTGT |
| *let-502* | M^EQDELRDQLV | ACGCAGCTCATCCTGCTCCA |
| *mig-2* | M^SSPSRQIKCV | ACACATTTGATCTGCCTCGA |
| *mpk-1* | M^ADGEAVISTV | AATGAGCATATCACTGCACT |
| *sem-5* | LNNRRGIFPS^NYVCPYNSNK | TTCTTCTTGCAGATGGCCGA |
| *sos-1* | M^SLHSASSDTV | AAGGGCATACGTAGTTGGAT |
| *wsp-1* | DEDDKNEWSD^STOP | ATGAAGATGACAAAAATGAA |
| *wve-1* | EGADDDEWDD^STOP | GAGACGAATGGATCAAGTTA |
| Chr I transgenes | Chr I:2,851,088 | GAAATCGCCGACTTGCGAGG |
| Chr IV transgenes | Chr IV:4,237,723 | ACTGTTGGATGCCTGTGTAG |

### Strategy used to generate *let-60* N-terminal knock-ins

#### Generation of *let-60* landing pad allele *let-60(re302[d10::let-60)* with *dpy-10* gRNA site

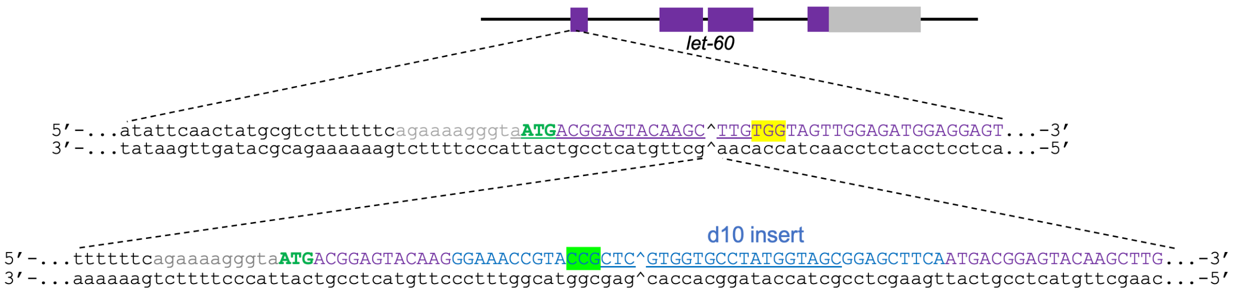


#### Insertion of mScarlet-I::2x HA 30x linker into *let-60(re302[d10::let-60)*

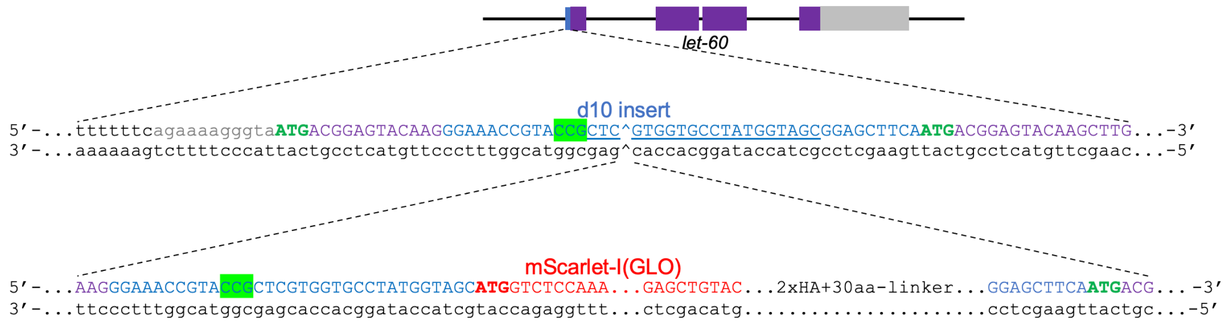


<u>2xHA:: 30aa linker sequence:</u>

TATCCATACGATGTCCCAGATTACGCTTACCCATATGACGTTCCAGACTATGCCGGTGGCGGTGGATCGGGAGGAGG
AGGTTCGGGTGGCGGAGGCAGTGGAGGTGGCGGAAGTGGCGGCGGTGGTTCAGGAGGAGGCGGATCC

#### Insertion of AID into *let-60(re302[d10::let-60)*

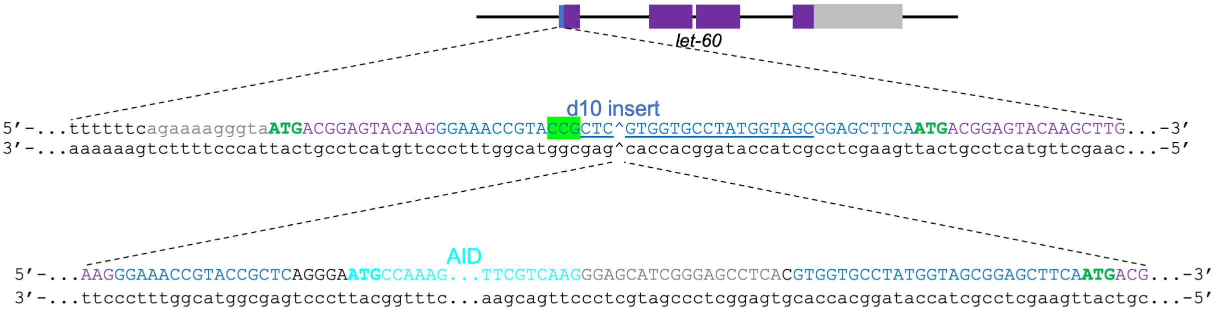


Purple: *let-60* coding sequence
Blue: *dpy-10* coding sequence
**Bold:** Initiator methionine codon
<u>Underlined</u>: guide RNA target sequence
^: Cas9 cut site
+strand PAM
-strand PAM

### ***Promoters and homology arms for single-copy transgene construction***

#### ***hlh-8* promoter:**

aaagctttcaaaatcccaaacatagcatcttacttactcgcctttgctcccaaagatttaccattttcttgtttttg
tttgttaatttcctctcccacccattttcaaccatgtcatttcacacattaaactcagtatgcatcgatttaattgt
tccaggctaattgggtttctgtcattgttaatagacacaatttttactgagaaatccttttcatatgtgtgaaattt
gttctttaaacgtgcatttcctatttatcgctctctagatttgatgccagttcgtgtagtttttcccaaccgttga
ctcatcgataccgcaatccatttcacccagtatttgtaccattgccgtgaattcacagtattacatggtcgtcccct
tttccacaaaatctctttcagactccctccaccctgatcccgttttaattgtttcttgctgatttccctttcaaccg
atcttcgtttttttttcaatagggtatatttgaaatgtcggaatgtcgattcagcagtaaatttgatatgacactct
gactttttaaatgtaaatctgattgtttatcagaattcaaccgggcccactgattattcattaattatgcgacaag
aagaataacgttatagtttttctttcttaagcatttttcatcatcaacatttttgtttagatttaattttgttgct
aataaaaaactacacaaaacgccaagcacatttatttgacttttttttttgagtaatgcagtaagtaggcaatttgt
tgtttttataattgattagttttaaggcctacttgcaaaactgagaaaattcttatcaagaattcaaaaccatttac
gattatcggattttaatatggacaatgttactcattttggataaatcaaaagagtttttggattttaaaattaata
actgtttattactgattaacaaaagattacggtaaattggagttacaaaatggacaagttttatttatgaatcaaga
catattttgataaaaaatatgggtaccccgccaatttaaaaataaatttttaacttaaaaaaaaacgtgtgaaat
gctttcatattttcaggcttacttagaaaaacaccattcctaagtctaacgaggaaaatgggaaacatgggaaatat
taccgaaacctgggaaatatttttattgattccaaattttcccttgattccaaatatcgatgtgaaaaaaaattaa
aacaaaaattactgattttatttaagcttgaaatcacaaattccatttttatgtcatacttcagatttaaacaaatt
tattttatgtgtgtttttttaatggtatgtcctaacgattttcctaattgacaactattatagattgaaaacacaga
atgccaagtaacgtaagaaattttttttcgaataccagacataatttcaataaaagaaaaatctttaaaaaaagtt
taactttatactacaataattttggttttatgaaggaatctgtattgcggcacatcatgtatttgctcaaggtttc
gttggttgacatatttttgtgatccgtggaatgagcttgcatagagtttagtcataattgggtttattttacataca
gtttgaagtatgttttgtctgttattcgaaatttttagttagtttggtaagttgaaaagaatttacaaaatttaac
taaacgaggacgctcatctcgattcttataattccataataattctacggtaaaagtcaagttatgcctcaaaacat
gtaatacataattaccaaaactacttaattaccgcatttccttagtattttaatagtggtgtagtcgaattttttt
attgctttattagactcaaaattgtctgcaaacacccaatttcataatgaaacttattgaaaacaatacactttgaa
acaatttagtatttcaggaataaggtcactggaacttcgaaaatgataatttgaataaccgttaataaaatattcaa
accaatttgcataatcttgattttgtatcatgatggtatagaatgggacttttgaaagatgtgaagtttcaattcag
taagtacaactttaaaattgggctgcaagatttcttcttaaattttcaacagtttcaaaacactgaaatcatgctga
actatggttagagactcgacaactttcctaaattttggagcagtttcaatggttttcgaatgtatactacatttgta
ccattttatgtgttagtgtaggttgctttgcttctaccttcactctcaaattctttttccgcggtaatttttcaac
tatcgaaacaaagaagaagtcatgaataactgtccgcatagagttctcacccgtcccctttcttttcaccgagcagt
cacgagagaaagaagaagaagcggaccgctgcagagattcttcgtagcggagccccgcccatcctctgttacgcata
tgtttggatctcatttcttccactcattaacgttccaggtatgttttttttgttgatttttaatgtgttacttttc
atatgtttattacttgaactgaccgatttcaaatatgattttcacag

#### ***egl-20*** $^{(-1261 – 610)}$ **enhancer:**

*Underlined sequence is the *pes-10* minimal promoter.

gttggatgttttcttataaaattatcttggaagtttttttaaaaattaaatactctgttacatttttttgcaatgttt
ggaattgaaatatagaacaatataaaacttctagagagaaaaattacttccagaatattgataaatgataaagaata
ttgataacctgaggatttgccactttttgacggcagcattaaaaattgcaattggatactcaagaagataaagtgat
caattatggagaaaataaaatactgtgtaaataaatgatttttttaaacatttttttccgaaatataggtgttccat
ttgtcacatgataattatgaatattcttgaatttctatgttttgaaaaattgaagctaaaaaataagtgtcaaatat
tgttaaagtcccaaaaatcttggaatttttcgtttcgtattgaccattcacttcttcccatccttttttcaccatttc
aacgtcttcctcttctatttgcataccgacaacgtcttcttctcttccattttctgaatagatccacgggccataaa
gcatgaattggaaaggatggagaagatgacaagataccacatccagcaccatcaattcgaccttcgaattttctaaa
cacattgacattcctcttattcatcattcttcatccctgcaggatcgatttttttgcaaattacgagcgttgtagggg
gcggagcgataggtcctataggttttggtatatcatcattcattcattcattggtacattcatttacccaccttcct
ctttctgagcttctctggagttctgtgcttccttttttcccttatctttatactgtaatttttaactttcaggcattg
attggatcattggccaaaggacccaaaggtatgtttcgaatgatactaacataacatagaacattttcaggaggacc
cttgcttggaggagctcagaaaa

### Chr I transgene insertion site (Chr I: 2,851,088)

**Guide RNA:** GAAATCGCCGACTTGCGAGG

**5' homology arm: Chr I:2,851,057-2,852,649 (- strand):**
caagtggggatcaggaagaagttgataaagtgattcaaaaactcgaggagctcaacacagcgatgaacccaatttg
agagcaacactgacgaagaactatgacgcgagggtaggcactaccttgaagctatccacctgcctgcctgccgccta
tttccatgtgcagtaggaacgtgtgtaggtataggcaggcattatcgcgtctcctttcgaactacctagtacctagt
aggtagtttgaagggaggcgcgataatgcctgcctatgtctacacatgtgcctactgcgcatggaaataggcggtag
tcaggcaggtagatagcctcataaaattcaatacttccttttcagaaagaatgctggaatgtattcaaagaggagtc
actcttcccattgctcagagaatatgaagagttgctcggcgagcaaaagttcatgattggagatcgatattcatggg
ctgacatcgctgtcgtcgagtttctaactcggtgtcagcagtgctacgatagcttctatctggctcatttcgagagg
ctccgaactttttgccagacttttgagtctcttccgcacattcgtccctacattcagggacgtgttgattcatttat
ctaatttttaaaatggatatttttcgtttttatttgtttgttttgctgataattcaataaagttacgcaacgaat
tttagacatttatgaaaaccgaaaatcgacaaaaatatttaagaaaaatagcagcaaaattcaatattttatcaaa
ttttgtttatggtaatgagaaagaaaacatacaatatttttctaagaagaagaaaaatgaccagaattttaagattt
ttctcgaatatacgaaaaattgataaactaaaattttcgatttttccatttattttgaaaaaaaaactcttgacggt
tttcaccatgcgaagaggtgagaaatattctgaaagttatgcattttgaaaaaattgcaaaaaaatacgttaaaatt
ttttgatctattgtttttttttgaaaaccggaatttaaaatggaatttttttttcaaaaattaaaattaaaatttaaa
tatttctagtgtttggatagtttttcaaaagctagatttaacccgcgagatgtcgaccgctagtgtagcttacagat
ggctgaagacttcgcaaaaatcttatcttagaaacccgcgaggtgtcggccgctaaaactttgtaaacctgccaaca
aaatgcttcgcattgtctgaacagatgaaaccttgaaacaaaacatatacaattgaaatcccgtaaagtgtcgggcg
caagaactgcgattagttgtataattggactagctcaaatttgaccaaaatctcatttttagacctctaagatgtcg
gccactataactttattcaacttgaaactgcaaaaaaacttatcaactgtatagattcaagtattatgtcaagtatt
ggttttagagctacagtacactctgataacaaaatctgcagaaatacctccctgtcaattcccaaaatacctcccct
catctcaattatcccgaaaatcggttagtcattgccatcagaaaatcgaatta

**5' homology arm: Chr I: 2,849,538-2,851,008 (- strand):**
gcaagtcggcgatttctttgaagttttgtcaaaagaagagacaaggacttgatagatacgctcatcagagtgagatt
cattgcaaatagtgtgtaaatcgtggagagacgagagaaaaggtgtgagaagcttagaagataaagaataggaagaa
gaacatatctcatttcctattttctctaatttttcgttcatttttctagacattacccatttgatggtttttcaata
tggttatcagggtactcattttggtagttttcaaagcaatttggagtttttttttaaattgtcaactacaaaaacgc
agcttaaaggtccaaatatcatagtataaaaattctttaaaaaattccaaattttatatgatttttttgaaaattgga
agaatgttagtttttgccattttttggaagtttttctccggattttatggtacttatatttcaaatataaattttt
gtaggaattcgagtatgcgagagagctatagtttactaaaaaaaagtggcaaaaactcagtaattgccactttttc
aaaaatgttttgaaaaattttactatgatatttggccattttggcaccatatgtacgttacttttacgaaacactc
taatttttgaaatttgagtttcatgtttaatcttttgtataaaatttttatggtccttttgtgacaggataaaaaat
tgcaaaaaaatagaaaactttgtttcgtgaaaacacctagtttaaaaattgtgggccacttagcgattctagtcaaa
aatcagacaaatgtcagctgccgtaacttttccggatcagggttatcaaattttaattttaattgtaaatgtagaa
aaatgatcggcaacatttgctaattgaaaatattttgaaaattgctatagtacatattaatctagtattaatatag
tattaatgttataatatcagttatgtgtttgaataactcaagcgtaagtttccgtaaaatatctttaaattttggta
attttttacggtttggtataaatttttgcatttttgattttctacactttttttcgatttcgcattttccgaattt
ctctgaaataaatcgaacaatcggaaaatttcgagggcacccttagccatgcagctcaggaatttgactttatgtt
cccattcaaatttttttaaaattaaataaaatgctaattgtaagataaattacaaaaattattaaatctcatcatcc
ttatgatgatagtgcagtgcatgaatatattttccaaataggatatgattcgtcgagtagcttcacgtaatccgtt
ttcttcaatcaccatttcgaactgcaattttttttttggatttttgtaacaatattgtcataaattgtgattttgat
attttgcgtagagcatcagaacctttctgctttgaattttaatgggattaaaaattataaaaaacttgcaccctaat
taaagcgc

### Chr IV transgene insertion site (Chr IV:4,237,723)

**Guide RNA:** ACTGTTGGATGCCTGTGTAG

**5' homology arm: Chr IV:4,236,614 – 4,237,722 (+ strand)**
aggaatcttcttcgcggacggaactcgccgctaagcggatcgcctttaatttattctttgccttatgcgttgctcaa
aataaaacattatttcatccaaaataatttttttgaggttttagatgaatttgatagtttagcttgtcactttctcct

gtgcgaagaaacttgctaagcaggtaagcagcatccttatcaagtttcgaccttatctctcagtagtcgcatacgtc
gtatccctcatctgctttgttttagccacttcaagggaattcagatcactttttctcatgtgttttcaaatttactt
ttgtggagtgaccttaattgaccctctcgtcataagtggacggtgggtagttacttaacaaatttcgagtacaggat
agtgtcaatggcatgagacatgtagtagccaagcaattatggcgaataagtttcaaaaaccaacattctgtcaaata
caaagacctgcaacattagagaattcattaaatacatcttcgaggaaatcaaattttcccgtagacgatcccgtaga
cctacttgtacaatatgttctagatatacactccgaacttaaaatatctcgaacatttgtgttgaatatttatttta
attctaggtttttcaaattccgcacgatttttgccatactactgcccaatagtttacatttattgggcctggatctg
taaggcacttttgggagctccctttcttttccttcaaattacccatacagatctataaatcacaatttaatacaaga
ttttattcaagttaaaaacataaaaggggctttgcggtggtccccatttcaccagagaactataaaaaatataaatt
taacatcaaatgttgcaattgattccctctcaaagctgcttgaagaaccctgattctgtcaagcctatgaagattta
aaaaaaattgggaagacccttagttccaaacaagtgtcgttgacccagtaggcgatattctgaaaagtcataaaatg
gggttgctaaaaaattggtcgctaacttacatttagctaggaatgttaattggaataactcataatttacagtaaaa
ttatcaaaaaatagcaaaaatcaatagagga

**3’ homology arm (Chr IV 4,237,729 – 4,238,654: (+ strand)**
cacaggcatccaacagtacgagttctgcgtttttgaattatagaatcaagcatgctccgcgtttgctgtactatttc
tcgcagagcagttttttttctcacactttattattctgtactgcccctccacgctataatttttgcatagagggcc
ataaaggaaaacggtacggtaaaaataatgggtcctttttaccgttgtgtagattgtttctaacgggatgcaaattt
tcactaaaactctggaaaactgatggaattctacgggcagccggcaattttagatatttgaagaatccttcttgatc
agtgccaagctttcccatggaccaaagatgtggatccaagatcaaatgcacaggatctgaactcctttgttccaaaa
acaaggatttcttttctatagtttgacctaaatattatagtagaacatatacatttgatgatttcatgaataattta
ttgttaaaaaataatacatgtgttaaaaaataattctaaaaaattaattggagcgtccttggcggatatggcggatt
ggaaggtcataagtaactcgaactaacctgtaaatataattaatttagaagtaatcattttgaaatgtttctaaatt
tttgtaaacgtaatcaatgatttacttacaaatttttcgaaatattagtagaaatataaaaaaccagtaaaattca
aaagaggccagccacacatgcatcttacaatacgaatgatgcgtttttttgtcgcctcggagaaggaatcaagcaggc
tccgcgttttctgtactatttctcccagagccgtttattttgcataggggggtgggtcataaacggagacgttttggt
ataaaatttgccccttcttccaccgttgaaaaaattggaaaatgaaccgggacaggaaaggggatttcaggataatg
gt
